# Stimulus-Driven Leakage in Naturalistic Neuroimaging

**DOI:** 10.1101/2025.04.01.646146

**Authors:** Seung-Goo Kim

## Abstract

This article elucidates a methodological pitfall of cross-validation for evaluating predictive models applied to naturalistic neuroimaging data—–namely, “stimulus-driven leakage.” While this problem has been well known as “leakage in training examples” in machine learning, it may be difficult to detect in practice due to conventions in neuroscience. Stimulus-driven leakage can occur when predictive modelling is applied to data from a conventional neuroscientific design, characterized by a limited set of stimuli repeated across trials and/or participants. It results in spurious predictive performance due to overfitting to repeated signals, even in the presence of independent noise. Through comprehensive simulations and real-world examples, following a theoretical formulation, the article underscores how such data leakage can occur and how severely it can compromise results and conclusions when combined with widely spread informal reverse inference. The article concludes with practical recommendations for researchers to avoid stimulus-driven leakage in their experimental design and analysis.

## 1 Introduction

### 1.1 Data leakage in predictive modelling

In statistical learning and machine learning, evaluation of a learned model is a critical step since the model is, by design, highly flexible in learning patterns in the data, which simultaneously gives rise to the possibility of overfitting to noise that always exists in the data. Thus, it is important to evaluate the model’s performance on an independent test set that was not used during the training and optimisation processes. Surprisingly often, designing a legitimate evaluation can be non-rivial (Kaufman et al., 2012). The key aspect is to keep the training, validation (i.e., optimisation), and test partitions completely separated. When this fails, data may ‘leak’ into the model from the validation or test set, which must be kept ‘sealed’. The model may then behave over-optimistically, leading to wrong conclusions such as overconfidence in its generalisability and overestimation of feature importance. This is known as ‘data leakage’, and it remains a major challenge in predictive modelling, even in recent machine-learning-based scientific research (Kapoor & Narayanan, 2023), including neuroimaging (Rosenblatt et al., 2024; Verstynen & Kording, 2023).

### 1.2 Data leakage in naturalistic neuroimaging

While the predictive modelling has been heavily used in clinical neuroimaging research (e.g., brain age gap estimation; Seitz-Holland et al., 2024), it remains relatively novel in many cognitive research domains. This paper primarily focuses on a specific field, often called ‘naturalistic neuroimaging’. It refers to the use of complex, real-world stimuli (e.g., movies, music, natural speech) in neuroimaging experiments to investigate brain function in more ecologically valid contexts (Hamilton & Huth, 2020; Nastase et al., 2020; Sonkusare et al., 2019). In particular, this idea has been widely adopted in domains where high-order cognitive and/or affective processes are involved, and a simple contrastive experimental approach can explain only little. For example, comparing brain responses to *music* vs. *non-music* to find neural correlates of “music perception” may be under an overly reductionist assumption (i.e., “music-as-fixed-effect” fallacy; Kim, 2022) that the human brain is governed by simple, interpretable rules that can extrapolate to explain complex behaviours (for more discussion, see Nastase et al., 2020).

In addressing the complexity of the real-world information, two different approaches have been popularised. On one hand, as a model-based approach, a temporal transfer function estimation based on the electrophysiological tradition of the receptive field mapping has been used (Lalor et al., 2009; Theunissen et al., 2000; Wu et al., 2006). On the other hand, as a model-free approach, a intersubject (or intertrial) correlation has been used to quantify the strength of the stimulus-driven signal in the data (Hasson et al., 2004).

A problem arises when the model-based approach is applied to a dataset collected for the model-free approach without due caution. In the model-free approach, the repeated stimuli across trials and/or participants are essential to identify stimulus-driven effects. Because the noise is independent across trials, cross-validation (CV) partitions that include trials with identical stimuli may appear valid—yet the repeated stimulus itself constitutes data leakage. In this paper, I term this specific form of leakage as *Stimulus-driven Leakage* (SDL) since the repeated stimulus is the source of the leakage. Importantly, SDL is not inherent to the model-based approach or to the use of naturalistic stimuli itself, but arises from their careless combination. According to Kaufman et al., 2012, SDL can be understood as a special form of *leakage in training examples*, which can be also seen as *Non-independence between train and test samples* in a recently proposed taxonomy (Kapoor & Narayanan, 2023).

### 1.3 Aims and scope of the current paper

While the true prevalence of the stimulus-driven leakage in published studies has yet to be found, a preliminary investigation suggests that it is not uncommon in the literature of naturalistic neuroscience including fMRI and M/EEG studies. In particular, while the accessibility of the predictive modelling has largely increased thanks to various open-source software packages (e.g., `mTRF Toolbox` [Crosse et al., 2016]; `scikit-learn` [Pedregosa et al., 2011]), some researchers may not be fully aware of the pitfalls of the predictive modelling, despite efforts to educate researchers about the best practices in encoding analysis (Crosse et al., 2021; Dupré la Tour et al., 2025). This overall situation increases the risk of the data leakage, which could lead to contamination of the literature. Therefore, this paper aims to explain the mechanism of the stimulus-driven leakage and to demonstrate how it can occur and how severely it can compromise the results.

The current paper focuses on a particular problem of predictive modelling: a time series prediction with a finite-impulse response model that fits variable neural delays, regularised with a ridge penalty. This form of predictive modelling is widely used in the naturalistic neuroimaging, especially when involving temporally dynamic stimuli (e.g., movies, music, natural speech). Nonetheless, stimulus-driven leakage can occur in any model in which the same stimuli exist in both training and test sets.

For those who are trained in statistical learning, this may seem obvious and intuitive. However, it may not be so clear to classically trained neuroscientists who are unfamiliar with statistical learning methods because of the independence of the noise across trials and subjects. In the Theory section, a formal analysis shows how the intuition stands. In the Simulation section, contributing factors that worsen the stimulus-driven leakage are identified. Using real-world data, the stimulus-driven leakage is demonstrated in the Real-Data section. Finally, implications for future analyses and experiments are discussed and practical recommendations are given in the Discussion section.

## 2 Theory

Consider a linear model:

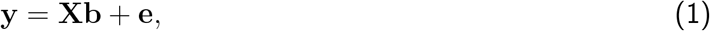

where **y** ∈ ℝ^*T* × 1^ is a response vector over *T* time points of a single response unit (e.g., a channel or a voxel), **X** ∈ ℝ^*T* × *FD*^ is a Toeplitz design matrix constructed from *F* features^1^ with *D* delays, forming a finite impulse response (FIR) model^2^, **b** ∈ ℝ^*FD* × 1^ is an unknown weight vector, **e** ∈ R^*T* × 1^ is a zero-mean, unit-variance Gaussian noise vector **e** ∼ 𝒩_*T*_ (**0, I**_*T*_) where **I**_*T*_ ∈ ℝ^*T* × *T*^ is an identity matrix. For convenience, features (i.e., before Toeplitz construction) and response variables are assumed to be standardized prior to analysis so that their sample means are zero and their variances are one.

A ridge solution (Hoerl & Kennard, 1970) to Equation 1 is given by

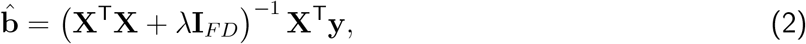

where *λ* > 0 is a regularisation hyperparameter.

For a true model (**X**) with a sufficient signal strength 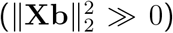, the prediction accuracy (either Pearson correlation or *R*^2^) with a minimal regularisation (*λ* → 0) is expected to be positive whereas that of a null model (i.e., a reasonable random features **U**) with a proper regularisation (*λ* ≫ 0) is expected to be null. This is how the cross-validation (CV) is supposed to works in usual cases. However, when the same stimulus is repeated across CV partitions, the expected prediction accuracy of the null model can be positive.

But when can the same stimulus be repeated across partitions? To illustrate, consider two possible modelling approaches for a dataset in which two stimuli were presented to three subjects (Figure 1). One is subject-specific modelling (Figure 1a; e.g., leave-one-stimulus-out CV) where each subject’s data are partitioned into training, validation, and test sets. The other is stimulus-specific modelling (Figure 1b; e.g., leave-one-subject-out CV). In the stimulus-specific modelling, the same stimulus is repeated across CV partitions, albeit with different noise realizations, since the same stimulus is presented to all subjects.

**Figure 1.**
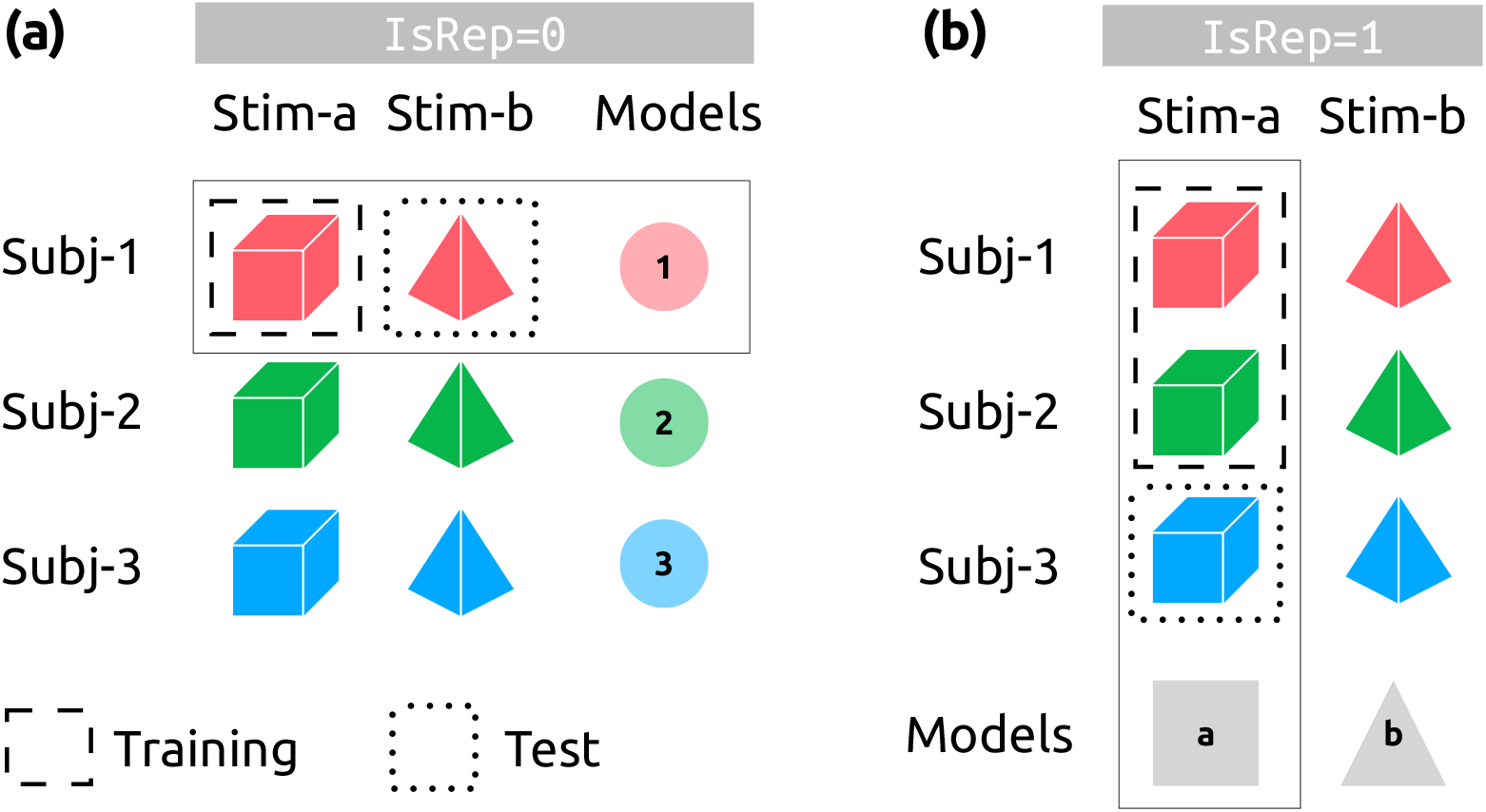
Schema of two cross-validation designs. For simplicity, let us say we have three subjects {1, 2, 3} (depicted in red, green, blue) and two stimuli {*a, b*} (depicted as a cube and a tetrahedron). Note that we assume any repetitions of stimuli within a subject are averaged prior to the cross-validation. **(a)** `IsRep=0`. Subject-specific models (pale coloured circles). Each model (e.g., pale red circle in solid rectangle) is trained with one stimulus (dashed rectangle) and tested on another stimulus (dotted rectangle). **(b)** `IsRep=1`. Stimulus-specific models (gray square and triangle). Each model (e.g., gray square in sold rectangle) is trained with two subjects (dashed rectangle) and tested on the other subject (dotted rectangle). The validation set, which could be in the training set of the outer loop, is not marked for simplicity.

While it may seem obvious to those trained in machine learning that the latter is a flawed CV design, I argue that stimulus-specific modelling may appear legitimate to researchers from other disciplines. This is due to the independence of noise across feature-response pairs, or simply the familiarity of leave-one-*subject*-out CV in other analyses (e.g., multi-voxel pattern analysis [MVPA]; Kriegeskorte et al., 2006). In particular, avoiding the repetition of identical noise in analysis is common practice in neuroimaging, largely thanks to the pedagogical work that popularised the term “double-dipping” (Kriegeskorte et al., 2009). Where double-dipping concerns the repetition of identical noise, SDL concerns the repetition of identical signal. Given that the data is the sum of signal and noise, SDL can be thought of as “inverse double-dipping”.

With that, the mechanism of SDL can be summarized in the following steps:

1. The repeated signal disables the regularisation of a seemingly valid optimisation process with independent noise.
2. As the regularisation hyperparameter approaches zero, the projection matrix becomes positive definite, which would have been null if properly regularised.
3. As a result, the expected prediction accuracy of the null model over random realisations is proportional to a bilinear form involving a non-zero vector and a positive-definite square matrix, which is positive.

Below are the simplified explanations of the above points. More detailed derivation is given in the Supplementary Theory.

### 2.1 Disabling regularisation

To denote partitions in cross-validation, let us use the subscript *i* for the *i*-th partition: **X**_1_ and **y**_1_ are the training set, **X**_2_ and **y**_2_ the validation set to optimise training, and **X**_3_ and **y**_3_ the test set to estimate out-sample prediction performance. When the identical stimulus is used in all sets (**s**_1_ = **s**_2_ = **s**_3_), the expected validation accuracy (i.e., Pearson correlation) of the null model **U** across random noise realisations is proportional to the following bilinear form:

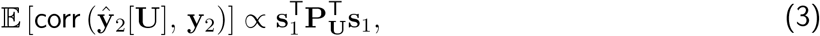

where the projection matrix based on the null model is 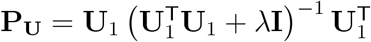.

In this case, the optimal *λ* that maximises the validation can be analytically found using singular vector decomposition (Hastie et al., 2009, Eq. 3.47):

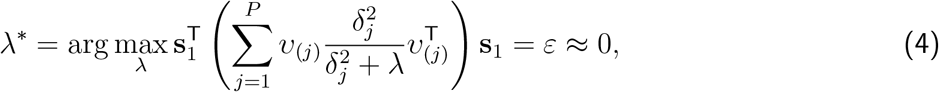

where *υ*_(*j*)_ is a left singular vector of **U**, *δ*_*j*_ is the corresponding singular value, and *ε* is the possible smallest positive value.

### 2.2 Positive definite projection matrix

When unregularised (i.e., *λ*^∗^ ≈ 0), the projection matrix based on the null model can be approximated:

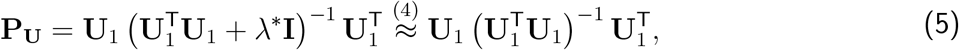

### 2.3 Positive null prediction accuracy

Therefore, the expected test accuracy of the null model is:

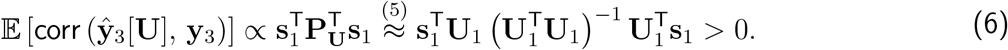

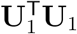 is positive definite due to its symmetry and independence of the columns in **U**_1_. Since the inverse operation preserves the signs of eigenvalues of a square matrix, its inversion 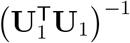 is also positive definite. Thus, the expected prediction accuracy (i.e., a bilinear form involving a non-zero vector and a positive-definite square matrix) is positive.

This may surprise some readers; however, Equation 6 clearly suggests that any random features may seem to predict ‘unseen’ (but leaked in truth) data. Depending on the signal-to-noise ratio (SNR), this may result in significant Type-I (false positive) errors.

## 3 Toy example

For a graphical illustration, a simple case of small-scale simulation (i.e., a toy example) is shown in Figure 2. In this example, 100 time points for 3 response variates were generated with two features and three delays. The SNR was 0 dB. In the first case (Figure 2a), the stimulus was not repeated across CV partitions. Thus, as expected, the prediction accuracies for the null models (pink) were around zero and the optimal ridge penalties were large (i.e., *λ* ≫ 0). However, in the second case (Figure 2b), the stimulus was repeated across CV partitions. Because of the stimulus-driven leakage, the prediction accuracies for the null models were well-above the threshold corresponding to Bonferroni-adjusted one-tailed *P* -value of 0.05, and the optimal ridge penalties were similar to that of the true models. Extended figures with more detailed explanations can be found in the Supplementary Results.

**Figure 2.**
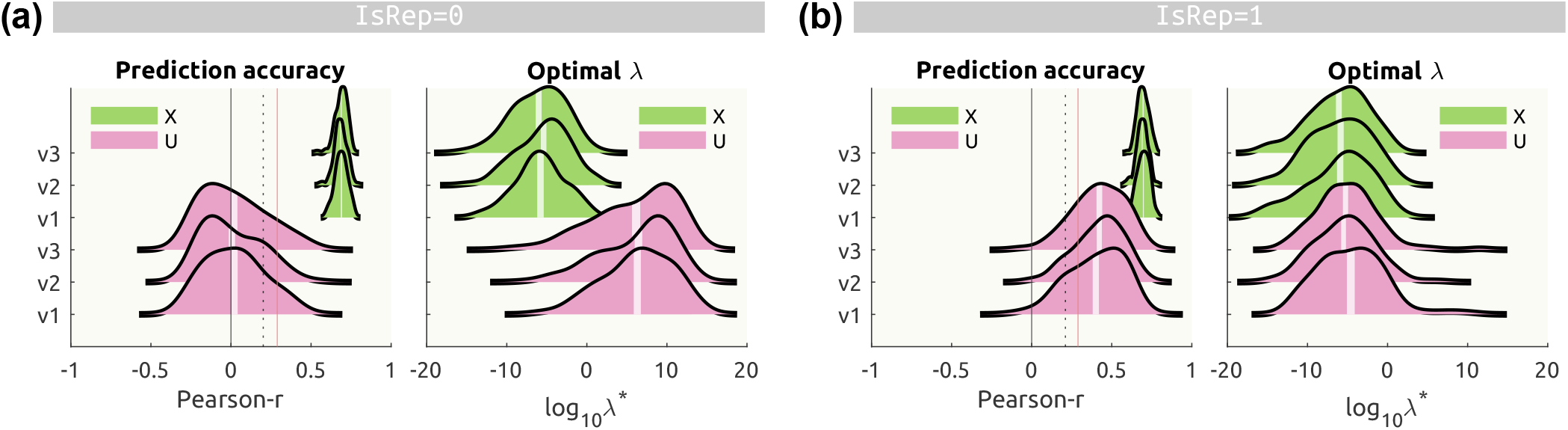
A toy example of stimulus-driven leakage. For each case when (a) the stimulus is not repeated across CV partitions and (b) the stimulus is repeated across CV partitions, prediction accuracies (left) based on the true features (lime green) and null features (pink) over 200 simulations are shown with the optimal ridge penalties (right). White vertical bands of each ridgeline represent the 95% confidence interval of the mean. Vertical lines for Pearson correlation mark non-parametrically estimated thresholds corresponding to one-tailed *P* = 0.05 (gray dashed), and its Bonferroni adjustment (red solid).

The spurious inflation of the prediction accuracy due to the stimulus-driven leakage (i.e., the SDL artefact) is positively proportional to the SNR. Not only that, the SDL artefact is greater when the null model is more flexible (i.e., with more features with less autocorrelation and/or more delays) and or the true features have strong temporal autocorrelation. Please see the Supplementary Results for the extensive simulations quantifying the effect sizes of these factors.

## 4 Real Data

Having explained the theory and shown a toy example, a logical empirical question would be—*Could the SDL happen with the real data in a significant manner* ? In this section, I demonstrate real-data examples of the SDL artefact using open-access data where healthy participants listened to various musical excerpts while measuring neural activity (electroencephalography [EEG] or functional magnetic resonance imaging [fMRI]) or behavioural ratings (Kaneshiro et al., 2020; Sachs et al., 2020). For clarification, none of these studies represents a spurious result. The datasets were used for their classical design (i.e., the same stimuli set for all participants) to demonstrate a fictitious analysis that introduces the SDL.

Based on the well-established encoding of the acoustic energy in the human auditory system, true features were the audio envelopes extracted from the musical stimuli using a cochlear model (Chi et al., 2005). Null features were either (a) the phase-randomised envelope (preserving spectral magnitudes and autocorrelation structures) as the most realistic one, (b) the normal noise, and (c) the uniform noise as the least realistic one. Details of the real data and analysis implementation are provided in the Supplementary Materials.

The magnitude of SDL artefact was estimated as the difference in null prediction accuracies between two CV schemes (i.e., leave-N-stimulus-out for subject-specific modelling vs. leave-n-subject-out for stimulus-specific modelling): 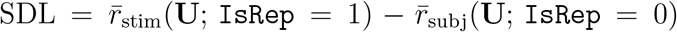 where *r*_stim_ is a prediction accuracy of a stimulus-specific model and *r*_subj_ is that of a subject-specific model. 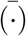 denotes averaging across null models. That is, if there were no false inflation of prediction accuracy due to SDL, the null predictions with and without stimulus repetitions should be equal (i.e., ℋ_0_ : 𝔼(SDL) = 0). Otherwise, the null prediction with stimulus repetitions is expected to be greater than the null prediction without repetitions (ℋ_*A*_ : 𝔼(SDL) *>* 0).

All data analyses were consistently done using an MATLAB package Linearised Encoding Analysis (LEA; https://github.com/seunggookim/lea).

### 4.1 Electroencephalography

Scalp electrical potential data were recorded in 48 healthy participants while listening to Western-style Indian pop music (i.e., Bollywood music; Kaneshiro et al., 2020). Using this EEG dataset, linearised encoding analysis was performed with the audio envelope as a true feature and its phase-randomised signals as null features.

Without stimulus repetition (`IsRep` = 0), a clear fronto-central topography is shown in the prediction accuracy (max *r*(*X*; 0) = 0.047, Figure 3**a**) as well as in the ridge hyperparameter (min log_10_ *λ*(*X*; 0) = 5.86, Figure 3**b**), reflecting the envelope encoding in the bilateral auditory cortices while listening to music. For this particular data, the estimated weights were stronger in the left than right fronto-central channels (Figure 3**c**). With the phase-randomised envelope, as expected, the null prediction accuracy was minimal (max *r*(*U* ; 0) = 0.006, Figure 3**d**; max 𝔼 [*r*(*U* ; 0)] = 0.001, Figure 3**g**) with all channels being highly regularised (min log_10_ *λ*(*U* ; 0) = 10.98, Figure 3**e**; min 𝔼 [log_10_ *λ*(*U* ; 0)] = 12.11, Figure 3**h**).

**Figure 3.**
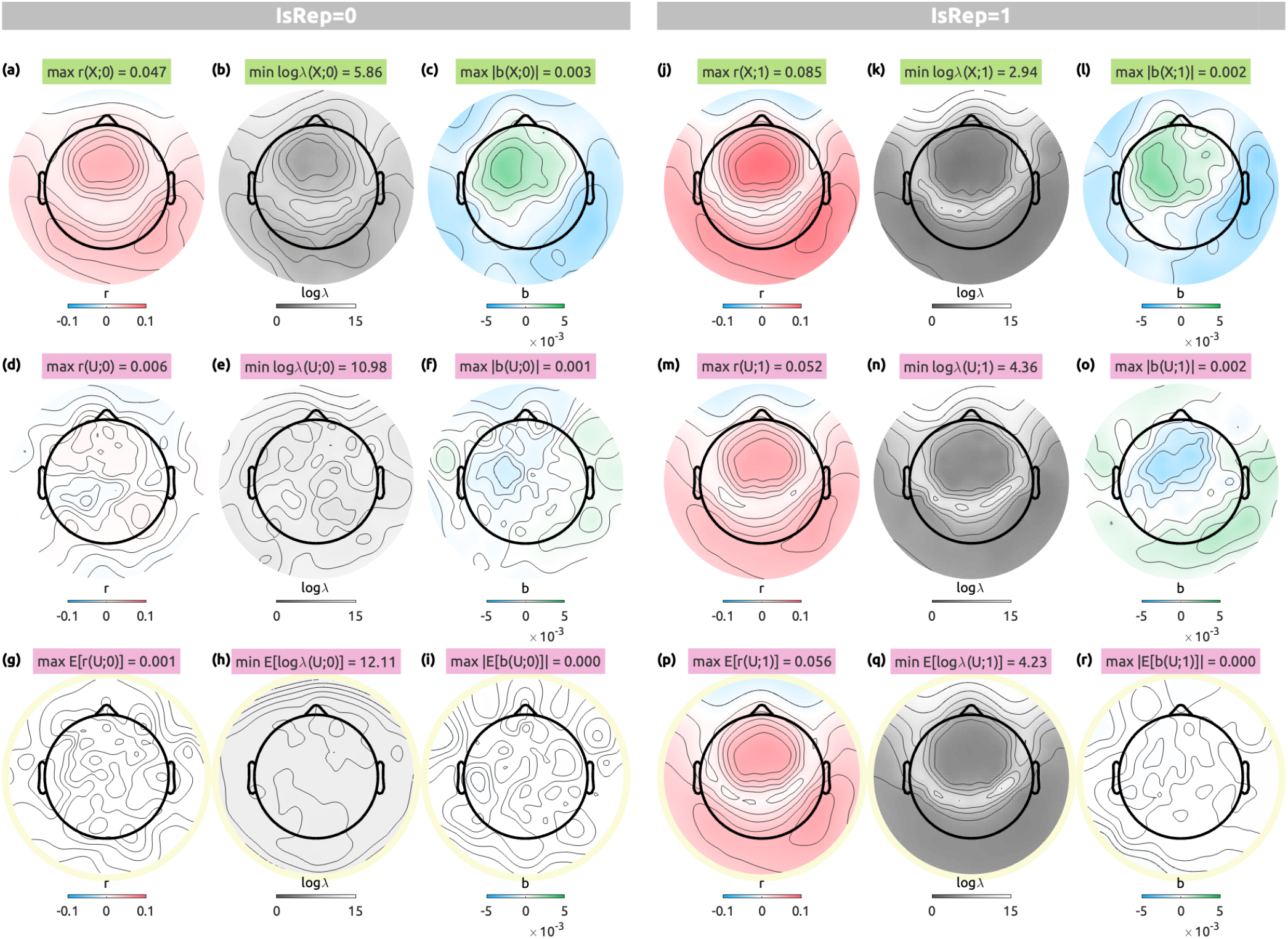
EEG linearised encoding analysis results with delays from 0 to 0.5 sec with an audio envelope (top row, **(a-c, j-l)**), a single case of a phase-randomised envelope (middle row, **(d-f, m-o)**), and an average of 100 phase-randomised envelopes (bottom row, circled in pale yellow, **(g-i, p-r)**). For each CV scheme (`IsRep` = 0, left panels, **(a-i)**; `IsRep` = 1, right panels, **(j-r)**), prediction accuracy (*r*, blue to red, **(a, d, g, j, m, p)**), logarithmic ridge hyperparameter (log_10_ *λ*, gray to white, **(b, e, h, k, n, q)**), transfer function weights that are summed over delays (*b*, blue to green, **(c, f, i, l, o, r)**) are shown along the columns. Note that stimulus repetition not only slightly inflated true prediction accuracies but even the predicted ‘brain activity pattern’ from null features (**m**,**p**), which was effectively indistinguishable from true predictions (**a, j**).

With stimulus repetition (`IsRep` = 1), true prediction accuracies were increased (max *r*(*X*; 1) = 0.085, Figure 3**j**). This is because, unlike the simulation, our feature (i.e., the audio envelope) was not the sole information that the human EEG data encode.

However, most strikingly, the null prediction accuracies with stimulus repetition showed an almost identical topography to the actual encoding results (Figure 3**m**,**p**), with even higher values than the true prediction without repetition (max *r*(*X*; 0) = 0.047, max *r*(*U* ; 1) = 0.052, max 𝔼 [*r*(*U* ; 1)] = 0.056). Note that, by definition, the null feature (phase-randomised envelopes) should not have predicted anything in the EEG data. However, when the identical stimuli were repeated over CV partitions, the regularisation was disabled—regardless of the given features—in channels where the stimulus-evoked response is strong (Figure 3**n**,**q**). Then, even random weights (Figure 3**o**,**r**; Supplementary Figure S11-13; i.e., widely different from the true weights) could successfully predict the repeated signal. Because the SDL artefact reflects the genuine biological signal that is driven by the repeated stimulus, the observed pattern of the prediction accuracy is indistinguishable from the true signal unless investigating the weights. This pattern of SDL was consistently observed, albeit weaker, even when the null features were unrealistic such as normal or uniform noise, or even when different sets of delays were applied (see Supplementary Figures S2–10).

Transfer function weights showed interesting patterns (see also Supplementary Figures S11–13). Spatially, the eigenvectors explaining the largest variance (i.e., PC1) of the phase-randomised envelope were similar to those of the true envelope while the eigenvectors of the uniform or normal noise without autocorrelation showed rather noisy patterns (i.e., in high spatial frequencies). Temporally, even though the uniform or normal noise features had no autocorrelation, the eigenvariate time series showed smooth patterns, reflecting the autocorrelation of the EEG data.

When comparing the null prediction accuracies between the CV schemes (e.g., Figure 3**g** vs. Figure 3**p**), the SDL effects were found to be significant in most channels except for the frontopolar electrodes (*P*_*F DR*_ < 0.01; Figure 4), exhibiting a fronto-central topography that is commonly found in association with the auditory cortical activity. It is noteworthy that the SDL effect is much stronger for the phase-randomised envelope, which preserves the autocorrelation structure of the stimulus. Also, the number of delays seems to further exacerbate the SDL artefact, consistent with the Supplementary Simulation Results.

**Figure 4.**
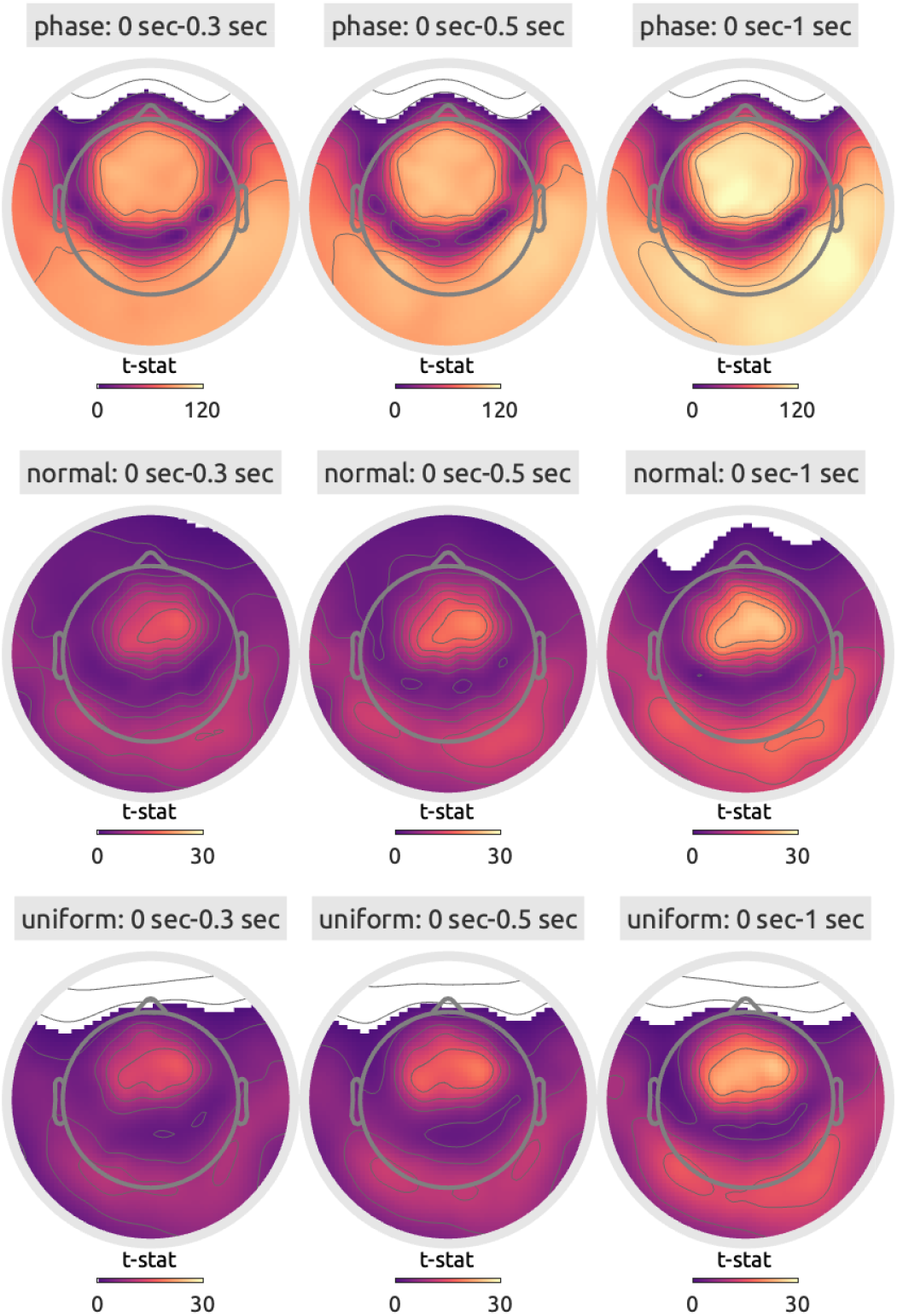
*t*-statistic maps comparing the null prediction accuracies between two CV schemes (e.g., Figure 3**g** vs. Figure 3**p**) to test the SDL effects in the EEG data for the phase-randomised envelope (top row), the normal noise (middle row), and the uniform noise (bottom row). Channels were thresholded by statistical significance with FDR adjustment (one-sided *P*_*F DR*_ < 0.01).

### 4.2 Functional magnetic resonance imaging

Blood-oxygen-level-dependent (BOLD) data were acquired in 39 healthy participants while listening to Western instrumental musical pieces that either evoke happiness or sadness as validated in independent listeners (Sachs et al., 2020). As done for the EEG dataset, linearised encoding analysis was performed with the fMRI data as responses, the audio envelope as a true feature, the phase-randomised envelope as a null feature, and delays from 3 to 9 seconds (Figure 5). Similarly to the EEG results, the phase-randomised envelope strikingly predicted the BOLD time series in the bilateral auditory cortices including the Heschl’s gyrus and planum temporale (Figure 5**m**,**p**) while no consistent pattern in the transfer function weights was found over the phase randomizations (Figure 5**r**), clearly demonstrating the SDL effect. Once again, the anatomical location and the extent of the heightened null prediction accuracies precisely matched the true encoding results (Figure 5**a**,**j**), which would seem ‘highly convincing’ to many neuroscientists.

**Figure 5.**
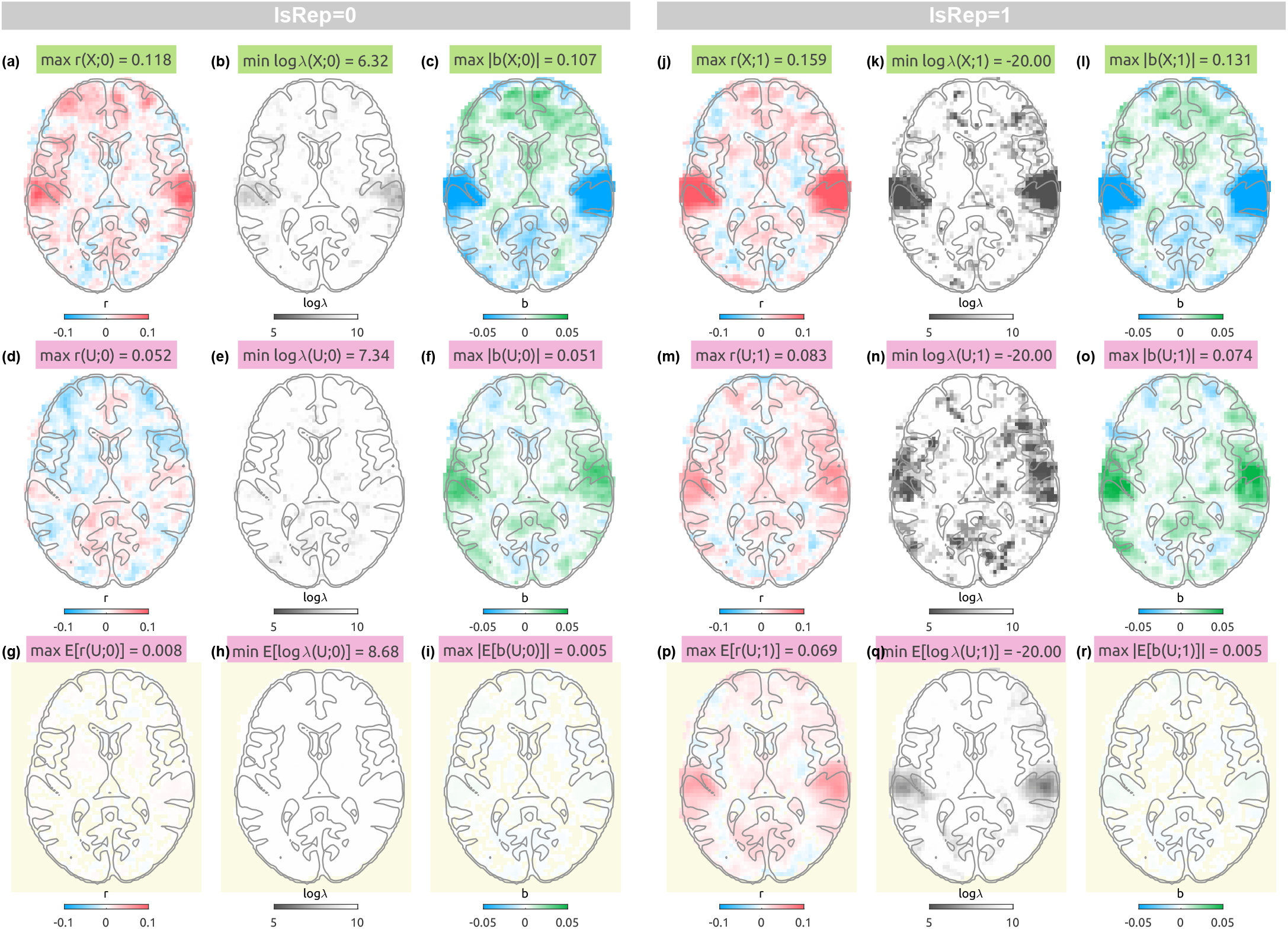
fMRI linearised encoding analysis results with delays from 3 to 9 sec with the audio envelope (top row, **(a-c, j-l)**), a single case of a phase-randomised envelope (middle row, **(d-f, m-o)**), and an average of 100 phase-randomised envelopes (bottom row, pale yellow background, **(g-i, p-r)**). For each CV scheme (`IsRep` = 0, left panels, **(a-i)**; `IsRep` = 1, right panels, **(j-r)**), prediction accuracy (*r*, blue to red, **(a, d, g, j, m, p)**), logarithmic ridge hyperparameter (log_10_ *λ*, gray to white, **(b, e, h, k, n, q)**), transfer function weights that are summed over delays (*b*, blue to green, **(c, f, i, l, o, r)**) are shown along the columns. The analysis was done in the 3-D space, but transverse slices (Montreal Neurological Institute [MNI]-coordinate Z = 8 mm) are chosen to display anatomical structures implicated in a meta-analysis on music-evoked emotions (Koelsch, 2020) such as the Heschl’s gyrus, planum temporale, and the inferior frontal cortex. The 3-D volumes can be viewed with the NeuroVault web viewer (https://identifiers.org/neurovault.collection:19626).

When analysed with different noise models and delays (see Supplementary Figures 14–23), the similar SDL pattern was consistently observed (i.e., inflated prediction accuracies in the auditory cortices). The SDL effect was statistically significant in not only the bilateral superior temporal gyri but the medial occipital cortices and the inferior frontal cortices, where acoustic energy is not expected to be encoded (Figure 6). Consistent with the EEG results, the SDL effect was stronger for the phase-randomised envelope than the normal or uniform noise as well as for longer delays than shorter ones.

**Figure 6.**
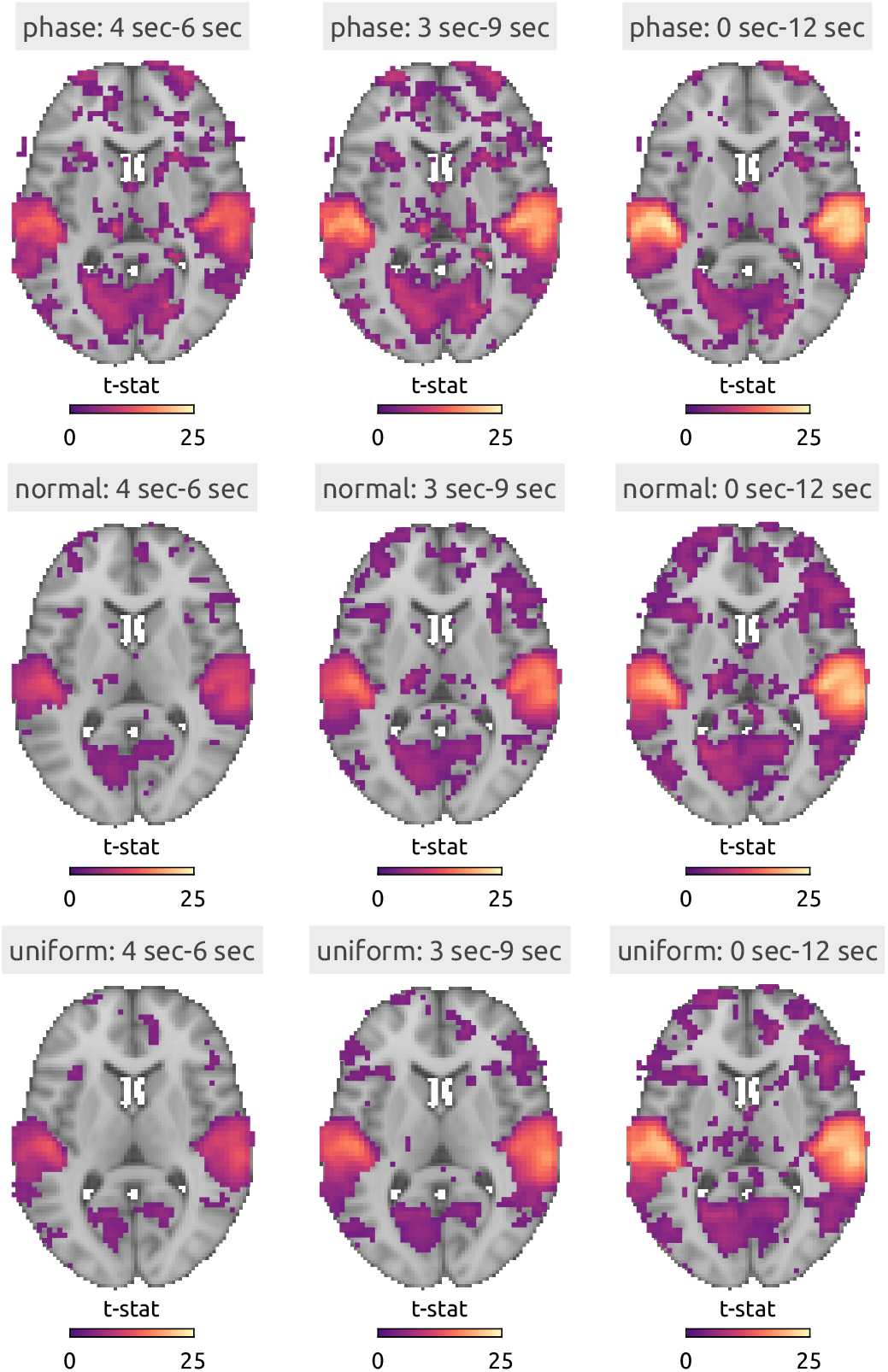
*t*-statistic maps on the transverse slices comparing the null prediction accuracies between two CV schemes (e.g., Figure 5**g** vs. Figure 3**p**) to test the SDL effects in the fMRI data with the phase-randomised envelope (top row), the normal noise (middle row), and the uniform noise (bottom row). Voxels were thresholded by statistical significance after FDR adjustment (one-sided *P*_*F DR*_ < 0.01). The background anatomical image is the MNI template included in FSL (`MNI152 T1 2mm brain.nii.gz`). The 3-D volumes can be viewed with the NeuroVault web viewer (https://identifiers.org/neurovault.collection:19626).

The transfer function weights (Supplementary Figures 24–26) display prominent “auditory components” even for normal and uniform noise in their eigenvectors while the corresponding eigenvariates were widely different from the weights estimated by the true feature.

### 4.3 Behavioural ratings

Continuous ratings of music-evoked emotions were sampled from the same 39 healthy participants who took part in the fMRI experiment above (Sachs et al., 2020). After the scanning session, participants listened to the same musical pieces again and rated their Emotionality (how happy/sad they felt) and Enjoyment (how much they enjoyed the piece) using a slider. A linearised encoding analysis was performed with the ratings as responses, the audio envelope as a true feature, the phase-randomised envelope as a null feature, and delays from 0 to 10 seconds (Figure 7). Once more, while the true envelope predicted Emotionality to some degree and Enjoyment to a greater extent (Figure 7**a**), the phase-randomised envelope also predicted both scales well above zero when the stimuli were repeated across CV partitions (Figure 7**m**,**p**) unlike when the stimuli were not repeated (Figure 7**g**).

**Figure 7.**
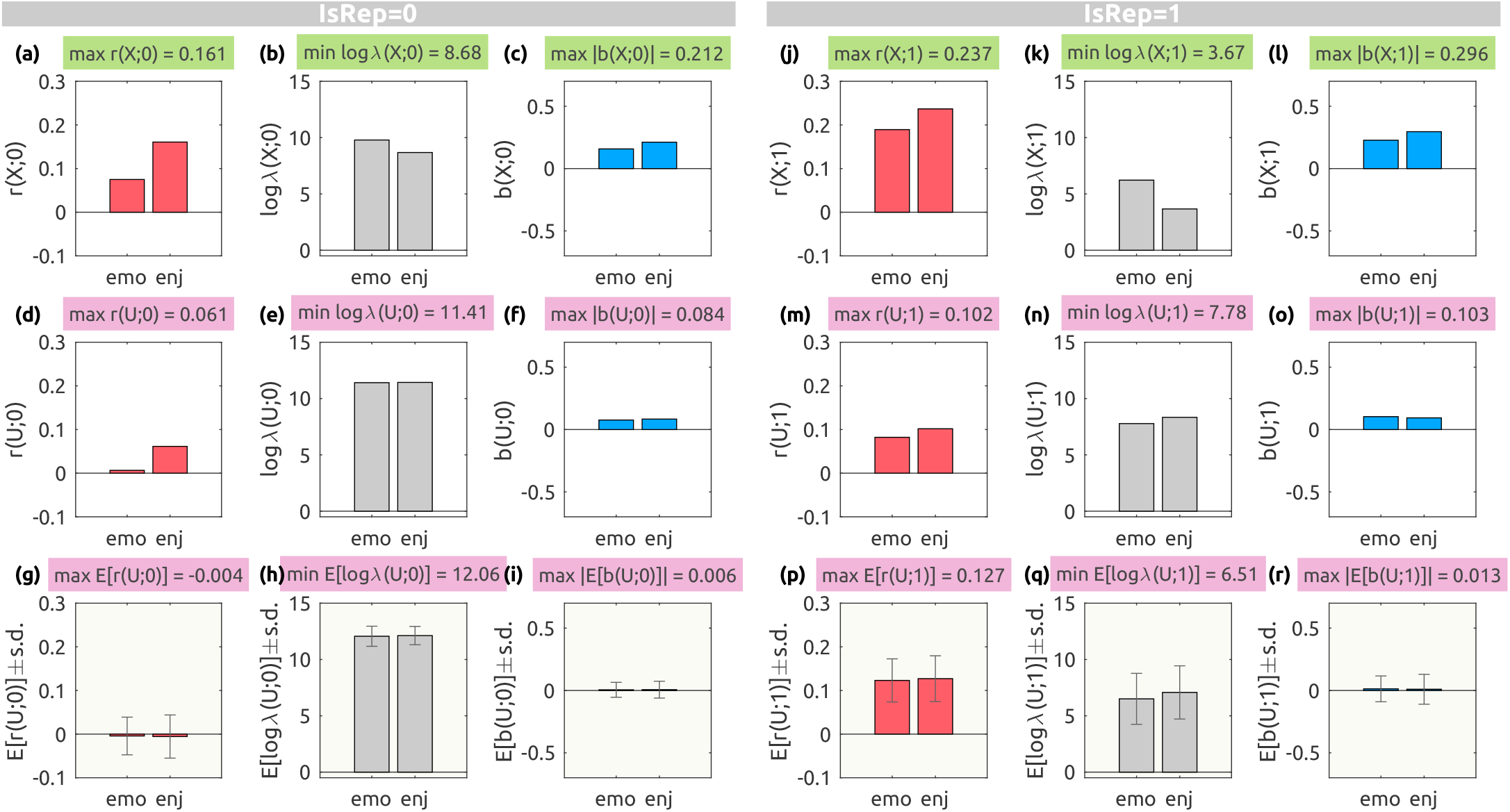
Behavioural linearised encoding analysis results with delays from 0 to 10 sec with an audio envelope (top row, **(a-c, j-l)**), a single case of the phase-randomised envelope (middle row, **(d-f, m-o)**), and an average of 100 phase-randomised envelopes (bottom row, pale yellow background, **(g-i, p-r)**). For each CV scheme (`IsRep` = 0, left panels, **(a-i)**; `IsRep` = 1, right panels, **(j-r)**), prediction accuracy (*r*, red bars, **(a, d, g, j, m, p)**), logarithmic ridge hyperparameter (log_10_ *λ*, gray bars, **(b, e, h, k, n, q)**), transfer function weights that are summed over delays (*b*, blue bars, **(c, f, i, l, o, r)**) are shown along the columns. For the averaged metrics (bottom row), the standard deviations are shown as error bars. Emo.: emotionality, Enj.: enjoyment.

The SDL effect was significant also in the behavioural ratings consistently across all noise models and delays (Figure 8; see also Supplementary Figures 27–34). Similarly to other modalities, the transfer function weights reflected the inherent autocorrelation structure of the behavioural data (Supplementary Figures 35–37).

**Figure 8.**
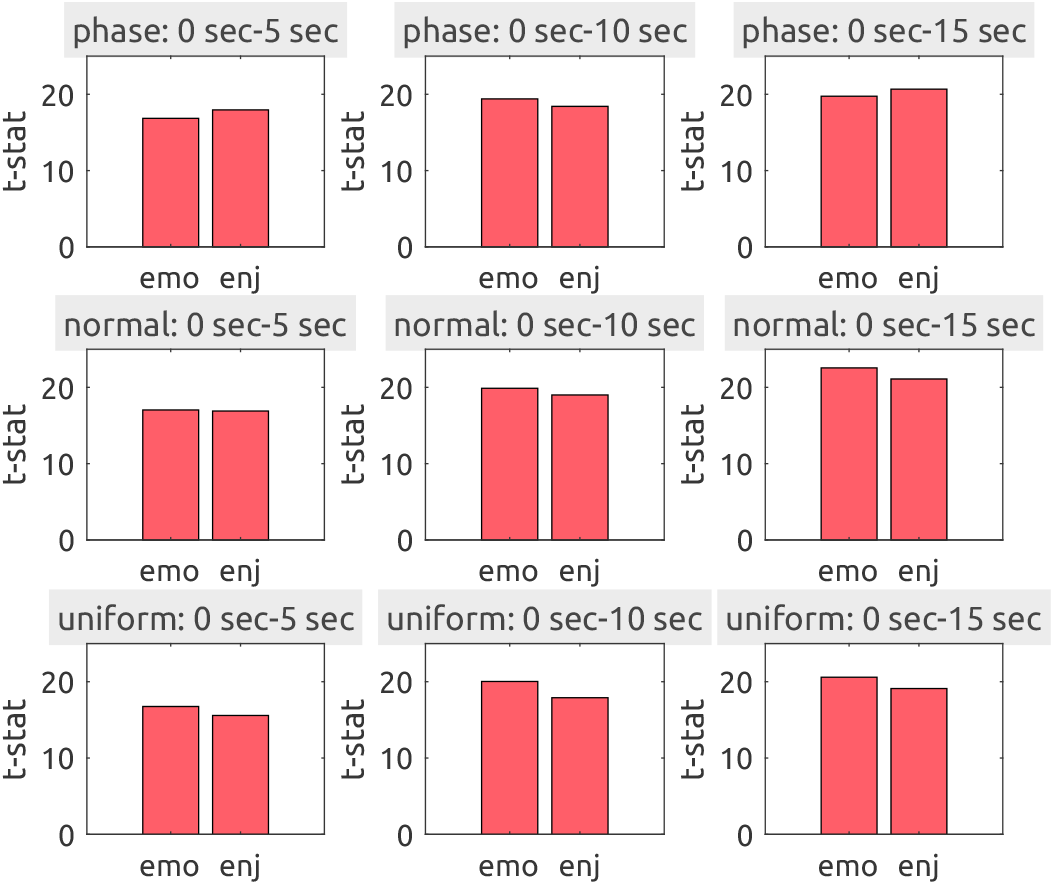
*t*-statistic bar plots comparing the null prediction accuracies between two CV schemes (e.g., Figure 7**g** vs. Figure 3**p**) to test the SDL effects in the behavioural data with the phase-randomised envelope (top row), the normal noise (middle row), and the uniform noise (bottom row). All effects were statistical significant after FDR adjustment (one-sided *P*_*F DR*_ < 0.01). Emo.: emotionality, Enj.: enjoyment.

## 5 Discussion

The primary objective of cognitive neuroscience is to comprehend how the brain executes information-processing operations (Kay, 2018). Linearised encoding analysis serves as a robust method to evaluate a model (i.e., transfer function) that describes how the brain encodes sensory information from the environment and processes this information further (Naselaris et al., 2011). However, it is critical to prevent data leakage when testing the model’s generalisability to unseen data. This paper elucidates how stimulus-driven leakage (SDL) can artificially inflate prediction accuracy and demonstrates realistic cases through extensive simulations and real data analyses.

Firstly, it was mathematically shown that the expected prediction accuracy of the null feature could exceed zero when the null features are identically repeated across CV partitions (Equation 6). In particular, the similarity between training and optimisation sets disables regularisation leading to an inflation of the null prediction accuracy.

Secondly, simulations showed that the SDL artefact is more pronounced with a higher SNR, greater flexibility (i.e., more delay points or higher dimensional features), and more similar autocorrelation structures between the true and null features (Supplementary Simulation results).

Lastly, the SDL artefact was consistently observed across popular data modalities in cognitive neuroscience (i.e., EEG, fMRI, behavioural ratings). In particular, the inflated prediction accuracy exhibited highly plausible spatial patterns even when predicted by uniform noise as a null feature. These patterns are driven by time-locked neural responses to repeated stimuli (widely known as inter-trial/-subject synchrony), not by random noise in the data. Therefore, when combined with *informal reverse inference* (i.e., falsely inferring a mental process from a brain activity pattern without accounting for base rates), which is also a common logical fallacy in cognitive neuroscience (Poldrack, 2006), SDL can lead to completely incorrect conclusions (e.g., a conclusion such as *“The auditory cortex was encoding this uniform random noise that was never presented to the participant*.*”* based on Supplementary Figure 15).

### 5.1 But my features are not just random noise!

For demonstrative purposes, I used random noise to show that the SDL effect can be observed with a feature that is not expected to be encoded in the real data. Certainly, no one would sincerely expect the auditory cortex to encode a random feature that cannot be extracted from the stimulus. In practice, researchers hypothesise that certain information extracted from the stimulus is encoded in the response based on some theoretical and/or empirical grounds. However, SDL can inflate the prediction accuracy of their hypothesised feature significantly regardless of whether the information is actually encoded in the response. This, in turn, can contribute to contamination of the literature and misguide future research.

Also, note that the SDL artefact was greater for the phase-randomised envelope than the normal or uniform noise (Figure 4, Figure 6, Figure 8). This is because the phase-randomised envelope preserves the autocorrelation structure of the stimulus, making it more similar to true features than random noise. Since the hypothesised feature is extracted from actual stimuli, it necessarily bears some resemblance to true features, regardless of the non-linear operations involved. Therefore, the risk of SDL is greater when the hypothesised feature is derived from the stimulus rather than from random noise.

Often nested regression models are compared to determine the unique predictive contribution of a feature of interest. For example, in our previous study (Leahy et al., 2021), the prediction accuracy of a model with an audio envelope (“reduced model”) was subtracted from the prediction accuracy of a model with the envelope and musical beats (“full model”) to test the encoding of musical beats^3^. Even in such cases, SDL can still inflate the prediction accuracy of the full model because the extra features would increase the flexibility of the model. Once again, in well-designed cross-validation without information leakage, irrelevant features would have been regularised (i.e., controlling the flexibility). However, in the presence of SDL, the regularisation is disabled, and the irrelevant features are not penalised, thus leading to an inflated prediction accuracy.

### 5.2 Is SDL relevant to other analyses?

As mentioned earlier, SDL is not unique to linearised encoding analysis or naturalistic neuroimaging. Researchers using other predictive analyses in cognitive neuroscience may therefore wonder whether SDL is relevant to their methods as well. In this section, I briefly review several methods and examine whether they could in principle be susceptible to SDL when applied to datasets with repeatedly presented stimuli. Note that the discussion here is not intended to evaluate or critique specific published studies.

#### 5.2.1 Beta image encoding

Unlike the FIR model that estimates transfer function weights for each time point, the beta image encoding model is on top of the classical GLM that estimates ‘beta’ weights for each stimulus. This approach is popular in visual fMRI experiments where the gazing and visual attention of participants is difficult (or unnecessary) to resolve in time (Kay et al., 2008) or auditory fMRI experiments with short (1–2 seconds) stimuli (Moerel et al., 2018). Thus, instead of directly handling the autocorrelated BOLD time series, an average activation amplitude (i.e., beta weight) during a short trial is first estimated using a GLM, either using a theoretical transfer function called the canonical hemodynamic response function (Henson et al., 1999) or by fitting a regularised FIR model on a split data (Prince et al., 2022).

The beta estimation can be described as a Generalized Least Squares (GLS) problem (Lage-Castellanos et al., 2019):

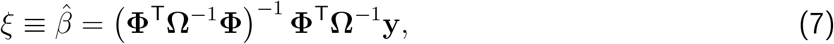

where *ξ* ∈ ℝ^*M* ×1^ is the estimated stimulus-response vector for *M* stimuli, **Φ** ∈ ℝ^*T* ×*M*^ is the cHRF- convolved design matrix for *T* time points, Ω ∈ ℝ^*T* ×*T*^ is the autocovariance matrix of the noise in the BOLD time series, and **y** ∈ ℝ^*T* ×1^ is the BOLD time series.

Once the temporal structures have been dealt with, the estimated beta weights are subjected to the linearised encoding model:

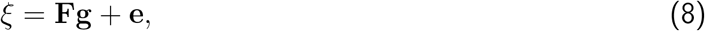

where **F** ∈ ℝ^*M* × *F*^ is a matrix that describes *F* features for *M* presented stimuli, **g** ∈ ℝ^*F* ×1^ is a feature-response vector of the voxel, and **e** ∈ ℝ^*M* ×1^ is unknown noise.

If two sets of beta images with *M* identical stimuli were partitioned into CV folds (e.g., even runs vs. odd runs where all *M* stimuli were presented in each run; i.e., the most common CV design for MVPA), the expected null prediction accuracy would be also positive:

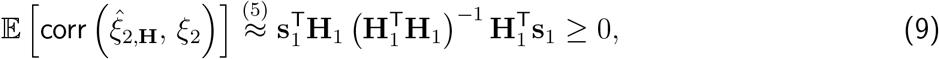

where **s**_*i*_ = **F**_*i*_**g** ∈ ℝ^*M* × 1^ is the underlying signal pattern (a “response profile”) for *M* stimuli in the *i*-th CV partition (*i* = 1, training; *i* = 2, testing), **H**_*i*_ ∈ ℝ ^*M* ×*F*^ is the null feature matrix.

Although it is standard practice to randomize the presentation order of stimuli across runs and participants, the beta image estimation process can reorganize the response profiles, aligning them in a consistent order across all runs (e.g., sequentially from the first to the *M* -th stimuli). Therefore, the risk of SDL in the beta image encoding analysis remains a concern.

#### 5.2.2 Stimulus reconstruction

Reconstruction of unseen stimuli (e.g., images or sounds) based on neural data demonstrates the remarkable potential of neuroimaging techniques for “mind-reading” (Han et al., 2019; Kay et al., 2008; Santoro et al., 2017). In practice, a set of linear models decodes features from neural data (e.g., beta images), followed by a reconstruction step where a simple classifier or a deep neural network (such as variational autoencoder) synthesizes the stimulus from the decoded features.

Since the decoding model is also a linear model like the encoding model, in principle, SDL can also occur. However, the goal of the analysis makes the requirement of “unseen stimuli” more explicitly. For example, if one fit a linear model with *N* − 1 subjects’ responses to Stimulus A, and then tries to recover, once again, Stimulus A from a response of the *i*-th subject, the problem of non-independence between training and test sets may be more visible. Having said that, it is still possible that latent similarity of stimuli is overlooked. Especially if there are too many stimuli to manually inspect, it is possible that some stimuli are indeed non-identical but highly similar (e.g., utterances by the same speaker; repetitions of musical phrases). Checking inter-stimulus correlation for all features prior to partitioning them into training and test sets will prevent such cases of hidden similarity.

#### 5.2.3 Multivariate classification

Multivariate classification analysis has been widely used in the cognitive neuroscience community, commonly known as multivoxel (or multivariate) pattern analysis (MVPA; Kriegeskorte et al., 2006) or single-trial classification in electrophysiological data such as EEG, MEG, and ECoG (Müller-Gerking et al., 1999; Pistohl et al., 2012; Quandt et al., 2012), either on the whole set of response units (e.g., whole-brain classification; Ryali et al., 2010) or local neighbors (e.g., a “searchlight”; Kriegeskorte et al., 2006). While a highly accurate classifier is necessary to build a brain-computer interface system, classification analysis can be useful even with low but significant accuracy—as is often the case in cognitive neuroscience research—since such a classifier still functions as a multivariate model that tests the existence of certain information in the brain.

Classification can be seen as model-free as compared to encoding or decoding analysis because classification does not require a definition of features. Note that the trained weight vector only represents a separation of given training examples in the response space, regardless of the physical or semantic features of the stimuli. For example, one may try to classify ‘happy music’ vs. ‘sad music’ based on the EEG responses. However, if the chosen exemplars of the classes are imbalanced (e.g., all exemplars of ‘happy music’ naturally happened to be faster and louder than ‘sad music’), classification could be strongly driven by different acoustics, not necessarily due to perceived or evoked emotions from the music. That is, without carefully matching the training examples for all relevant features, the interpretation of classification analysis may be confounded.

Moreover, in principle, the repetition of stimuli (or at least classes) across data points is not just unavoidable but in fact necessary for the classification. In order to train a classifier, not only linear but also non-linear one, balanced training and testing examples of all classes are required (Hastie et al., 2009). In other words, it is impossible to train a classifier for an ‘unseen class’. A classifier can only classify an ‘unseen instance’ of a known class. Therefore, SDL does not apply to classification analysis, although other forms of data leakage—such as global preprocessing or feature selection before split—remain a great concern.

#### 5.2.4 Representation similarity analysis

Finally, let us consider a method that evaluates the second-order isomorphism across representational systems (Kriegeskorte et al., 2008), widely known as representational similarity analysis (RSA). This flexible method can define a ‘model’ distance matrix between stimuli either based on class labels (as in classification analysis) or feature descriptors (as in encoding analysis). The original formulation of RSA does not involve cross-validation, as the RSA itself is neither a predictive model nor a classification method (Kriegeskorte et al., 2008). A later extension introduced a cross-validated, squared Mahalanobis distance estimator—known as “crossnobis”—to enhance both reliability and interpretability (Diedrichsen & Kriegeskorte, 2017). When estimating a crossnobis distance between two conditions in a leave-one-out cross-validation (LOOCV) scheme, it is assumed that the number of responses for both conditions remains balanced across all CV partitions. Thus, if identical stimuli are repeated across partitions, the crossnobis distance for the brain representation distance matrix (RDM) would be biased by the stimulus-specific (rather than condition-specific) activity. While this is not over-fitting to identical noise, it would be over-fitting to the idiosyncratic signal of a particular stimulus. However, even in this case, null descriptors would define a model RDM that is irrelevant to the brain RDM. Therefore, a false conclusion that “a brain region represents null information” would not be supported by the association between the model and brain RDMs. Thus, the risk of SDL in RSA appears minimal.

### 5.3 How do we detect and prevent SDL?

Since avoiding data leakage in predictive analysis is non-trivial for scientists without a machine learning background, valuable pedagogical resources such as Bernett et al., 2024 have been developed to propose guiding questions that help researchers identify and avoid data leakage. Here, I discuss ways to detect SDL and alternative analyses and designs that can prevent it.

#### 5.3.1 Algorithmic detection

As shown earlier, SDL occurs when the identical stimulus is presented more than once across CV partitions. Thus, a simple way to detect the risk of SDL would be looking for the identical (or similar) data pairs in the dataset, before partitioning the data into training, validation, and test sets.

This was already illustrated in the toy example (see Supplementary Results: Simulation). When no stimulus was repeated across CV partitions the inter-trial correlation (ITC) was on average zero across 200 random samplings. However, when the stimulus was repeated across CV partitions, the ITC was on average about 0.8. Because this correlation can be cheaply computed prior to costly optimisation and modelling fitting, this can be a useful diagnostic tool to assess risk for SDL.

Checking ITC is also useful to detect latent similarity across stimuli as well. For example, two audio files are named differently but contain similar music, the researcher would not know about the leakage in the training examples. The feature-ITC can be a useful tool to detect such latent similarity.

For users’ convenience, an automatic validation test for a given CV design based on feature-ITC and response-ITC is implemented as a default option in the MATLAB package for Linearised Encoding Analysis (LEA; https://github.com/seunggookim/lea).

#### 5.3.2 Alternative analyses

If a risk of SDL is detected in your planned analysis, what should you have to do next? One way to handle it find an alternative CV design that does not introduce SDL.

##### Subject-wise modelling

As shown in Figure 1, if possible, an easy solution is to adopt subject-wise modelling instead of stimulus-wise modelling. Unless there exists strong similarity between subjects (e.g., identical twins, i.e., literal ‘twinning’) in different CV partitions, subject-wise modelling can avoid SDL.

##### Averaging responses

In practice, acquiring high-quality neuroimaging data remains time-consuming and expensive, while its SNR—particularly for non-invasive in-vivo methods—remains poor. Moreover, acquiring extensive data from vulnerable populations (e.g., patients, young children, elderly people) poses not only financial and logistical but also ethical concerns. Thus, many researchers are motivated to aggregate the limited data across multiple subjects to further reduce the sampling variance, as in stimulus-wise modelling.

If the noise ceiling is deemed undesirable for subject-wise modelling (see Lage-Castellanos et al., 2019 for estimation methods), one can average responses for the identical stimulus before the encoding analysis. For example, if an identical set of stimuli is used for all participants, one can average participants’ responses for each stimulus to create an ‘average-subject’. Then, subject-level inference can be performed (e.g., using randomised features that preserve autocorrelation, such as linear shifting or phase randomisation).

#### 5.3.3 Alternative designs

Moreover, if this risk is recognised at an early stage of the study (e.g., during study design or after a few pilot sessions), one can consider alterative designs that deter SDL.

##### Hold-out validation

A straightforward but expensive (i.e., requiring more data) way to prevent SDL is to use *hold-out validation* instead of cross-validation. In hold-out validation, the test set is used only once to estimate the prediction performance. Thus, the test set is deliberately designed to contain different stimuli, even before data acquisition. Because the testing accuracy is proportional to the signal strength of the test set (see Supplementary Equation 11), it is desirable to design the test set to have high signal strength (e.g., by averaging multiple presentations; Han et al., 2019; Huth et al., 2016; Nishimoto et al., 2011).

##### Single-use stimulus

Another design to prevent SDL is to use a stimulus only once during the whole study. That is, a stimulus is presented to only one participant, only once, and never used again (i.e., a single-use stimulus). This design prevents any accidental stimulus repetition across CV partitions and thus avoids the risk of SDL in any CV designs. However, as mentioned above, the poor SNR of non-invasive neuroimaging data may need to be addressed. In that case, a stimulus may be presented multiple times to only one participant, but then the responses must be averaged across trials before any predictive analysis.

### 5.4 Limitations of the current paper

While the current paper provides a comprehensive analysis of SDL, some limitations should be acknowledged.

First, the paper focuses on a particular, relatively simple model—single-penalty ridge regression. That is, other types of regularisation (e.g., multi-penalty ridge, LASSO, elastic net) and non-linear modelling techniques (e.g., k-nearest neighbours regression, support vector regression, non-linear kernel ridge regression) were not explored. While the key mechanism of SDL (i.e., disabling regularisation due to the similarity in the underlying signal between the training and validation sets) is likely to be present in these other methods, determining the extent to which SDL influences them requires further analysis.

Second, no neural spike data or intracranial recordings—which are gradually becoming more accessible in human patients—were analysed in this paper. The SDL effect was only demonstrated in the context of continuous-valued data acquired from non-invasive methods (e.g., EEG, fMRI, and behavioural ratings). Sparse spike data or firing rate data may behave differently from continuous-valued data. However, given that the SDL effect is driven by similarity in the underlying signal, it is likely to occur when the raw data are processed so that stimulus-evoked, time-locked responses are strongly present (e.g., high-gamma power envelopes of local neuronal populations; Jacobs & Kahana, 2009; Ray et al., 2008).

### 5.5 Conclusion

The current paper shows that SDL is a critical but under-recognized risk in encoding analysis and stimulus reconstruction. SDL may lead to spurious conclusions suggesting that irrelevant information is encoded in neural signals. When carefully designed, the model-based approach remains a powerful information-based tool for naturalistic neuroimaging.

## Data and Code Availability

All analysis code, simulation data, and real data analysis results are available at Zenodo: https://doi.org/10.5281/zenodo.15100830. An executable demo of simulation is available at CodeOcean: https://codeocean.com/capsule/4591394/. The EEG raw dataset is available at Standford Digital Repository: https://purl.stanford.edu/sd922db3535. The fMRI and behavioural raw datasets are available at Open-Neuro: https://openneuro.org/datasets/ds003085. A MATLAB package for Linearised Encoding Analysis (LEA) on multimodal data is available at Zenodo release: https://doi.org/10.5281/zenodo.15107756 and GitHub: https://github.com/seunggookim/lea. A freely executable demo of LEA on real data is available at MATLAB Online: https://s.gwdg.de/7cOQmw. fMRI analysis results in 3-D NIfTI format can be viewed and downloaded at NeuroVault: https://identifiers.org/neurovault.collection:19626.

## Author Contributions

*Seung-Goo Kim:* Conceptualization, Methodology, Software, Validation, Formal analysis, Investigation, Data Curation, Writing—Original Draft, Writing—Review & Editing, Visualization.

## Funding

Max Planck Society supported this work.

## Declaration of Competing Interests

The author declares that the research was conducted in the absence of any commercial or financial relationships that could be construed as a potential conflict of interest.

## Acknowledgements

The author thanks Dr. Daniela Sammler, Dr. Vincent Shi-chen Chien, Mr. Seung-Cheol Baek, and three anonymous reviewers for their constructive feedback and expert insights on earlier versions of the manuscript. The author acknowledges the free image sources from www.irasutoya.com (Takashi Mifune), which reserves the copyrights of the source illustrations.

## Footnotes

1 While *feature* and *predictor* are often interchangeably used, in this article a feature refers to a variable that describes the characteristics of interest of the input, while a predictor refers to a feature with a specific delay (i.e., each column of a design matrix). Thus, *F* features with *D* delays produce *P* = *FD* predictors.

2 This paper focuses on the FIR model for its popularity in time-invariant linear system identification (Crosse et al., 2021; Huth et al., 2016; Wu et al., 2006). Nonetheless, the conclusion of this section generalises to any design matrix (i.e., **X** does not need to be a Toeplitz matrix).

3 While the comparison of nested models is a standard practice in Ordinary Least Squares (OLS) regression, it can be complicated with regularised regression such as ridge. The major problem is over-regularisation of relevant features due to irrelevant features in the full model when a single penalty hyperparameter is used for all features (feature spaces). A multi-penalty model can address this issue better, e.g., Kim et al. (2024) and La Tour et al. (2022).

## Notes

https://zenodo.org/records/15100830

https://s.gwdg.de/7cOQmw

https://neurovault.org/collections/19626

https://codeocean.com/capsule/4591394

### Competing Interest Statement

The authors have declared no competing interest.

### Summary of Updates

This revision improved the previous manuscript in terms of clarity and brevity.

