## Supplementary Materials for "Stimulus-Driven Leakage in Naturalistic Neuroimaging"

### Supplementary Materials for "Stimulus-Driven Data Leakage in Naturalistic Neuroimaging"

Seung-Goo Kim*

Research Group Neurocognition of Music and Language,

Max Planck Institute for Empirical Aesthetics, Grüneburgweg 14, 60322 Frankfurt, Germany

March 24, 2026

#### S1 Supplementary Theory

##### S1.1 Finite impulse response model

A finite impulse response (FIR) model is commonly used to identify a time-invariant linear system. Please see Table 1 for the definitions of variables and notations used in this article. Let us consider an FIR model:

$$\mathbf{y} = \mathbf{X}\mathbf{b} + \mathbf{e}, \tag{S1}$$

where $\mathbf{y} \in \mathbb{R}^{T\times 1}$ is a response vector over $T$ time points of a single response unit (e.g., a channel or a voxel), $\mathbf{X} \in \mathbb{R}^{T\times FD}$ is a nonzero Toeplitz design matrix of $F$ features[1] with $D$ delays such that $\|\mathbf{x}_{(i)}^{\mathsf{T}}\mathbf{x}_{(i)}\| > 0$ for all $i \in \{1, \cdots, FD\}$, $\mathbf{x}_{(i)}$ is the $i$-th column of $\mathbf{X}$, $\|\cdot\|$ denotes the $l_2$-norm, which leads to $\|\mathbf{X}^{\mathsf{T}}\mathbf{X}\|_{\mathrm{F}} > 0$ where $\|\cdot\|_{\mathrm{F}}$ denotes the Frobenius norm. $\mathbf{b} \in \mathbb{R}^{FD\times 1}$ is an unknown weight vector (often called a temporal response function; or more generally a transfer function), $\mathbf{e} \in \mathbb{R}^{T\times 1}$ is a zero-mean, unit-variance Gaussian noise vector $\mathbf{e} \sim \mathcal{N}_T(\mathbf{0}, \mathbf{I}_T)$ where $\mathbf{I}_T \in \mathbb{R}^{T\times T}$ is an identity matrix. For convenience, we further assume that we standardise predictors and response variables prior to analysis

[1]While *feature* and *predictor* are often interchangeably used, here a feature refers to a variable that describes the characteristics of interest of the input, while a predictor refers to a feature with a specific delay (i.e., each column of a design matrix). That is, $F$ features and $D$ delays produce $P = FD$ predictors.

Table 1: Definition of variables, parameters, and notations.

| Variables | Description |
|---|---|
| $\mathbf{X} \in \mathbb{R}^{T \times FD}$ | True finite impulse response design matrix |
| $\mathbf{U} \in \mathbb{R}^{T \times FD}$ | Null finite impulse response design matrix |
| $\mathbf{y} \in \mathbb{R}^{T \times 1}$ | Response vector |
| $\mathbf{b} \in \mathbb{R}^{FD \times 1}$ | Regression coefficient vector |
| $\mathbf{s} \in \mathbb{R}^{T \times 1}$ | Signal time series |
| $\mathbf{e} \in \mathbb{R}^{T \times 1}$ | Noise time strategies |
| $\mathbf{L} \in \mathbb{R}^{FD \times FD}$ | Tikhonov regularization matrix |
| **Parameters** | **Description** |
| $T$ | Number of time points |
| $F$ | Number of features |
| $D$ | Number of delays |
| $P$ | Number of predictors ($= FD$) |
| $\phi$ | AR(1) temporal correlation parameter |
| $\theta$ | AR(1) spatial correlation parameter |
| $\rho$ | Multicollinearity parameter |
| $\lambda$ | regularization parameter |
| **Notations** | **Description** |
| $\lVert \cdot \rVert$ | $l_2$-norm of a vector |
| $\lVert \cdot \rVert_{\mathrm{F}}$ | Frobenius norm of a matrix |
| $(\cdot)$T | Transposition |
| $(\cdot)_{(i)}$ | $i$-th column vector of a matrix |
| $\mathrm{diag}(\cdot)$ | Diagonal matrix with the diagonal elements from the given vector |
| $\mathcal{L}(\cdot;\cdot)$ | Optimization function |
| $\cdot^*$ | Optimal solution |
| $\hat{\cdot}$ | Estimate of a variable |
| $(\cdot)_i$ | $i$-th cross-validation set |
| $\bar{\cdot}$ | Mean |
| $\mathbb{E}[\cdot]$ | Expectation of a random variable |
| $\propto$ | Proportional to |
| $\approx$ | Approximately equal to |
| $\equiv$ | Equivalent to |

so that their sample means are zero and sample variances are one.

Here, let us assume that we have access to the true weights, which are nonzero (i.e., $\|\mathbf{b}\| > 0$). With this model, we can generate a training dataset and a test dataset using the identical weights $\mathbf{b}$ but independent predictors and noise:

$$\mathbf{y}_i = \mathbf{X}_i\mathbf{b} + \mathbf{e}_i = \mathbf{s}_i + \mathbf{e}_i, \tag{S2}$$

where $(\cdot)_i$ denotes the $i$-th independent partition in cross-validation with $i = 1$ for a training set and $i = 2$ for a test set. The true signal is denoted as $\mathbf{s}_i \equiv \mathbf{X}_i\mathbf{b} \in \mathbb{R}^{T\times 1}$.

To avoid overfitting to noise, regularisation such as $l_2$-norm penalty (i.e., ridge penalty) is often used:

$$\hat{\mathbf{b}} = \left(\mathbf{X}_1^\mathsf{T}\mathbf{X}_1 + \mathbf{L}\right)^{-1}\mathbf{X}_1^\mathsf{T}\mathbf{y}_1, \tag{S3}$$

where $\mathbf{L} \in \mathbb{R}^{FD\times FD}$ is a Tikhonov regularisation matrix. In the usual case of ridge regression (i.e., a single penalty applied to all predictors), $\mathbf{L} = \lambda\mathbf{I}_{FD}$. In the general case of multi-penalty ridge (Hoerl & Kennard, 1970), predictor-delay-wise penalties can be defined: $\mathbf{L} = \text{diag}(\lambda_1, \lambda_2, \ldots, \lambda_{FD})$. For now, we assume a single penalty applies to all predictors, except for the intercept, which remains unregularised.

In practice, the hyperparameter (i.e., $\lambda$) is typically optimised using the third, independent partition of data, known as a *validation* set in statistical learning (Hastie et al., 2009).

$$\hat{\mathbf{b}} = \left(\mathbf{X}_1^\mathsf{T}\mathbf{X}_1 + \mathcal{L}(\{\mathbf{X}_1, \mathbf{y}_1\}; \{\mathbf{X}_3, \mathbf{y}_3\})\right)^{-1}\mathbf{X}_1^\mathsf{T}\mathbf{y}_1, \tag{S4}$$

where $\mathcal{L}(\cdot;\cdot)$ is an optimiser that finds the *optimal* regularisation matrix $\mathbf{L}^*$ minimising prediction error for given pairs of design and response variables $\{\mathbf{X}_i, \mathbf{y}_i\}$ and $\{\mathbf{X}_j, \mathbf{y}_j\}$. With this, a regularised prediction for $\mathbf{y}_2$ based on $\mathbf{X}$ is:

$$\hat{\mathbf{y}}_{2,\mathbf{X}} = \mathbf{X}_2\hat{\mathbf{b}} = \mathbf{X}_2\left(\mathbf{X}_1^\mathsf{T}\mathbf{X}_1 + \mathbf{L}^*\right)^{-1}\mathbf{X}_1^\mathsf{T}\mathbf{y}_1 = \mathbf{P}_\mathbf{X}\mathbf{y}_1, \tag{S5}$$

where $\mathbf{P}_\mathbf{X} \in \mathbb{R}^{T\times T}$ is a regularised projection matrix based on $\mathbf{X}$. Because of the standardisation ($\|\mathbf{y}\| = 1$) and the nonnegativity of the denominator, the expected value of a prediction accuracy metric (e.g., Pearson correlation) over random noise can be seen as proportional to the expected inner product of the prediction and the response:

$$\mathbb{E}\left[\text{corr}\left(\hat{\mathbf{y}}_{2,\mathbf{X}},\, \mathbf{y}_2\right)\right] = \mathbb{E}\left[\frac{\left(\hat{\mathbf{y}}_{2,\mathbf{X}}\right)^\mathsf{T}\mathbf{y}_2}{\|\hat{\mathbf{y}}_{2,\mathbf{X}}\|\|\mathbf{y}_2\|}\right] \propto \mathbb{E}\left[\left(\hat{\mathbf{y}}_{2,\mathbf{X}}\right)^\mathsf{T}\mathbf{y}_2\right]. \tag{S6}$$

This can be further expanded as

$$
\begin{aligned}
\mathbb{E}\left[(\hat{\mathbf{y}}_{2,\mathbf{X}})^{\mathsf{T}}\,\mathbf{y}_2\right] &= \mathbb{E}\left[\mathbf{y}_1^{\mathsf{T}}\mathbf{P}_{\mathbf{X}}^{\mathsf{T}}\mathbf{y}_2\right] \\
&= \mathbb{E}\left[(\mathbf{s}_1+\mathbf{e}_1)^{\mathsf{T}}\mathbf{P}_{\mathbf{X}}^{\mathsf{T}}(\mathbf{s}_2+\mathbf{e}_2)\right] \\
&= \mathbf{s}_1^{\mathsf{T}}\mathbf{P}_{\mathbf{X}}^{\mathsf{T}}\mathbf{s}_2 + \mathbb{E}\left[\mathbf{e}_1^{\mathsf{T}}\right]\mathbf{P}_{\mathbf{X}}^{\mathsf{T}}\mathbf{s}_2 + \mathbf{s}_1^{\mathsf{T}}\mathbf{P}_{\mathbf{X}}^{\mathsf{T}}\mathbb{E}\left[\mathbf{e}_2\right] + \mathbb{E}\left[\mathbf{e}_1^{\mathsf{T}}\right]\mathbf{P}_{\mathbf{X}}^{\mathsf{T}}\mathbf{e}_2,
\end{aligned} \tag{S7}
$$

where only the first term remains nonzero since $\mathbb{E}\left[\mathbf{e}\right] = \mathbf{0}$. That is,

$$
\mathbb{E}\left[\mathsf{corr}\left(\hat{\mathbf{y}}_{2,\mathbf{X}},\, \mathbf{y}_2\right)\right] \propto \mathbf{s}_1^{\mathsf{T}}\mathbf{P}_{\mathbf{X}}^{\mathsf{T}}\mathbf{s}_2. \tag{S8}
$$

With a sufficiently strong signal $\|\mathbf{b}\| \gg 0$, the optimal regularisation approaches zero: $\mathbf{L}^* \approx \mathbf{0}$ (Hastie et al., 2009), which allows for approximating the projection matrix as:

$$
\mathbf{P}_{\mathbf{X}} = \mathbf{X}_2\left(\mathbf{X}_1^{\mathsf{T}}\mathbf{X}_1 + \mathbf{L}^*\right)^{-1}\mathbf{X}_1^{\mathsf{T}} \approx \mathbf{X}_2\left(\mathbf{X}_1^{\mathsf{T}}\mathbf{X}_1\right)^{-1}\mathbf{X}_1^{\mathsf{T}}. \tag{S9}
$$

Consequently, Equation S8 can be approximated as

$$
\mathbb{E}\left[\mathsf{corr}\left(\hat{\mathbf{y}}_{2,\mathbf{X}},\, \mathbf{y}_2\right)\right] \propto \mathbf{s}_1^{\mathsf{T}}\mathbf{P}_{\mathbf{X}}^{\mathsf{T}}\mathbf{s}_2 \overset{(S9)}{\approx} (\mathbf{X}_1\mathbf{b})^{\mathsf{T}}\left\{\mathbf{X}_2\left(\mathbf{X}_1^{\mathsf{T}}\mathbf{X}_1\right)^{-1}\mathbf{X}_1^{\mathsf{T}}\right\}^{\mathsf{T}}(\mathbf{X}_2\mathbf{b}). \tag{S10}
$$

Due to the symmetry of the inverse covariance $\left\{\left(\mathbf{X}_i^{\mathsf{T}}\mathbf{X}_i\right)^{-1}\right\}^{\mathsf{T}} = \left\{\left(\mathbf{X}_i^{\mathsf{T}}\mathbf{X}_i\right)^{\mathsf{T}}\right\}^{-1} = \left(\mathbf{X}_i^{\mathsf{T}}\mathbf{X}_i\right)^{-1}$ and the nonzero assumption $\|\mathbf{X}_i^{\mathsf{T}}\mathbf{X}_i\|_{\mathrm{F}} > 0$, this can be further simplified:

$$
\begin{aligned}
\mathbb{E}\left[\mathsf{corr}\left(\hat{\mathbf{y}}_{2,\mathbf{X}},\, \mathbf{y}_2\right)\right] \propto \mathbf{s}_1^{\mathsf{T}}\mathbf{P}_{\mathbf{X}}^{\mathsf{T}}\mathbf{s}_2 &\approx \mathbf{b}^{\mathsf{T}}\mathbf{X}_1^{\mathsf{T}}\left[\mathbf{X}_1\left\{\left(\mathbf{X}_1^{\mathsf{T}}\mathbf{X}_1\right)^{-1}\right\}^{\mathsf{T}}\mathbf{X}_2^{\mathsf{T}}\right]\mathbf{X}_2\mathbf{b} \\
&= \mathbf{b}^{\mathsf{T}}\left(\mathbf{X}_1^{\mathsf{T}}\mathbf{X}_1\right)\left(\mathbf{X}_1^{\mathsf{T}}\mathbf{X}_1\right)^{-1}\mathbf{X}_2^{\mathsf{T}}\mathbf{X}_2\mathbf{b} \\
&= \mathbf{b}^{\mathsf{T}}\mathbf{X}_2^{\mathsf{T}}\mathbf{X}_2\mathbf{b} = \|\mathbf{X}_2\mathbf{b}\| = \|\mathbf{s}_2\| > 0.
\end{aligned} \tag{S11}
$$

That is, with a sufficiently strong signal, the prediction accuracy is expected to be positive.

#### S1.2 Null model

In the above, we assumed that we have access to the true predictors $\mathbf{X}$ and true weights $\mathbf{b}$ unlike many real-world scenarios. Please note that the predictors in the current setting are not objectively observable conditions (e.g., the presence or absence of sounds) but a selected set of features of the stimulus that are hypothesized to be relevant to the neural responses by the researchers (e.g., spectro-temporal modulation). In practice, while we clearly know which stimulus we presented, defining predictors requires knowledge of which information is encoded in the human brain, which is ultimately unknown and often is the very aim of the study (e.g., *"Is information X encoded in the brain region A or not?"*).

Replication of the entire analysis with a null model is a common approach to infer statistical significance of the observation. Suppose we use a Gaussian noise as null features $\mathbf{u}_i \sim \mathcal{N}(\mathbf{0}, \sigma\mathbf{I})$ and to delay them to create a Toeplitz matrix $\mathbf{U}_i \in \mathbb{R}^{T \times FD}$ to make their prediction for $\mathbf{y}_2$ as

$$\hat{\mathbf{y}}_{2,\mathbf{U}} = \mathbf{U}_2\hat{\mathbf{b}}_0 = \mathbf{U}_2 \left(\mathbf{U}_1^\mathsf{T}\mathbf{U}_1 + \mathcal{L}(\{\mathbf{U}_1, \mathbf{y}_1\}; \{\mathbf{U}_3, \mathbf{y}_3\})\right)^{-1} \mathbf{U}_1^\mathsf{T}\mathbf{y}_1 = \mathbf{P}_\mathbf{U}\mathbf{y}_1. \tag{S12}$$

Note that actual predictors that researchers would use based on theories, prior evidence, and intuitions should be much more informative than this null features. That is, $\mathbf{U}$ is expected to perform worse than any reasonable predictors. Put differently, if we can somehow *magically* make *significant* predictions using the null predictors $\mathbf{U}$, it indicates there is something critically flawed in our analysis.

As we assume that the null predictors $\mathbf{U}$ are independently generated from the true predictors $\mathbf{X}$, a valid optimisation process such as cross-validation (Hastie et al., 2009) would lead to a strong regularisation of non-informative predictors (i.e., $\mathbf{U}$). This simplifies the projection matrix as a scaled inner product of two null feature matrices:

$$\mathbf{P}_\mathbf{U} = \mathbf{U}_2 \left(\mathbf{U}_1^\mathsf{T}\mathbf{U}_1 + \lambda^*\mathbf{I}\right)^{-1} \mathbf{U}_1^\mathsf{T} \overset{\lambda^* \gg 0}{\approx} \mathbf{U}_2(\lambda^*\mathbf{I})^{-1}\mathbf{U}_1^\mathsf{T} = \frac{1}{\lambda^*}\mathbf{U}_2\mathbf{U}_1^\mathsf{T}. \tag{S13}$$

Thus, given the $\mathbf{U}_i$, the expected value of the prediction accuracy over random noise is given as

$$\begin{aligned} \mathbb{E}\left[\mathsf{corr}\left(\hat{\mathbf{y}}_{2,\mathbf{U}},\, \mathbf{y}_2\right)\right] &\propto \mathbf{s}_1^\mathsf{T}\mathbf{P}_\mathbf{U}^\mathsf{T}\mathbf{s}_2 \\ &\overset{(S13)}{\approx} (\mathbf{X}_1\mathbf{b})^\mathsf{T} \left(\frac{1}{\lambda^*}\mathbf{U}_2\mathbf{U}_1^\mathsf{T}\right)^\mathsf{T} (\mathbf{X}_2\mathbf{b}) \\ &= \frac{1}{\lambda^*}\mathbf{b}^\mathsf{T}\mathbf{X}_1^\mathsf{T}\mathbf{U}_1\mathbf{U}_2^\mathsf{T}\mathbf{X}_2\mathbf{b}. \end{aligned} \tag{S14}$$

which converges to zero as $\lambda^*$ approaches infinity:

$$\lim_{\lambda^* \to \infty} \mathbb{E}\left[\mathsf{corr}\left(\hat{\mathbf{y}}_{2,\mathbf{U}},\, \mathbf{y}_2\right)\right] = 0. \tag{S15}$$

That is, the expected prediction accuracy of the null model is null.

#### S1.3 Repetition of stimulus

So far, we assumed that the three partitions (i.e., training, test, and validation sets) are independent of each other, with independent stimuli and independent noise, but only sharing the identical weights. However, presenting multiple repetitions of an identical, short stimulus—from tens to thousands of times—has been one of the most classical techniques in neuroscience to cancel out random noise in the data and reveal time-locked neural responses that are consistently evoked by the stimulus (e.g., event-related potential, time-locked BOLD response). More recently, for investigating the representation of naturalistic stimuli, a design to present an identical set of stimuli to multiple participants has been popularized in order to reveal stimulus-driven responses in terms of inter-subject correlation (Hasson et al., 2004) or to find a common functional coordinate via hyperalignment (Haxby et al., 2020).

A problem occurs when the encoding analysis is naïvely applied to such data. To illustrate the point, let us consider an ideal design to reveal such a time-locked response, where the underlying signal is identical across partitions but the noise is independent: $\mathbf{s}_1 = \mathbf{s}_2 = \mathbf{s}_3$ but $\mathbf{e}_1 \neq \mathbf{e}_2 \neq \mathbf{e}_3$. Then, our projection matrix with $\lambda^* \approx 0$ will be:

$$\mathbf{P}_\mathbf{X} = \mathbf{X}_1 \left(\mathbf{X}_1^\mathsf{T}\mathbf{X}_1 + \mathbf{L}^*\right)^{-1} \mathbf{X}_1^\mathsf{T} \approx \mathbf{X}_1 \left(\mathbf{X}_1^\mathsf{T}\mathbf{X}_1\right)^{-1} \mathbf{X}_1^\mathsf{T}, \tag{S16}$$

when $\mathbf{X}_1^\mathsf{T}\mathbf{X}_1$ is invertible. This simplifies the expected prediction accuracy to

$$\begin{aligned}\mathbb{E}\left[\text{corr}\left(\hat{\mathbf{y}}_{2,\mathbf{X}},\, \mathbf{y}_2\right)\right] \propto \mathbf{s}_1^\mathsf{T}\mathbf{P}_\mathbf{X}^\mathsf{T}\mathbf{s}_1 &\overset{(S16)}{\approx} (\mathbf{X}_1\mathbf{b})^\mathsf{T}\mathbf{X}_1 \left(\mathbf{X}_1^\mathsf{T}\mathbf{X}_1\right)^{-1} \mathbf{X}_1^\mathsf{T}(\mathbf{X}_1\mathbf{b}) \\ &= \mathbf{b}^\mathsf{T}\mathbf{X}_1^\mathsf{T}\mathbf{X}_1 \left(\mathbf{X}_1^\mathsf{T}\mathbf{X}_1\right)^{-1} \mathbf{X}_1^\mathsf{T}\mathbf{X}_1\mathbf{b} \\ &= \mathbf{b}^\mathsf{T}\mathbf{X}_1^\mathsf{T}\mathbf{X}_1\mathbf{b} = \|\mathbf{s}_1\| > 0.\end{aligned} \tag{S17}$$

Equivalently, Equation S17 being positive can be shown based on that the covariance matrix $\mathbf{X}_1^\mathsf{T}\mathbf{X}_1$ is symmetric and that $\mathbf{X}_1$ is a rectangular matrix with independent columns, which means $\mathbf{X}_1^\mathsf{T}\mathbf{X}_1$ is positive definite. By definition, $\mathbf{b}^\mathsf{T}\mathbf{X}_1^\mathsf{T}\mathbf{X}_1\mathbf{b} > 0$ for $\mathbf{b} \neq \mathbf{0}$. This leads to a rather unsurprising conclusion—with a sufficiently strong signal, the expected prediction accuracy will be positive, also with the repeated signals[2].

In the case of the null model, however, a repeated strong signal (i.e., $\|\mathbf{s}_1\| = \|\mathbf{s}_2\| = \|\mathbf{s}_3\| \gg 0$), regardless of the null features, can alter the optimisation process, disabling proper regularisation. In this

[2] Notably, here the expected prediction accuracy is proportional to the square of the signal in the training set, rather than the test set (Equation S9). This may affect generalisation performance in hold-out validation (where the performance is evaluated only once on a separate test set), but in cross-validation the performance will be averaged out across sets. Either way, repetition of stimulus across sets leads to a spurious inflation of performance estimation.

case, the optimisation prediction accuracy for a given $\lambda \geq 0$ is:

$$\begin{aligned}
\mathbb{E}\left[\text{corr}\left(\hat{\mathbf{y}}_{3,\mathbf{X}}[\lambda],\, \mathbf{y}_3\right)\right] = \mathbb{E}\left[\frac{\left(\hat{\mathbf{y}}_{3,\mathbf{X}}[\lambda]\right)^\mathsf{T}\mathbf{y}_3}{\|\hat{\mathbf{y}}_{3,\mathbf{X}}\|\|\mathbf{y}_3\|}\right] &\propto \mathbb{E}\left[\left(\hat{\mathbf{y}}_{3,\mathbf{X}}[\lambda]\right)^\mathsf{T}\mathbf{y}_3\right] = \mathbb{E}\left[\mathbf{y}_1^\mathsf{T}\mathbf{P}_\mathbf{X}^\mathsf{T}[\lambda]\mathbf{y}_3\right] \\
&= \mathbb{E}\left[(\mathbf{s}_1+\mathbf{e}_1)^\mathsf{T}\mathbf{P}_\mathbf{X}^\mathsf{T}[\lambda](\mathbf{s}_1+\mathbf{e}_3)\right] \\
&= \mathbf{s}_1^\mathsf{T}\left\{\mathbf{X}_1\left(\mathbf{X}_1^\mathsf{T}\mathbf{X}_1+\lambda\mathbf{I}\right)^{-1}\mathbf{X}_1^\mathsf{T}\right\}^\mathsf{T}\mathbf{s}_1
\end{aligned} \tag{S18}$$

If $\mathbf{X} \in \mathbb{R}^{T\times P}$ where $P = FD$ is orthonormal and full-rank, Equation S18 can be simplified as a quadratic form of a scaled Ordinary Least Squares projection matrix and the signal as (Hastie et al., 2009, pp. 64):

$$\mathbf{s}_1^\mathsf{T}\left\{\mathbf{X}_1\left(\mathbf{X}_1^\mathsf{T}\mathbf{X}_1+\lambda\mathbf{I}\right)^{-1}\mathbf{X}_1^\mathsf{T}\right\}^\mathsf{T}\mathbf{s}_1 = \frac{1}{1+\lambda}\mathbf{s}_1^\mathsf{T}\mathbf{X}_1\left(\mathbf{X}_1^\mathsf{T}\mathbf{X}_1\right)^{-1}\mathbf{X}_1^\mathsf{T}\mathbf{s}_1 = \frac{\|\mathbf{s}_1\|}{1+\lambda}, \tag{S19}$$

for which the optimal $\lambda$ to maximize the prediction accuracy is zero ($\because \|\mathbf{s}_1\| > 0$):

$$\lambda^* = \arg\max_\lambda \frac{\|\mathbf{s}_1\|}{1+\lambda} = 0. \tag{S20}$$

More generally, the singular value decomposition of $\mathbf{X}$ can be used; $\mathbf{X} = \mathbf{U}\mathbf{D}\mathbf{V}^\mathsf{T}$ where $\mathbf{U} \in \mathbb{R}^{T\times P}$ and $\mathbf{V} \in \mathbb{R}^{P\times P}$ are orthonormal, and the diagonal matrix $\mathbf{D} \in \mathbb{R}^{P\times P}$ contains the singular values: $d_1 \geq d_2 \geq \cdots \geq d_P \geq 0$. Then, Equation S18 can be written as (Hastie et al., 2009, Eq. 3.47):

$$\mathbf{s}_1^\mathsf{T}\left\{\mathbf{X}_1\left(\mathbf{X}_1^\mathsf{T}\mathbf{X}_1+\lambda\mathbf{I}\right)^{-1}\mathbf{X}_1^\mathsf{T}\right\}^\mathsf{T}\mathbf{s}_1 = \mathbf{s}_1^\mathsf{T}\left(\sum_{j=1}^{P}\mathbf{u}_{(j)}\frac{d_j^2}{d_j^2+\lambda}\mathbf{u}_{(j)}^\mathsf{T}\right)\mathbf{s}_1. \tag{S21}$$

For any $d_j = 0$ (i.e., $\mathbf{X}$ is rank-deficit), $\lambda$ needs to be positive for the prediction accuracy value to be defined. Nonetheless, the prediction accuracy is maximized when $\lambda^* = \epsilon \approx 0$ for $\epsilon$ is the smallest positive value.

When unregularised (i.e., $\mathbf{L}^* \approx \mathbf{0}$), the projection matrix can be approximated as

$$\mathbf{P}_\mathbf{U} = \mathbf{U}_1\left(\mathbf{U}_1^\mathsf{T}\mathbf{U}_1+\mathbf{L}^*\right)^{-1}\mathbf{U}_1^\mathsf{T} \approx \mathbf{U}_1\left(\mathbf{U}_1^\mathsf{T}\mathbf{U}_1\right)^{-1}\mathbf{U}_1^\mathsf{T}, \tag{S22}$$

Thus, similarly to Equation S17, the expected null prediction accuracy of the Red Team can be approximated:

$$\mathbb{E}\left[\text{corr}\left(\hat{\mathbf{y}}_{2,\mathbf{U}},\, \mathbf{y}_2\right)\right] \propto \mathbf{s}_1^\mathsf{T}\mathbf{P}_\mathbf{U}^\mathsf{T}\mathbf{s}_1 \overset{(S22)}{\approx} \mathbf{s}_1^\mathsf{T}\mathbf{U}_1\left(\mathbf{U}_1^\mathsf{T}\mathbf{U}_1\right)^{-1}\mathbf{U}_1^\mathsf{T}\mathbf{s}_1. \tag{S23}$$

$\mathbf{U}_1^\mathsf{T}\mathbf{U}_1$ is positive definite due to its symmetry and independence of the columns in $\mathbf{U}_1$. Since the inverse operation preserves the signs of eigenvalues of a square matrix, its inversion $\left(\mathbf{U}_1^\mathsf{T}\mathbf{U}_1\right)^{-1}$ is also positive definite. A substitution ($\mathbf{d} \equiv \mathbf{U}_1^\mathsf{T}\mathbf{s}_1 \neq \mathbf{0}$) can clarify that Equation S23 is positive by definition:

$$\mathbf{s}_1^\mathsf{T}\mathbf{P}_\mathbf{U}^\mathsf{T}\mathbf{s}_1 \approx \mathbf{d}^\mathsf{T}\left(\mathbf{U}_1^\mathsf{T}\mathbf{U}_1\right)^{-1}\mathbf{d} > 0. \tag{S24}$$

This may surprise some readers; however, Equation S24 implies that the null prediction of the Red Team is expected to be greater than zero. Depending on the signal-to-noise ratio, this may result in Type-I (false positive) errors. This is due to the circular fallacy introduced by the *leakage in training examples*, i.e., SDL.

### S2 Supplementary Results: Simulation

#### S2.1 Simulation methods

##### S2.1.1 Predictors

True features were sampled from an $F$-dimensional multivariate Gaussian distribution with a correlation between adjacent parameters as: $\mathbf{x} \sim \mathcal{N}_F(\mathbf{0}, \mathbf{\Sigma}_X)$ where the covariance is defined as:

$$\mathbf{\Sigma}_X = \begin{bmatrix} 1 & \rho_X & \rho_X^2 & \cdots & \rho_X^{F-1} \\ \rho_X & 1 & \rho_X & \cdots & \rho_X^{F-2} \\ \rho_X^2 & \rho_X & 1 & \cdots & \rho_X^{F-3} \\ \vdots & \vdots & \vdots & \ddots & \vdots \\ \rho_X^{F-1} & \rho_X^{F-2} & \rho_X^{F-3} & \cdots & 1 \end{bmatrix}. \tag{S25}$$

The $f$-th feature $\mathbf{x}^{(f)} = [x_1^{(f)}, x_2^{(f)}, \dots, x_T^{(f)}]^\mathsf{T}$ is given with an AR(1) temporal autocorrelation $\phi_X$ as:

$$x_t^{(f)} = x_t^{(f)} + \phi_X x_{t-1}^{(f)}. \tag{S26}$$

Null features $\mathbf{u}$ were created in the same way as $\mathbf{x}$, but independently. Only causal (i.e., nonnegative) delays were considered from $\{0\}$ to $\{0, \dots, D-1\}$ in creating a design matrix by horizontally concatenating

Toeplitz matrices as:

$$\mathbf{X} = \begin{bmatrix} x_1^{(1)} & \cdots & x_{1-(D-1)}^{(1)} & x_1^{(2)} & \cdots & x_{1-(D-1)}^{(2)} & \cdots & x_1^{(F)} & \cdots & x_{1-(D-1)}^{(F)} \\ \vdots & \ddots & \vdots & \vdots & \ddots & \vdots & \cdots & \vdots & \ddots & \vdots \\ x_T^{(1)} & \cdots & x_{T-(D-1)}^{(1)} & x_T^{(2)} & \cdots & x_{T-(D-1)}^{(2)} & \cdots & x_T^{(F)} & \cdots & x_{T-(D-1)}^{(F)} \end{bmatrix}. \tag{S27}$$

For convenience, all predictors were standardised to zero-mean and unit variance.

##### S2.1.2 Responses

Responses $\mathbf{Y}_i \in \mathbb{R}^{T \times V}$ were generated by plugging in a design matrix $\mathbf{X}_i$, true weights $\mathbf{B} \in \mathbb{R}^{FD \times V}$, and noise $\mathbf{E}_i \in \mathbb{R}^{T \times V}$ to the FIR model Equation S1 for the $i$-th partition as $\mathbf{Y}_i = \mathbf{X}_i\mathbf{B} + \mathbf{E}_i$ where $\mathbf{Y}_i = \left[\mathbf{y}_i^{(1)}, \mathbf{y}_i^{(2)}, \cdots, \mathbf{y}_i^{(V)}\right]$, $\mathbf{B} = \left[\mathbf{b}^{(1)}, \mathbf{b}^{(2)}, \cdots, \mathbf{b}^{(V)}\right]^\mathsf{T}$, and $\mathbf{E}_i = \left[\mathbf{e}_i^{(1)}, \mathbf{e}_i^{(2)}, \cdots, \mathbf{e}_i^{(V)}\right]$.

True weights of the $v$-th variate $\mathbf{b}^{(v)}$ (a column of a response variable) were created from the multivariate Gaussian distribution with an AR(1) temporal correlation $\phi_B$ along the delays and a covariance $\sigma_b$ across predictors corresponding to the rows of the design matrix as:

$$\mathbf{b}^{(v)} = \left[b_{1,1}^{(v)}, \cdots, b_{1,D}^{(v)}, b_{2,1}^{(v)}, \cdots, b_{2,D}^{(v)}, \cdots, b_{F,1}^{(v)}, \cdots, b_{F,D}^{(v)}\right]^\mathsf{T} \tag{S28}$$

where $b_{f,d}^{(v)}$ is a scalar coefficient for the $v$-th variate, the $f$-th feature, and the $d$-th delay. Noise $\mathbf{e}^{(v)}$ at the $v$-th variate was also created as Gaussian noise with an AR(1) temporal correlation $\phi_E$ as $e_t^{(v)} = e_t^{(v)} + \phi_E e_{t-1}^{(v)}$. In case of multivariate models ($V \geq 2$), weights and noise have spatial autocorrelation: $\mathbf{b}^{(v)} = \mathbf{b}^{(v)} + \theta_B \mathbf{b}^{(v-1)} + \theta_B \mathbf{b}^{(v+1)}$ and $\mathbf{e}^{(v)} = \mathbf{e}^{(v)} + \theta_E \mathbf{e}^{(v-1)} + \theta_E \mathbf{e}^{(v+1)}$ where a variate $v$ is a neighbour of $v-1$ and $v+1$ except for boundaries ($v = 1$ and $v = V$).

Before summing the signal $\mathbf{XB}$ and error $\mathbf{E}$, the mean variance of error across variates was scaled so that it achieved the intended signal-to-noise ratio as $S = 10 \log_{10} \sigma_S^2 / \sigma_E^2$. Then, all response variates were standardised to zero-mean and unit-variance.

##### S2.1.3 optimisation and evaluation

Ridge parameters were optimised via a grid search ($\lambda$-grid $= 10^{[-10,-9,\ldots,10]}$) for each variate independently. To implement a nested 4-by-3-fold cross-validation (CV), in total 4 "trials" were generated for each random sampling. For the 4-fold-outer-loop, 3 trials (outer-training) vs. 1 trial (outer-test) were partitioned. Then, within each 3-fold-inner-loop with the 3 trials, 2 trials (inner-training) vs. 1 trial (inner-validation) were partitioned. In total 12 different partitions (each was called a "CV-fold") were used for training (50%), validation (25%), and test (25%) models. Prediction accuracy of Pearson correlation was averaged across the CV-folds. Random sampling was repeated for 1000 times for each combination of model parameters.

###### S2.1.4 Computational considerations

To minimise the number of inversion operations, a general linear model (GLM) formulation was used. That is, for each possible $\lambda$, a regularised covariance matrix was inverted only once for all variates (i.e., voxels or channels). The sum of squared errors was then temporarily stored, allowing the optimal $\lambda$ to be determined for each variate. This approach is more efficient than a naïve method of inverting the covariance matrix separately for each variate, which would redundantly repeat the same calculation for all variates. Also, when computing Predictions, variates with the same optimal $\lambda$ were grouped together into GLM models to reduce the number of inversions.

Computation was carried out using an in-house high-performance computing (HPC) server, where a user is allowed to utilise up to 192 CPUs of Intel Xeon Gold 6130 [2.10 GHz] in parallel. The actual utilisation varied between 32–192 CPUs, depending on the demands of other users. An individual job of 1,000 random samplings took 100–400 seconds of CPU time. A total of 45,899 jobs, which amounted to approximately 7 months of CPU time, was completed in about one week on the HPC server.

##### S2.2 Simulation examples

Here, I show the details of the simulations using a simple exemplar case (i.e., a toy example). The distributions of Pearson correlation and optimal ridge penalty are shown in the main text. In this example, two features with a high temporal autocorrelation were generated without repetitions (Figure S1**a**) and with repetitions (Figure S2**a**). The null features were created as independent Gaussian noise (Figure S1**m**). Thus, all coefficients were highly regularized for the null features (Figure S1**r**; pink). Consequently, the null predictions were mostly flat (Figure S1**o,p**; dotted lines) and expectedly, the null prediction accuracies were around $r = 0$ (Figure S1**l**; pink).

However, with the stimulus repeated across sets (i.e., identical signals in the training and test sets), the regularization for the null features was much smaller—almost close to the true features (Figure S2**r**; pink vs. lime green)—and the null prediction accuracies were around $r = 0.5$ (Figure S2**l**; pink). That is, due to the repetition of the stimulus, even with independent noise (Figure S2**e,k**), the prediction accuracies

Table 2: Parameters of the simulations. Number of time points $T = 100$, $\lambda$-grid $= 10^{[-10,-9,\ldots,15]}$, Number of samplings $K = 1000$.

| Category | Parameter | Notation |
|---|---|---|
| Complexity of model | Number of delays | $D$ |
| | Number of features | $F$ |
| Dimensionality of data | Number of variates | $V$ |
| Strength of signal | Signal-to-noise ratio (SNR) | $S$ |
| AR(1) temporal autocorrelation (if $T \geq 2$) | of true predictors | $\phi_X$ |
| | of null predictors | $\phi_U$ |
| | of true weight | $\phi_B$ |
| | of noise | $\phi_E$ |
| AR(1) spatial autocorrelation (if $V \geq 2$) | of true weight | $\theta_B$ |
| | of noise | $\theta_E$ |
| Multicollinearity (if $F \geq 2$) | of true predictors | $\rho_X$ |
| | of null predictors | $\rho_U$ |
| Presence of information leakage | Binary flag for the stimulus repetition | `IsRep` |

based on the null features were falsely inflated.

#### S2.3 Simulation results

In this section, I highlight major factors that worsen the Type-I error due to SDL. To keep the number of combinations manageable, univariate models ($F = 1$, $V = 1$) were first considered. Then, while iteratively pruning out irrelevant factors, models with multivariate features ($F$) and multivariate responses ($V$) were considered. Methodological details of the simulation are described in the Supplementary Materials. The parameters of simulation are summarised in Table 2.

##### S2.3.1 Univariate-feature, univariate-response

The first batch of simulations was restricted to a univariate feature (the number of features $F = 1$) and a univariate response (the number of variates $V = 1$). The explored parameter levels were: $D \in \{1, 3, 5, 7, 9, 11\}$, $S \in \{-10, 0, 10\}$, $\phi_X \in \{0, 0.5, 1\}$, $\phi_U \in \{0, 0.5, 1\}$, $\phi_E \in \{0, 0.5, 1\}$, $\phi_B \in \{0, 0.5, 1\}$, $\texttt{IsRep} \in \{0, 1\}$. With these parameter levels, total 2,916 combinations were created, each sampled 1,000 times.

To illustrate the most distinctive effects, prediction accuracies averaged across 1,000 random sampling are shown in Figure S3. Expectedly, the SNR increased the prediction accuracy based on the true

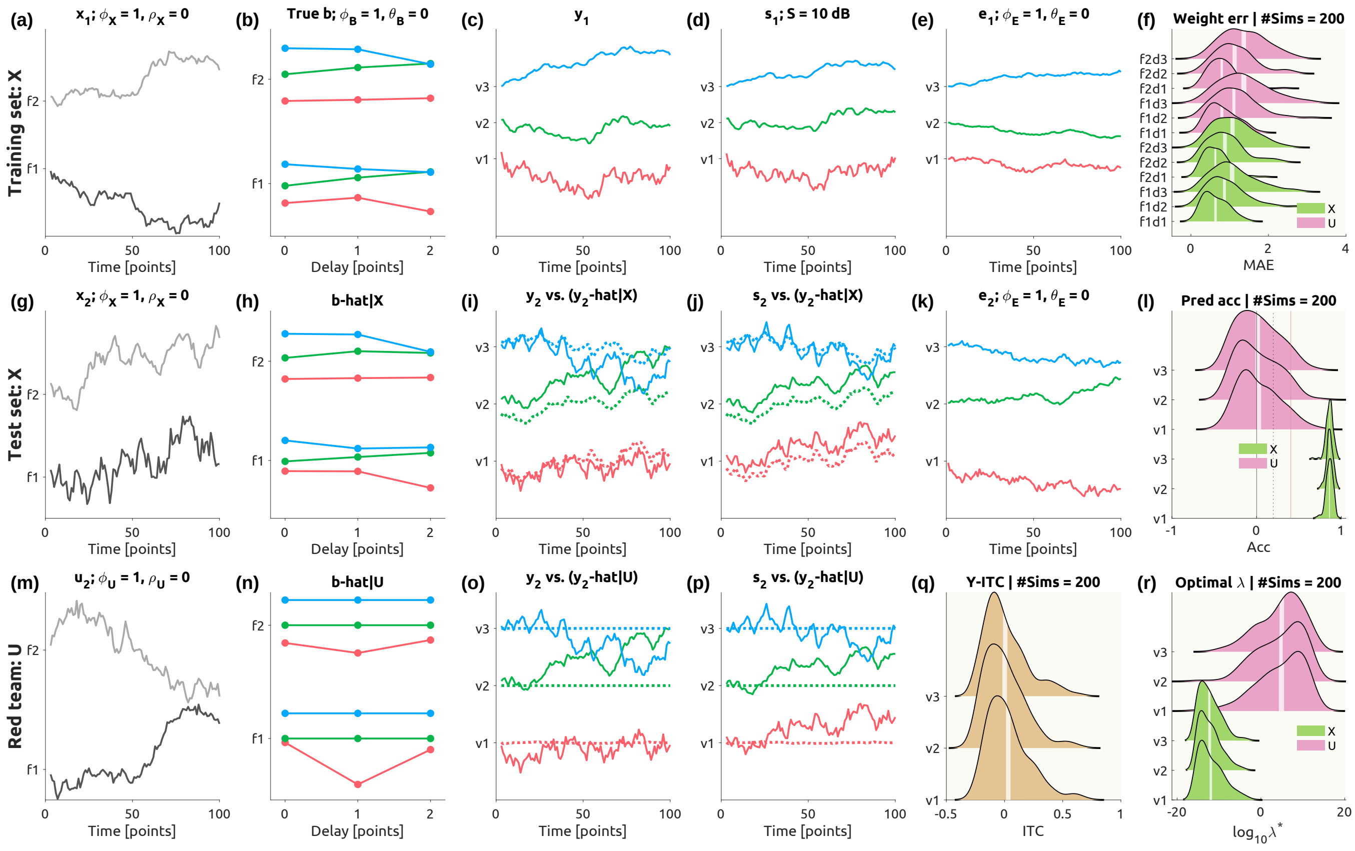


Figure S1: A toy example without stimulus repetition. Apart from the right-most panels in pale beige **(f, l, r)**, the panels in the first row **(a–e)** correspond to the training set with the true features $\mathbf{X}_1$, the panels in the second row **(g–k)** correspond to the test set with the true features $\mathbf{X}_2$, and the panels in the third row **(m–p)** correspond to the Red Team's null features $\mathbf{U}_2$. The panels in the first column **(a, g, m)** show the true features $\mathbf{x}$ or null features $\mathbf{u}$ in gray scale. The panels in the second column show the true weights $\mathbf{b}$ **(b)** or estimation based on the true features **(h)** or null features **(n)** where the RGB color represents one of three response variates. The panels in the third column **(c, i, o)** show the response $\mathbf{y}$ (solid lines) and prediction based on true or null features $\mathbf{y}$-hat (dashed lines), and the panels in fourth column **(d, j, p)** show the true signal $\mathbf{s}$ (solid lines) and the prediction (dashed lines). The two panels in the fifth column **(e, k)** show true noise $\mathbf{e}$. In the pale beige panels **(f, l, q, r)**, distributions from 200 simulations are shown in ridgeline plots with the 95% confidence interval of the mean shown in white strips: **(f)** mean absolute error (MAE) of weight estimation based on the true features ($\mathbf{X}$; lime green) or null features ($\mathbf{U}$; pink), **(l)** prediction accuracies in Pearson's correlation coefficients with a black vertical line for an absolute zero, a gray vertical line for an uncorrected $P < 0.05$ (assuming independent time points), and a red vertical line for a Bonferroni-corrected $P < 0.05$ adjusted for the number of variates. **(q)** inter-trial correlation (ITC) of the responses (brown), **(r)** exponents of the geometrically averaged optimal $\lambda$'s. Parameters to generate this simulation set are: $T = 100, V = 3, F = 2, D = 3, S = 1\text{dB}, \phi_X = 1, \rho_X = 0, \phi_B = 1, \theta = 0, \phi_E = 1, \theta_E = 0, \texttt{IsRep} = 0$. See Table 2 for the explanation of parameters.

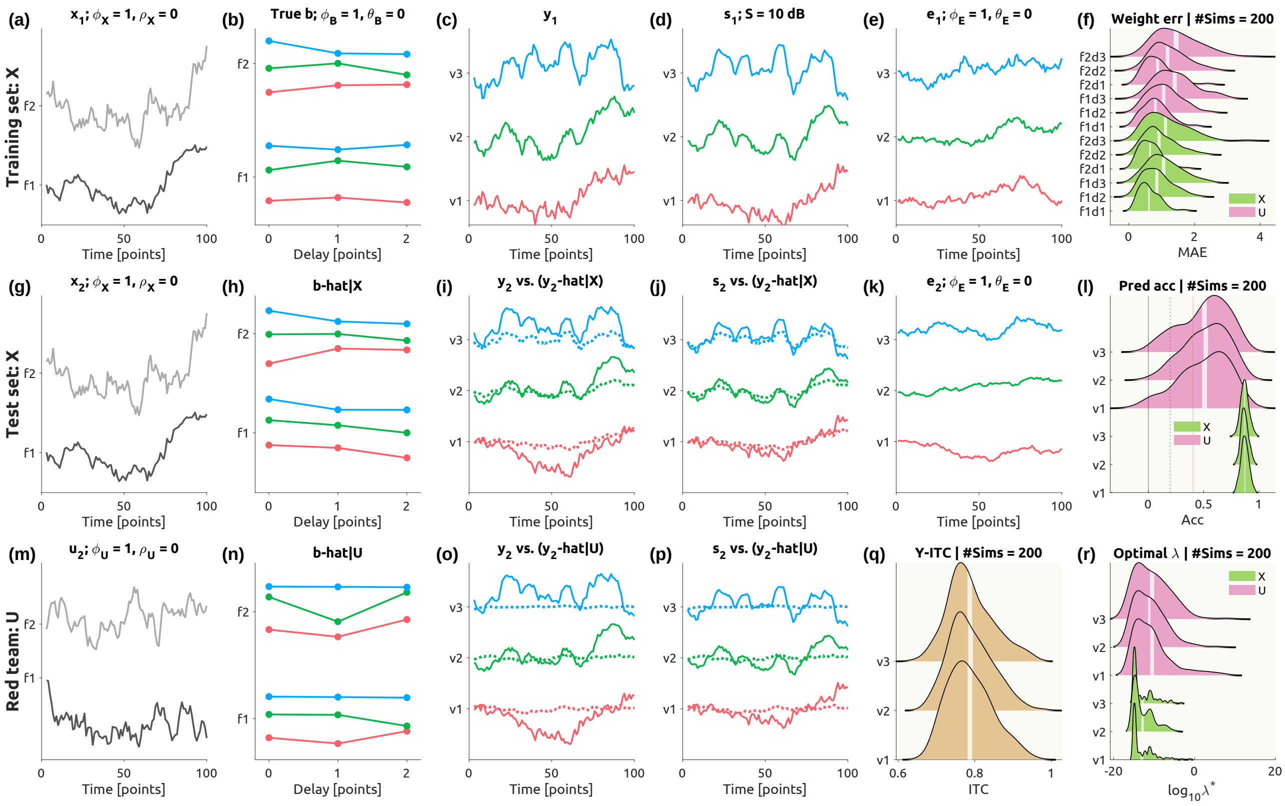


Figure S2: A toy example with stimulus repetition. The visualization scheme is identical to Figure S1. Parameters to generate this simulation set are identical to those in Figure S1, except $\texttt{IsRep} = 1$.

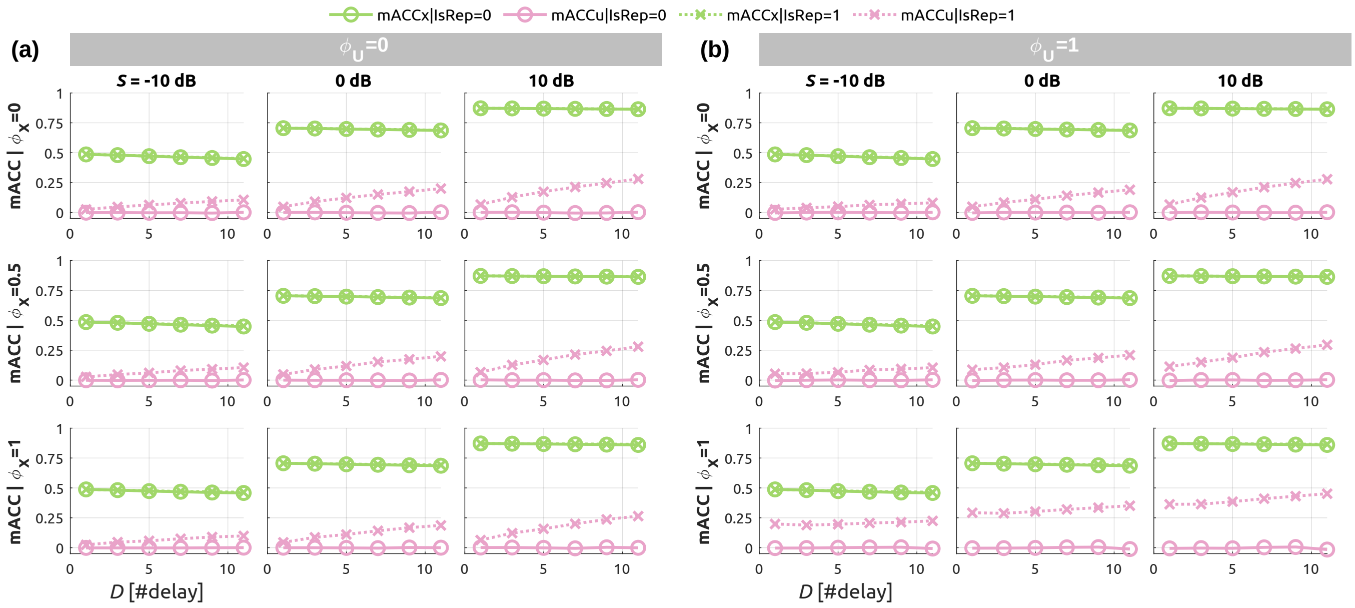


Figure S3: Simulations of univariate-feature, univariate-response models. Mean prediction accuracies are plotted over the number of delays $D$ when the temporal autocorrelation **(a)** $\phi_U = 0$ and **(b)** $\phi_U = 1$. Each marker corresponds to the averaged Pearson correlation coefficients of 1000 simulations. Predictions based on true features ($\mathbf{X}$) are shown in lime green and null features ($\mathbf{U}$) in pink. Open circles with solid lines indicate accuracies when the true signals were not repeated. Crosses with dashed lines represent cases where the true signals were repeated. Each column corresponds to a specified SNR level. $\phi_X = 0$ in the top row and $\phi_X = 1$ in the bottom row. Other parameters were as follows: $K = 1000$, $\phi_B = 0$, $\phi_E = 0$. Other combinations of parameters showed generally similar patterns. See Table 3 and Table 4 for effect sizes.

features (green markers). The prediction based on the null features remained zero when the stimuli were independent across the cross-validation (CV) partitions (pink circles). However, when the stimuli were identical across the partitions, the null prediction accuracy increased over the number of delays and the SNR (pink crosses). In particular, the effect of temporal autocorrelation $\phi_X$ when $\phi_U = 1$ substantially increased the null prediction with the repeated stimuli.

To comprehensively assess the possible effects, a full factorial model on mean prediction accuracies was fitted with all seven variables as in Wilkinson notation:

$$r \sim 1 + D * S * \phi_X * \phi_U * \phi_E * \phi_B * \texttt{IsRep}, \tag{S29}$$

where $r$ is the mean prediction accuracy averaged for each set of 1000 simulations either based on true or null predictors, $1$ represents an intercept, and $*$ denotes factor crossing as in $a * b = a + b + a : b$ with : denoting an interaction. This yielded linear models with 128 terms from the intercept to the seven-way

Table 3: Strong effects ($\eta_p^2 \geq 0.160$) on the prediction accuracies in univariate models with univariate responses using true predictors $\mathbf{X}$. Full anova table: https://zenodo.org/records/15101528/files/uni-uni_anova-x.xlsx.

| Contrast | SS | $F$-stat | $\eta_p^2$ |
|---|---|---|---|
| $S$ | 8.377e+01 | 1.867e+05 | 0.985 |
| $\phi_X$:$\phi_E$ | 2.645e-01 | 5.895e+02 | 0.175 |
| $S$:$\phi_E$ | 2.037e-01 | 4.539e+02 | 0.140 |
| $S$:$\phi_X$:$\phi_E$ | 3.187e-01 | 7.101e+02 | 0.203 |

interaction, 2,916 observations, and adjusted $R^2 = 0.985$ for $\mathbf{X}$ and $0.957$ for $\mathbf{U}$, which are reasonable given that the accuracy metrics are averaged within each combination of parameters. Due to the large number of observations, $P$-values were not highly selective ($P_{\text{Bonferroni}} < 0.0001$ for many of contrasts; 9 for $\mathbf{X}$, 27 for $\mathbf{U}$ out of 127). Thus, on top of $P_{\text{Bonferroni}} < 0.0001$, only effects with moderate effect sizes ($\eta_p^2 \geq 0.16$) were considered for further discussion (Table 3, Table 4).

As expected, the SNR consistently increased the prediction accuracy ($\eta_p^2[S] = 0.985$ for $\mathbf{X}$; $\eta_p^2[S] = 0.660$ for $\mathbf{U}$). Most importantly, the interaction of the stimulus repetition (`IsRep`) with the signal strength (SNR, $S$), and the autocorrelation of the true and null features ($\phi_X$, $\phi_U$) were markedly found on the prediction accuracy based on the null features ($\max \eta_p^2 = 0.661$, Table 4). Put differently, the SDL effect (i.e., a false inflation of the null prediction accuracy by the repetition of stimulus across CV partitions) was most pronounced when the true underlying signal was strong and the true features were temporally autocorrelated. This finding is consistent with Equation S17 above, showing that the SDL effect depends on the nonzero strength of the underlying signal. The clearer the signal, the more likely the SDL effect will occur when analysed in a circular CV design.

In addition, the SDL effect strongly interacted with the complexity of the underlying (and fitted) models (i.e., `IsRep` : $D$; Table 4). While in a correct CV design (`IsRep` $= 0$), the model complexity did not increase the prediction accuracy as this was regularised by independent optimisation and evaluation. However, in a circular CV design (`IsRep` $= 1$), the model complexity increased the prediction accuracy (hence the SDL effect) since the model was fitted to the identical signal across CV partitions. Naturally, this effect further interacted with the strength of the signal (`IsRep` : $D$ : $S$; Table 4).

The effect of the autocorrelation of the weight time series $\phi_B$ was found to be negligible for both $\mathbf{X}$ and $\mathbf{U}$ ($\eta_p^2 < 0.009$). Guided by these results, we fixed the autocorrelation of the weight time series to zero ($\phi_B = 0$) in the following simulations.

Table 4: Strong effects ($\eta_p^2 \geq 0.160$) on the prediction accuracies in univariate models with univariate responses using null predictors $\mathbf{U}$. Full anova table: https://zenodo.org/records/15101528/files/uni-uni_anova-u.xlsx.

| Contrast | SS | $F$-stat | $\eta_p^2$ |
|---|---|---|---|
| IsRep | 1.569e+01 | 3.864e+04 | 0.933 |
| $S$ | 2.199e+00 | 5.413e+03 | 0.660 |
| $D$ | 1.504e+00 | 3.704e+03 | 0.571 |
| $\phi_X$ | 4.312e-01 | 1.062e+03 | 0.276 |
| $\phi_U$ | 3.607e-01 | 8.881e+02 | 0.242 |
| $S$:IsRep | 2.209e+00 | 5.438e+03 | 0.661 |
| IsRep:$D$ | 1.458e+00 | 3.589e+03 | 0.563 |
| $\phi_X$:$\phi_U$ | 6.987e-01 | 1.720e+03 | 0.382 |
| $\phi_X$:IsRep | 4.435e-01 | 1.092e+03 | 0.281 |
| $\phi_U$:IsRep | 3.357e-01 | 8.266e+02 | 0.229 |
| $S$:$D$ | 2.164e-01 | 5.328e+02 | 0.160 |
| $\phi_X$:$\phi_U$:IsRep | 7.277e-01 | 1.792e+03 | 0.391 |
| $S$:IsRep:$D$ | 2.408e-01 | 5.929e+02 | 0.175 |

##### S2.3.2 Multivariate-feature, univariate-response

Since most often we are interested in multivariate features (e.g., motion energies, spectrograms, deep ANN embeddings), it is of interest how the dimensionality and multicollinearity of features influence SDL artifact.

The explored parameter levels were: $D \in \{1, 5, 9\}$, $F \in \{5, 10, 15, 20\}$, $S \in \{-10, 0, 10\}$, $\phi_X \in \{0, 0.5, 1\}$, $\phi_U \in \{0, 0.5, 1\}$, $\phi_E \in \{0, 0.5, 1\}$, $\phi_B = 0$, $\rho_X \in \{0, 0.5, 1\}$, $\rho_U \in \{0, 0.5, 1\}$, IsRep $\in \{0, 1\}$. With these levels, the full combinations amounted to 17,496, for each of which, once more, 1,000 random samplings were carried out.

Figure S4 displays the simulated effects. The number of features $F$ reduced the true models' prediction accuracies without repetitions (green solid lines), and more so with more complex response functions (e.g., $D = 9$). In contrast, increasing the number of features led to higher prediction accuracies in null models with stimulus repetition (pink dashed lines). Moreover, at a higher SNR (e.g., 10 dB), the null prediction accuracies (pink dashed lines) were even higher than the true prediction accuracies (green solid lines) with many independent predictors (e.g., 9 delays $\times$ $\geq$15 features when $\rho_X = 0$; Figure S4**a**). This suggests that, in a bad combination, the null prediction accuracy can go beyond the noise ceiling (i.e., green solid lines; assuming we know the true predictors), which is the plausibly highest prediction accuracy bounded by the noise level of the signal. This crossing of the true and null prediction accuracies was attenuated when features were highly correlated ($\rho_X = 1$; Figure S4**b**), which reduced the effective degrees of freedom.

To quantify the observed effects, once again, a full factorial model on mean prediction accuracies was fitted with all nine variables:

$$r \sim 1 + D * F * S * \phi_X * \phi_U * \phi_E * \rho_X * \rho_U * \texttt{IsRep}. \tag{S30}$$

A full factorial model with 17,496 observations and 511 terms resulted in adjusted $R^2 = 0.975$ for $\mathbf{X}$ and $0.923$ for $\mathbf{U}$ (Table 5, Table 6). A strong effect of the number of features $F$ was found for $\mathbf{X}$ ($\eta_p^2 = 0.276$) but only an intermediate effect for $\mathbf{U}$ ($\eta_p^2 = 0.102$). Additionally, its interaction with stimulus repetition (IsRep) for the null features was only intermediate $\mathbf{U}$ ($\eta_p^2 = 0.101$), suggesting the dimension of the features alone was not a strong factor for the SDL effect.

However, the interaction between the stimulus repetition and the multicollinearity of the null features ($\rho_U$ : IsRep) was strong ($\eta_p^2 = 0.442$; Table 6), where the SDL effect was more pronounced when the null features exhibited lower autocorrelation (see Supplementary Figure S1). That is, when the null features have greater effective degrees of freedom (i.e., lower autocorrelation), the model could more flexibly fit the repeated signal, inflating the SDL effect.

In addition, an interesting finding was that the true prediction accuracy decreased as the number of features $F$ and the number of delays $D$ increased. This was due to the limited number of samples ($T = 100$) as compared to the high dimensionality of the feature space, which made the linear model ill-posed. While regularisation makes fitting feasible, the inherent limitation exists. Put differently, these results suggest that a sufficient number of samples is required to faithfully estimate the transfer function of the high-dimensional feature space.

##### S2.3.3 Multivariate-feature, multivariate-response

Finally, it was tested whether the dimensionality and spatial autocorrelation of responses (variates) affect the SDL artifact. First, it is worth noting that the encoding model is typically a univariate-response model (e.g., "voxel-wise" or "channel-wise"). The assumption of spatially (i.e., across response units) independent noise is similar to that of the classical general linear model (Friston et al., 1994), where a diagonal covariance structure across lattice sampling grids (e.g., voxels) is assumed[3]. Likewise, the encoding model is independently optimised for each response unit in the current paper[4]. Thus, while

[3] Therefore, the term 'massive-univariate' would be more fitting than 'multivariate'.

[4] In some EEG studies, hyperparameters were averaged across response units (i.e., EEG channels). This practice introduces spatial dependency that leads to a suboptimal regularisation for individual response units.

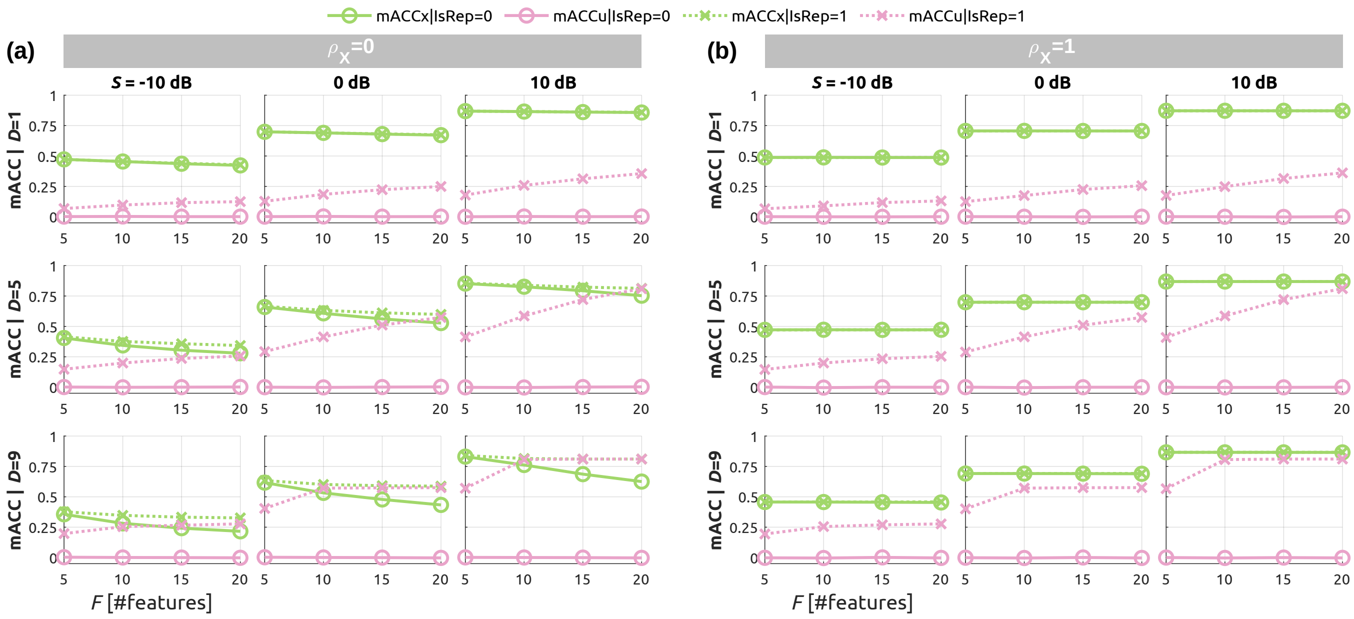


Figure S4: Simulations of multivariate-feature, univariate-response models. Mean prediction accuracies are plotted over the number of features $F$ when the multicollinearity **(a)** $\rho_X = 0$ and **(b)** $\rho_X = 1$. Marker styles and colors match to those in Figure S3. Each column represents a specified SNR level. The number of delays is $D = 1$ in the top rows and $D = 9$ in the bottom rows. Other parameters were as follows: $K = 1000$, $\phi_U = 0$, $\phi_B = 0$, $\phi_E = 0$, $\rho_U = 0$. See Table 5 and Table 6 for effect sizes. See Supplementary S1 for additional cases with $\rho_U = 0$ and $\rho_U = 1$ when $\rho_X = 0$.

Table 5: Strong effects ($\eta_p^2 \geq 0.160$) on the prediction accuracies of multivariate models with univariate responses using true predictors $\mathbf{X}$. Full ANOVA table: https://zenodo.org/records/15101528/files/mult-uni_anova-x.xlsx.

| Contrast | SS | $F$-stat | $\eta_p^2$ |
|---|---|---|---|
| $S$ | 5.729e+02 | 5.984e+05 | 0.972 |
| $\rho_X$ | 2.374e+01 | 2.479e+04 | 0.593 |
| $D$ | 1.258e+01 | 1.314e+04 | 0.436 |
| IsRep | 6.991e+00 | 7.302e+03 | 0.301 |
| $F$ | 6.196e+00 | 6.471e+03 | 0.276 |
| $\phi_X$ | 4.010e+00 | 4.188e+03 | 0.198 |
| $\phi_E$ | 3.557e+00 | 3.715e+03 | 0.179 |
| $\phi_X$:$\phi_E$ | 5.048e+00 | 5.273e+03 | 0.237 |
| $\rho_X$:$D$ | 3.213e+00 | 3.356e+03 | 0.165 |
| $\rho_X$:IsRep | 2.841e+00 | 2.967e+03 | 0.149 |

Table 6: Strong effects ($\eta_p^2 \geq 0.160$) on the prediction accuracies of multivariate models with univariate responses using null predictors $\mathbf{U}$. Full anova table: https://zenodo.org/records/15101528/files/mult-uni_anova-u.xlsx.

| Contrast | SS | $F$-stat | $\eta_p^2$ |
|---|---|---|---|
| `IsRep` | 4.161e+02 | 1.057e+05 | 0.862 |
| $S$ | 6.946e+01 | 1.765e+04 | 0.510 |
| $\rho_U$ | 5.282e+01 | 1.342e+04 | 0.441 |
| $D$ | 3.429e+01 | 8.715e+03 | 0.339 |
| $S$:`IsRep` | 6.933e+01 | 1.762e+04 | 0.509 |
| $\rho_U$:`IsRep` | 5.296e+01 | 1.346e+04 | 0.442 |
| `IsRep`:$D$ | 3.469e+01 | 8.816e+03 | 0.342 |

Table 7: Strong effects ($\eta_p^2 \geq 0.160$) on the prediction accuracies of multivariate models with multivariate responses using true predictors $\mathbf{X}$. Full ANOVA table: https://zenodo.org/records/15101528/files/mult-mult_anova-x.xlsx.

| Contrast | SS | $F$-stat | $\eta_p^2$ |
|---|---|---|---|
| $S$ | 1.161e+04 | 1.417e+07 | 0.976 |
| $\rho_X$ | 3.606e+02 | 4.404e+05 | 0.557 |
| $D$ | 2.177e+02 | 2.659e+05 | 0.432 |
| `IsRep` | 8.921e+01 | 1.090e+05 | 0.237 |
| $F$ | 8.851e+01 | 1.081e+05 | 0.236 |
| $\phi_X$ | 7.909e+01 | 9.660e+04 | 0.216 |
| $\phi_E$ | 7.119e+01 | 8.696e+04 | 0.199 |
| $\phi_X$:$\phi_E$ | 1.057e+02 | 1.292e+05 | 0.269 |
| $\rho_X$:$D$ | 5.227e+01 | 6.384e+04 | 0.154 |
| $S$:$\phi_X$:$\phi_E$ | 4.999e+01 | 6.106e+04 | 0.148 |

the spatial dependency may affect the family-wise error rates of multiple testing (since many correction methods exploit neighbouring supports), it is unexpected that the spatial dependency directly alters the unit-wise optimisation and following weight estimation. However, for completeness, simulations were carried out with following parameters: $D \in \{1, 7, 11\}$, $F \in \{5, 10, 20\}$, $V \in \{5, 10, 20\}$, $S \in \{-10, 0, 10\}$, $\phi_X \in \{0, 0.5, 1\}$, $\phi_U \in \{0, 0.5, 1\}$, $\phi_E \in \{0, 0.5, 1\}$, $\phi_B = 0$, $\theta_B \in \{0, 0.5, 1\}$, $\theta_E \in \{0, 0.5, 1\}$, $\rho_X \in \{0, 0.5, 1\}$, $\rho_U \in \{0, 0.5, 1\}$, `IsRep` $\in \{0, 1\}$. With these parameter levels, there were 354,287 total combinations, as before, each sampled 1,000 times. A full factorial model, incorporating all twelve variables, was fitted using 354,287 observations and 4,096 terms. As expected, the spatial dependency $\theta_B$ and $\theta_E$ neither showed any marked main effects nor interactions ($\eta_p^2 < 0.160$; Table 7, Table 8).

Table 8: Strong effects ($\eta_p^2 \geq 0.160$) on the prediction accuracies of multivariate models on multivariate responses with null predictors $\mathbf{U}$. Full ANOVA table: https://zenodo.org/records/15101528/files/mult-mult_anova-u.xlsx.

| Contrast | SS | $F$-stat | $\eta_p^2$ |
|---|---|---|---|
| IsRep | 7.636e+03 | 2.446e+06 | 0.875 |
| $S$ | 1.269e+03 | 4.066e+05 | 0.537 |
| $\rho_U$ | 8.979e+02 | 2.876e+05 | 0.451 |
| $D$ | 7.222e+02 | 2.313e+05 | 0.398 |
| $S$:IsRep | 1.270e+03 | 4.068e+05 | 0.538 |
| $\rho_U$:IsRep | 8.967e+02 | 2.872e+05 | 0.451 |
| IsRep:$D$ | 7.226e+02 | 2.315e+05 | 0.398 |

### S3 Supplementary Methods: Real Data

For clarity, the original studies (Kaneshiro et al., 2020; Sachs et al., 2020) focused on inter-subject synchrony in neural responses (EEG and fMRI) without specific hypotheses about the encoded features. Thus, these studies did not suffer from SDL.

#### S3.1 EEG data

**Data source** The original study investigated the electroencephalographic correlates of temporal structure and beat in Western-style Indian pop music with Hindi lyrics (i.e., Bollywood music) (Kaneshiro et al., 2020). The raw dataset of 48 healthy participants was downloaded from the Stanford Digital Repository (https://purl.stanford.edu/sd922db3535).

**Data acquisition** Scalp electrical potential data were acquired using a 128-channel Geodesic EEG System 300 (Electrical Geodesics, Inc., Oregon, USA) at a sampling rate of 1 kHz with vertex reference and electrode impedances less than 60 k$\Omega$.

**Preprocessing** The authors' preprocessed data (`CleanEEG_*`) from the repository were used with the channel location file (`GSN-HydroCel-125.sfp`). As explained in the original publication (Kaneshiro et al., 2020), the authors' preprocessing steps involved bandpass filtering (0.3–50 Hz), downsampling (125 Hz), ocular artifacts removal using independent component analysis, bad channel interpolation, average re-referencing, and epoching. In addition, based on previous EEG encoding analyses where the low-frequency bands of the EEG signal mostly carried the audio envelope encoding (Di Liberto et al., 2015, 2020), we further bandpass filtered the EEG data to

$\delta$ and $\theta$ bands (1–8 Hz) using the FIR filter in the MATLAB Signal Processing Toolbox (R2022b, RRID:SCR_001622).

**Data dimensions** The analysed EEG data were comprised of 125 channels, 32,878–33,982 time points (263.02–271.86 seconds at 125 Hz), 96 runs (48 subjects $\times$ 2 run) per stimulus with 12 stimuli (3 versions of 4 songs). Two runs with the same stimuli were averaged for each participant to increase the signal-to-noise ratio of the evoked response. In the original study (Kaneshiro et al., 2020), a participant was randomly assigned to one of 4 versions of the 4 songs (e.g., Intact-Song-A, Measure-shuffled-Song-B, Reserved-Song-C, Phase-scrambled-Song-D). The current study only analysed the 3 versions except for phase-scrambled version, which did not evoke strong responses.

#### S3.2 fMRI data

**Data source** The original study investigated the intersubject correlation in fMRI time series while participants were listening to sad music (Sachs et al., 2020). The published data include 2 sad musical pieces and 1 happy musical piece. The raw dataset of 39 healthy participants was downloaded from OpenNeuro (RRID:SCR_005031, https://openneuro.org/datasets/ds003085/versions/1.0.0).

**Data acquisition** Blood-oxygen-level-dependent (BOLD) signals were acquired using a 3-T Prisma magnetic resonance imaging system with a 32-channel head coil (Siemens Healthineers, Erlangen, Germany). Eight-fold accelerated multiband, T2*-weighted, gradient-echo, echo-planar imaging sequence was used to acquire 40 transverse slices at the sampling rate of 1 Hz and at the isotropic spatial resolution of 3 mm. Anatomical scans were also collected using a T1-weighted contrast sequence at the isotropic spatial resolution of 1 mm.

**Preprocessing** After correcting for the susceptibility artifacts using the reversed phase-encoding images with `topup` in FSL (v6.0.2; https://fsl.fmrib.ox.ac.uk/fsl/), SPM12 (v6225; https://www.fil.ion.ucl.ac.uk/spm/software/spm12/) was used for slice timing correction and realignment of the functional 4-D time series data. Advanced Normalization Tools (v2.3.5; https://github.com/ANTsX/ANTs) were used to perform symmetric diffeomorphic registration between individuals' anatomical 3-D images and the standard template 3-D image (`MNI152_T1_2mm_brain.nii.gz`) from FSL as well as the coregistration (rigid-body affine transform) between the anatomical 3-D image and the temporally averaged functional 3-D image within each subject. The rigid-body and diffeomorphic transformations were combined and applied to each volume of the realigned 4-D functional image, which was resampled only once at the isotropic resolution of 3 mm. Thereafter, the functional images were spatially smoothed with an isotropic Gaussian kernel (with a full width at half maximum of 6 mm). ICA-AROMA (v0.4.4.beta) was used to automatically reject 'noise' components to attenuate head motion artifacts and non-BOLD image intensity perturbations (https://github.com/maartenmennes/ICA-AROMA).

**Data dimensions** The analysed fMRI data consisted of 62,062 "brain" voxels (based on a brain mask created using `bet` in FSL), 178–525 time points (or seconds), 3 stimuli, and 39 subjects. Due to limited field of views in some subjects, only 61,572 voxels had valid values in the linearised encoding analysis results. The authors of the original study excluded 3 subjects from their analysis for either motion artifacts and emotional ratings but included in the shared data repository. In the current analysis, all 39 subjects were analysed for no apparent motion artifacts after ICA-AROMA.

#### S3.3 behavioural data

**Data source** As part of the fMRI study (Sachs et al., 2020), behavioural data of 39 healthy participants were downloaded from OpenNeuro (https://openneuro.org/datasets/ds003085/versions/1.0.0).

**Data acquisition** After the fMRI session, participants listened to the same stimuli while rating their evoked instantaneous emotions using a physical slider ("fader") in a silent room. Emotional scales of *Emotionality* (the intensity of the evoked feelings of sadness or happiness, depending on the intended emotion of each piece) and *Enjoyment* (the momentary feelings of enjoyment) were rated, one at a time. In total, a participant listened to an identical stimulus for three times including the fMRI session. The slider position values (an integer from 0 to 127) were sampled at about 30.3 Hz.

**Preprocessing** The imported time series were linearly detrended and downsampled at 5 Hz after an anti-aliasing low-pass filtering using the `resample` function in the MATLAB Signal Processing Toolbox (R2022b).

**Data dimensions** The analysed behavioural data comprised 2 emotional scales, 841–2,576 time points depending on the stimulus (178–525 seconds at 5 Hz), 3 stimuli, and 39 subjects.

#### S3.4 Linearised encoding analysis

**Features** True features $\mathbf{X}$ were the audio envelope extracted from the music samples for the well-established auditory response in various types of human brain data including fMRI (Giraud et al., 2000; Harms et al., 2005; Overath et al., 2012), EEG (Aiken & Picton, 2008), and intracranial electrocorticography (Kubanek et al., 2013). The envelope was created by summing the output of the "cochlear model" with a 128-channel filterbank, of which characteristic frequencies ranged loglinearly from 180 to 7,040 Hz, as in the NSL Auditory-cortical MATLAB Toolbox (v2001; http://nsl.isr.umd.edu/downloads.html), and then further downsampled to the sampling rate of the human data (EEG: 125 Hz; fMRI: 1 Hz, behavioural ratings: 5 Hz). Null features $\mathbf{U}$ were either (a) uniform noise, (b) normal noise, or (c) the phase-randomised envelope. Phase-randomisation preserves the amplitude spectrum of the envelope, thus preserves the temporal autocorrelation. This is necessary to correctly estimate the null correlation distribution of the autocorrelated noise. In previous studies (Kaneshiro et al., 2020; Leahy et al., 2021), phase-randomisation was used to generate an empirical null distribution for non-parametric statistical inference. The phase randomisation was done using fast Fourier transform (FFT), random rotation of phases while preserving complex conjugation for positive and negative frequencies, and followed by an inverse FFT. While this method ('unwindowed Fourier transform' in Theiler et al., 1992) may introduce spurious high frequencies for non-stationary signals, the envelope of stimuli that include zeros (i.e., silent periods) at the beginning and end of the musical stimuli can be assumed as stationary (i.e., the values of the last samples are similar to the first samples). To estimate the central tendency, 100 noise realisations were created for all cases. All features (either true or null) were standardised prior to fitting.

**FIR modelling** As in the simulations, an FIR model was regularised by ridge hyperparameters that are specific for response units (i.e., EEG channels, fMRI voxels, rating scales). Accounting for the inherent time scales of the measures, different delays of the features were used. To demonstrate the effect of the model's flexibility, 3 cases of delays (common choice, shorter, and longer) were used. For EEG, delays of [0 to 0.3 sec; 39 samples], [0 to 0.5 sec; 64 samples], and [0 to 1 sec; 126 samples] were used. For fMRI, [4 to 6 sec; 3 samples], and [3 to 9 sec; 7 samples], [0 to 12 sec; 13 samples]. For behaviours, [0 to 5 sec; 26 samples], [0 to 10 sec; 51 samples], and [0 to 15 sec; 76 samples]. To avoid transient onset/offset effects at the boundaries of the musical stimuli, the first and last 15 seconds were excluded from the analysis. Both stimuli and responses were standardised before the FIR modelling.

**Cross-validation** In all cases, the CV partition was 3-by-2 nested k-fold (i.e., 33%, 33%, 33% for training, validation, test sets) for the given structures of the real data (i.e., 3 stimuli per participant). To compare CV schemes, prediction accuracies were averaged across CV folds and models (either subject-specific or stimulus-specific).

**Statistical inference** The SDL effect is defined as the difference between null prediction accuracies: $\mathrm{SDL} = \bar{r}_{\mathrm{stim}}(\mathbf{U};\, \texttt{IsRep} = 1) - \bar{r}_{\mathrm{subj}}(\mathbf{U};\, \texttt{IsRep} = 0)$. The null hypothesis is that the expected

SDL effect is zero $\mathcal{H}_0 : \mathbb{E}(\mathrm{SDL}) = 0$ and the alterative hypothesis is that SDL effect is positive $\mathcal{H}_A : \mathbb{E}(\mathrm{SDL}) > 0$. Non-parametric $P$-values were computed by permutation test ($K =$ 10,000). That is, for 200 (100 randomisations $\times$ 2 CV-designs) vectors of prediction accuracies, the binary variable `IsRep` was randomly permuted ($\max = C(200, 100) > 10^{57}$), and the two-sample $t$-statistics between two CV-designs were calculated for 10,000 times to form a null distribution. Resulting one-sided $P$-values were further corrected for the multiple response units using false discovery rate (FDR) adjustment (Yekutieli & Benjamini, 1999) to control the family-wise error rate as $P_{\mathrm{FDR}} < 0.01$.

**Software implementation** All encoding analyses of the multimodal datasets (EEG, fMRI, behaviour) were done using an MATLAB package named Linearised Encoding Analysis (LEA; https://github.com/seunggookim/lea), developed by the author.

### S4 Supplementary Figures

#### S4.1 Real data results

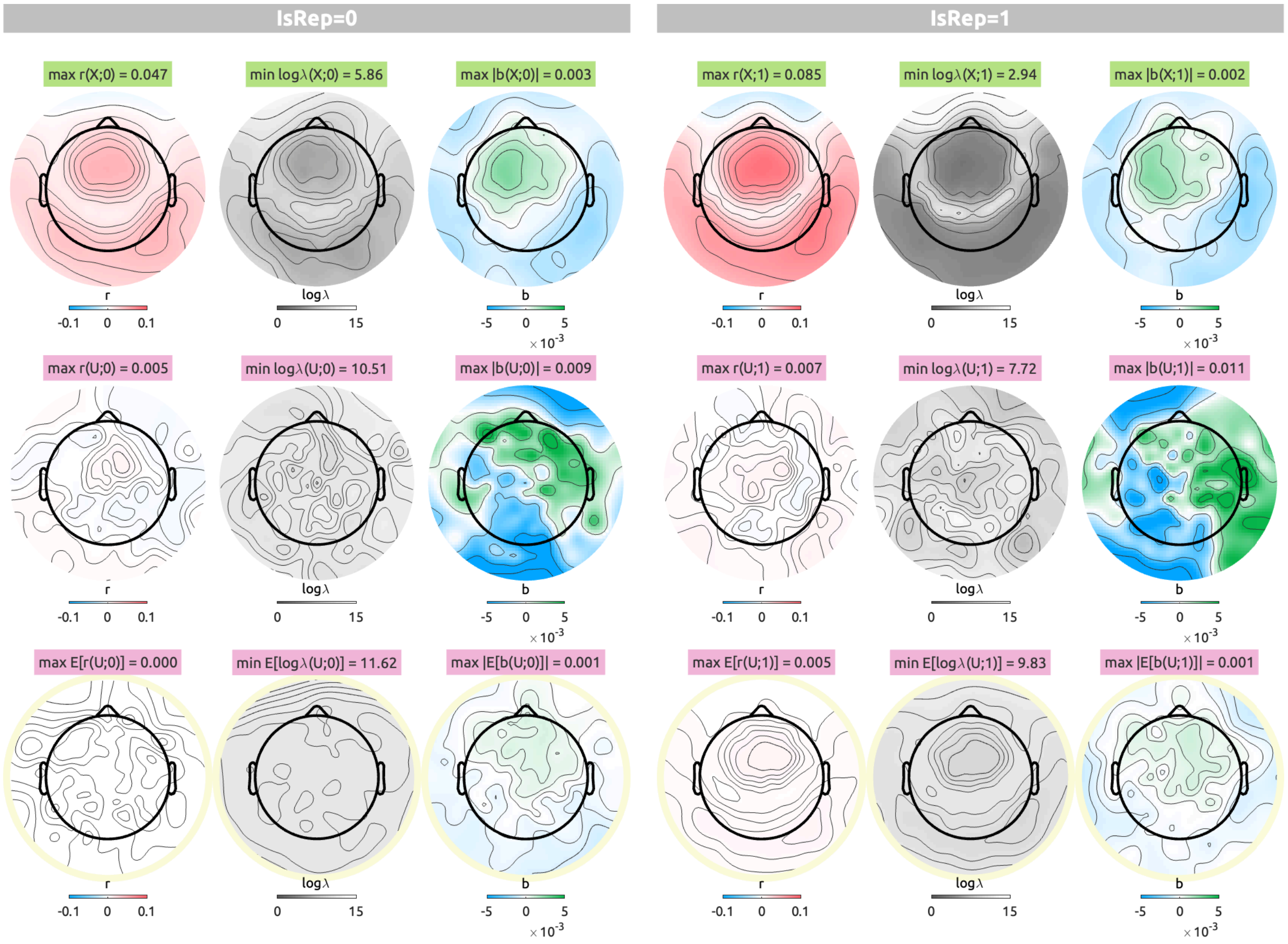


Figure S5: EEG linearised encoding analysis results with delays from 0 to 0.5 sec with an audio envelope (top row), a single case of the normal noise (middle row), and an average of 100 normal noises (bottom row; circled in pale yellow). For each CV scheme ($\texttt{IsRep} = 0$, left panels; $\texttt{IsRep} = 1$, right panels), prediction accuracy ($r$; blue to red), logarithmic ridge hyperparameter ($\log_{10} \lambda$; gray to white), summed weights ($b$; blue to green) are shown along the columns.

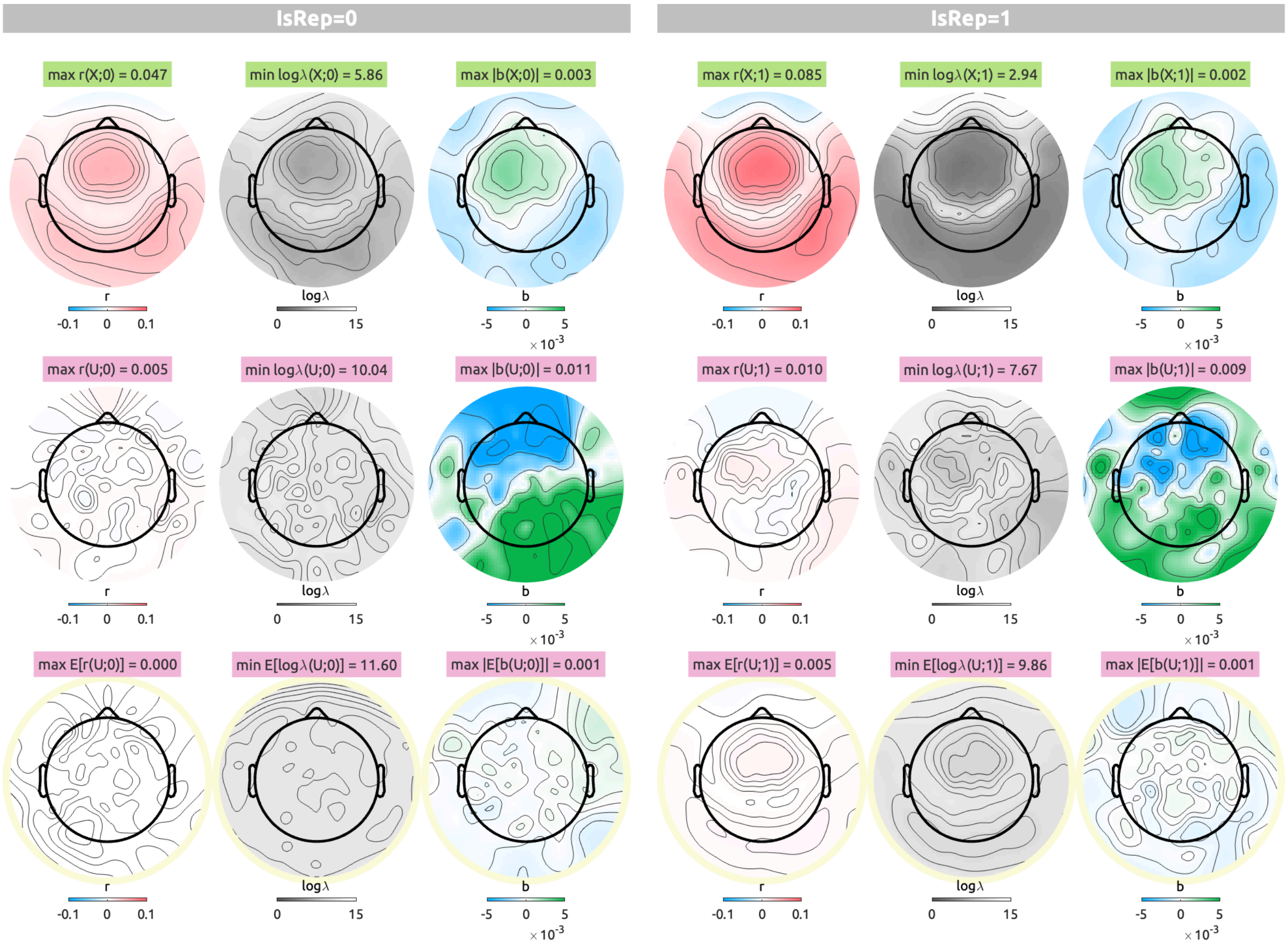


Figure S6: EEG topographies of linearised encoding analysis results with delays from 0 to 0.5 sec with the uniform noise as the null feature. The visualisation scheme is identical to Figure S5.

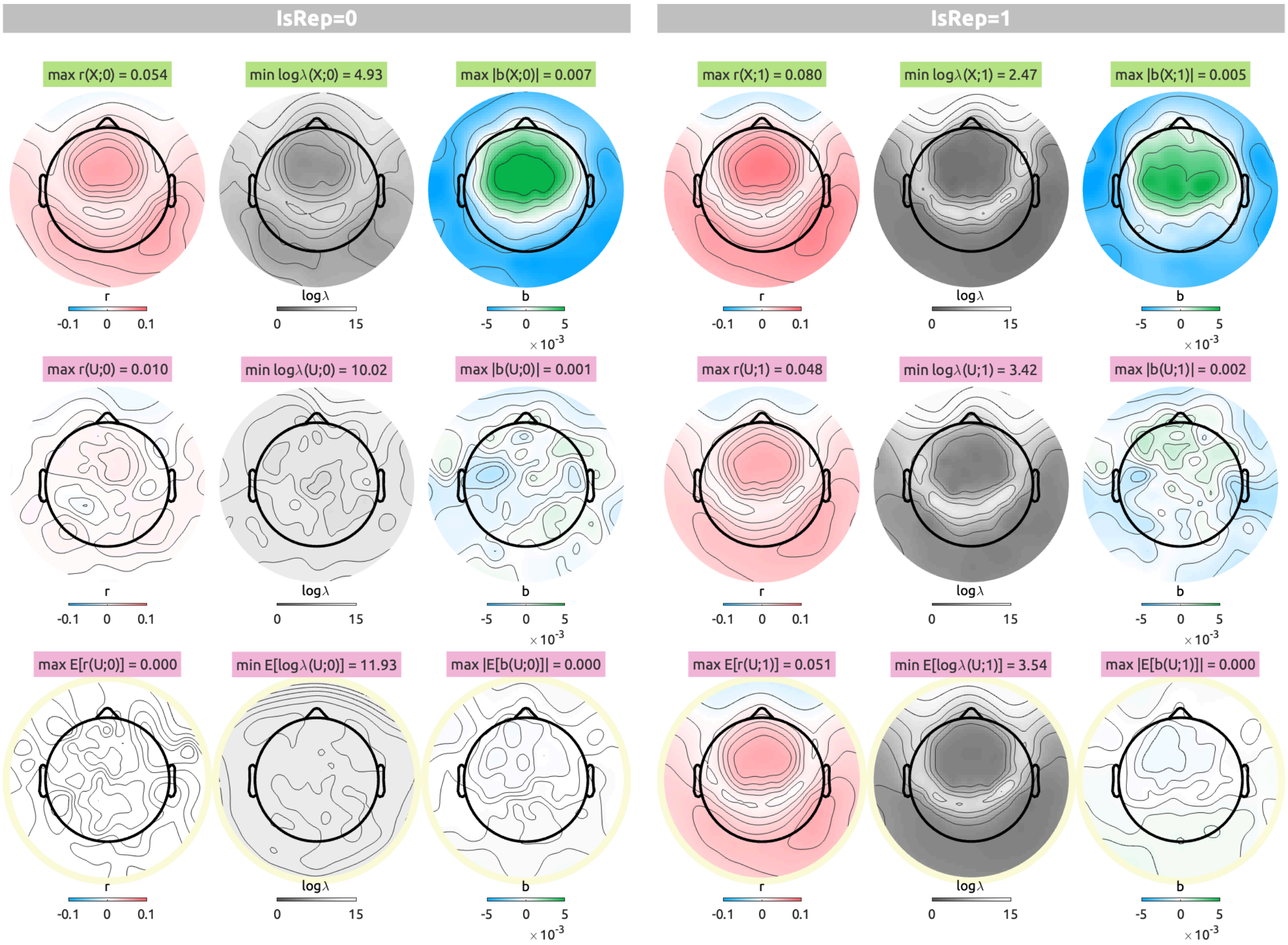


Figure S7: EEG topographies of linearised encoding analysis results with a delay from 0 to 0.3 sec with the phase-randomised envelope as the null feature. The visualisation scheme is identical to Figure S5.

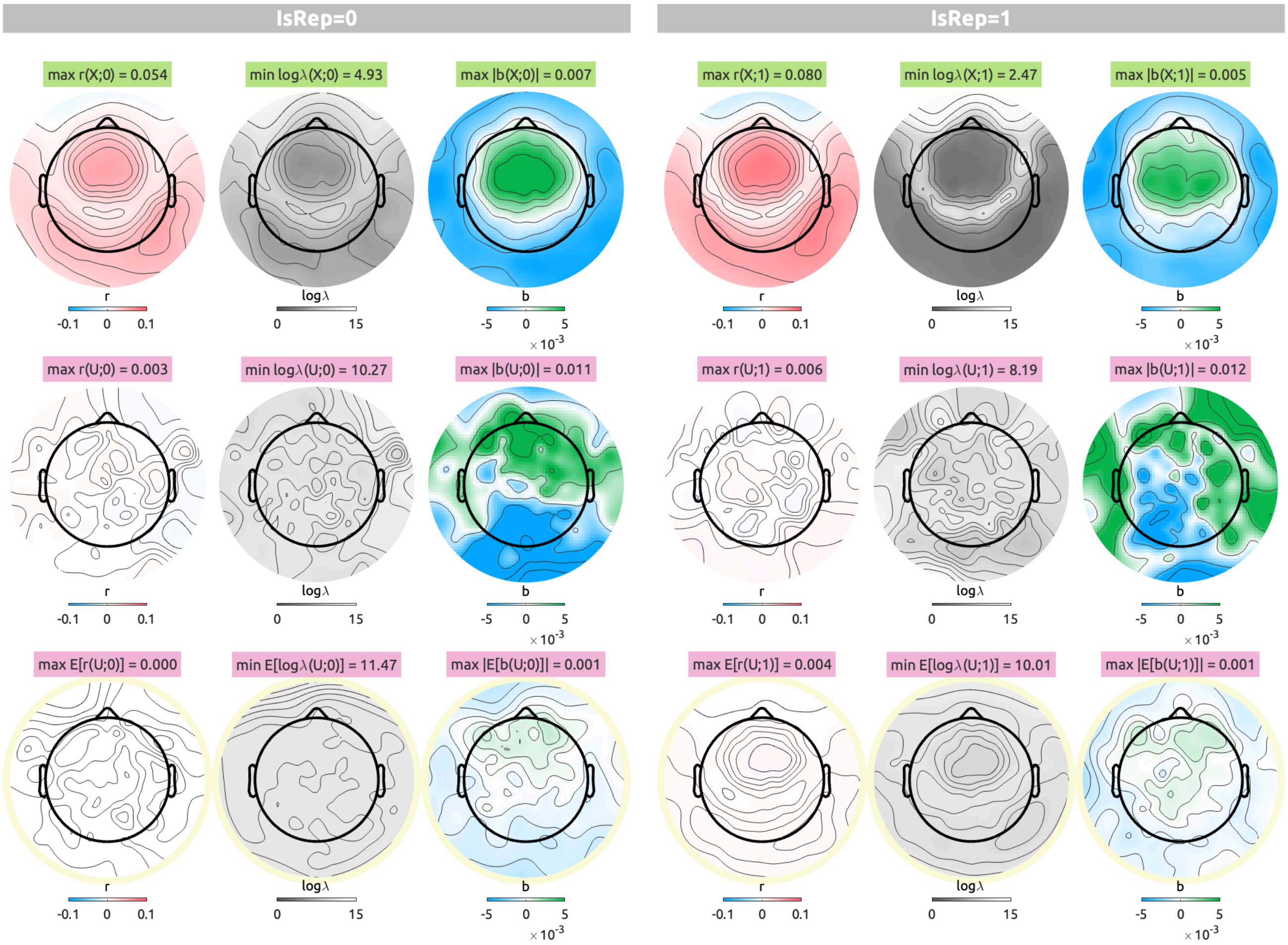


Figure S8: EEG topographies of linearised encoding analysis results with a delay from 0 sec to 0.3 sec with the normal noise as the null feature. The visualisation scheme is identical to Figure S5.

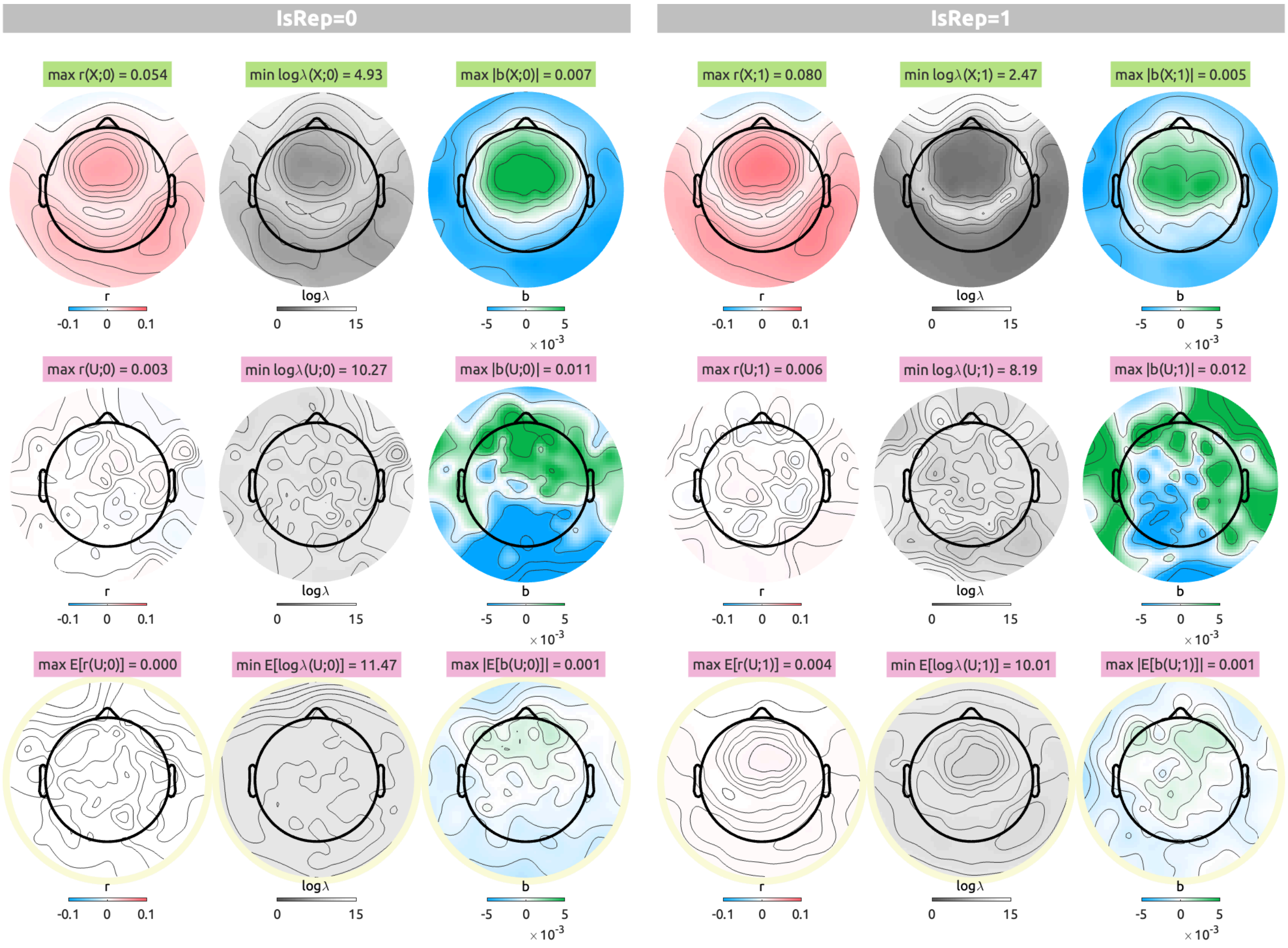


Figure S9: EEG topographies of linearised encoding analysis results with a delay from 0 sec to 0.3 sec with the normal noise as the null feature. The visualisation scheme is identical to Figure S5.

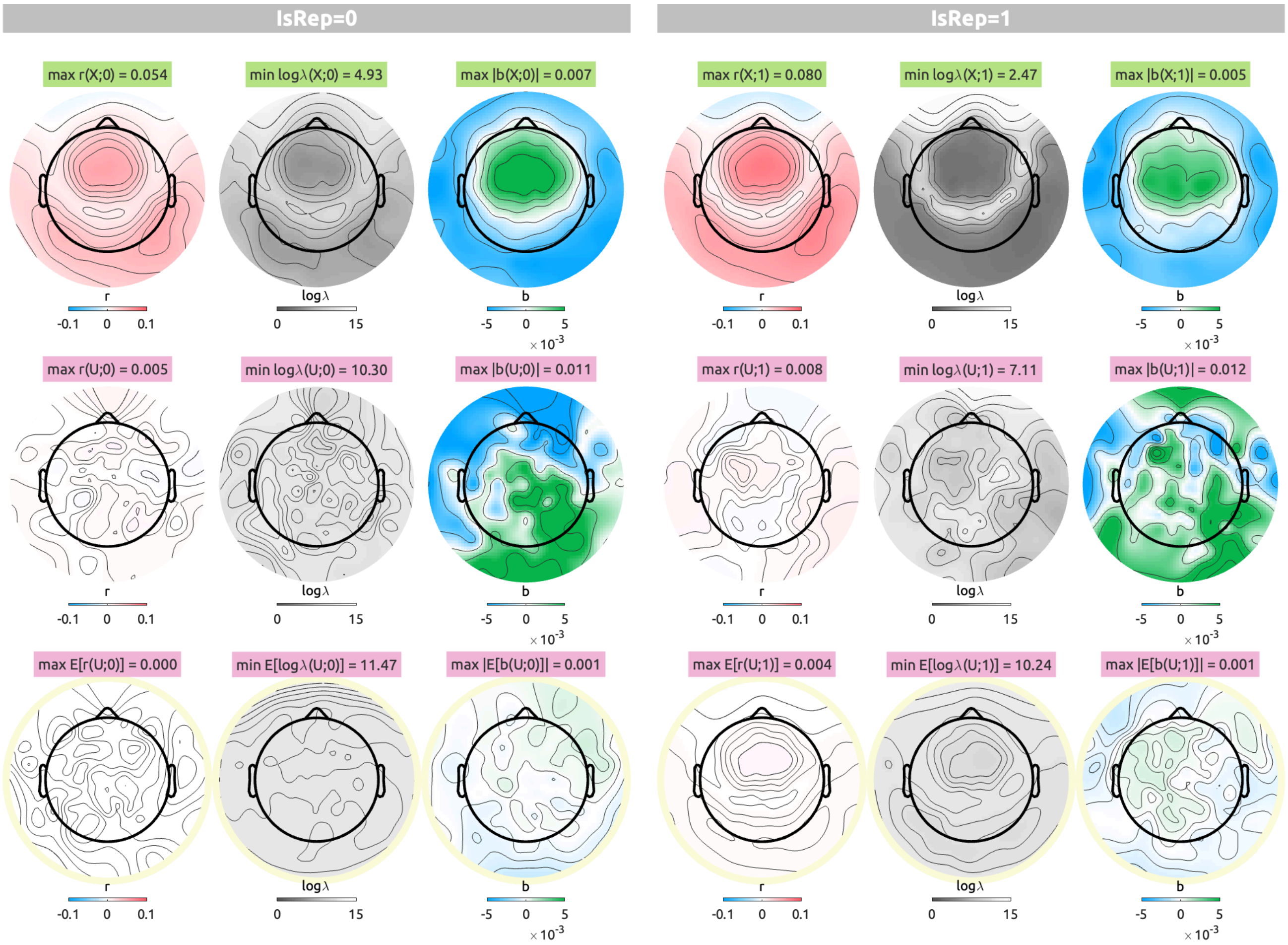


Figure S10: EEG topographies of linearised encoding analysis results with a delay from 0 to 0.3 sec with the uniform envelope as the null feature. The visualisation scheme is identical to Figure S5.

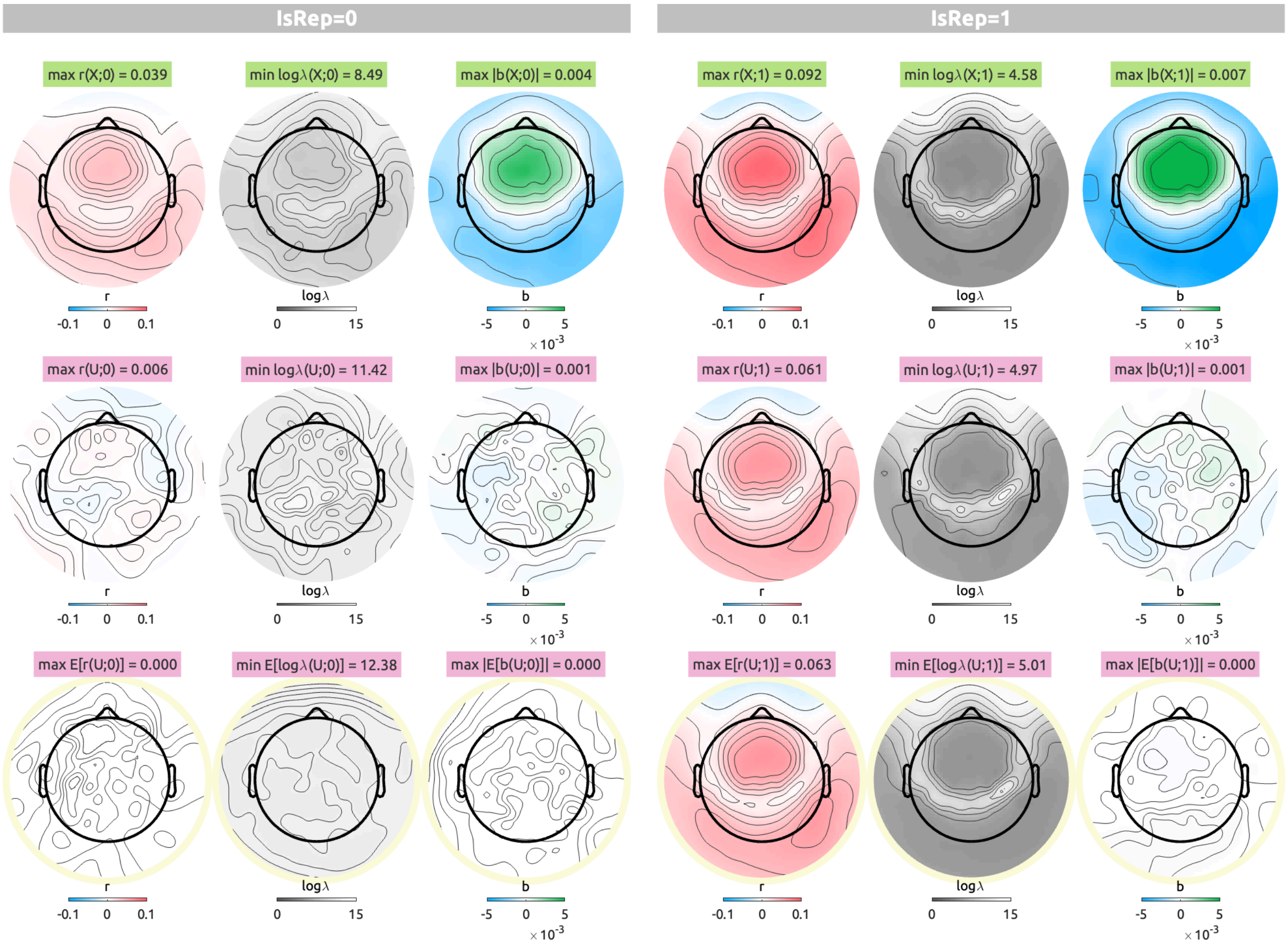


Figure S11: EEG topographies of linearised encoding analysis results with delays from 0 to 1 sec with the phase-randomised envelope as the null feature. The visualisation scheme is identical to Figure S5.

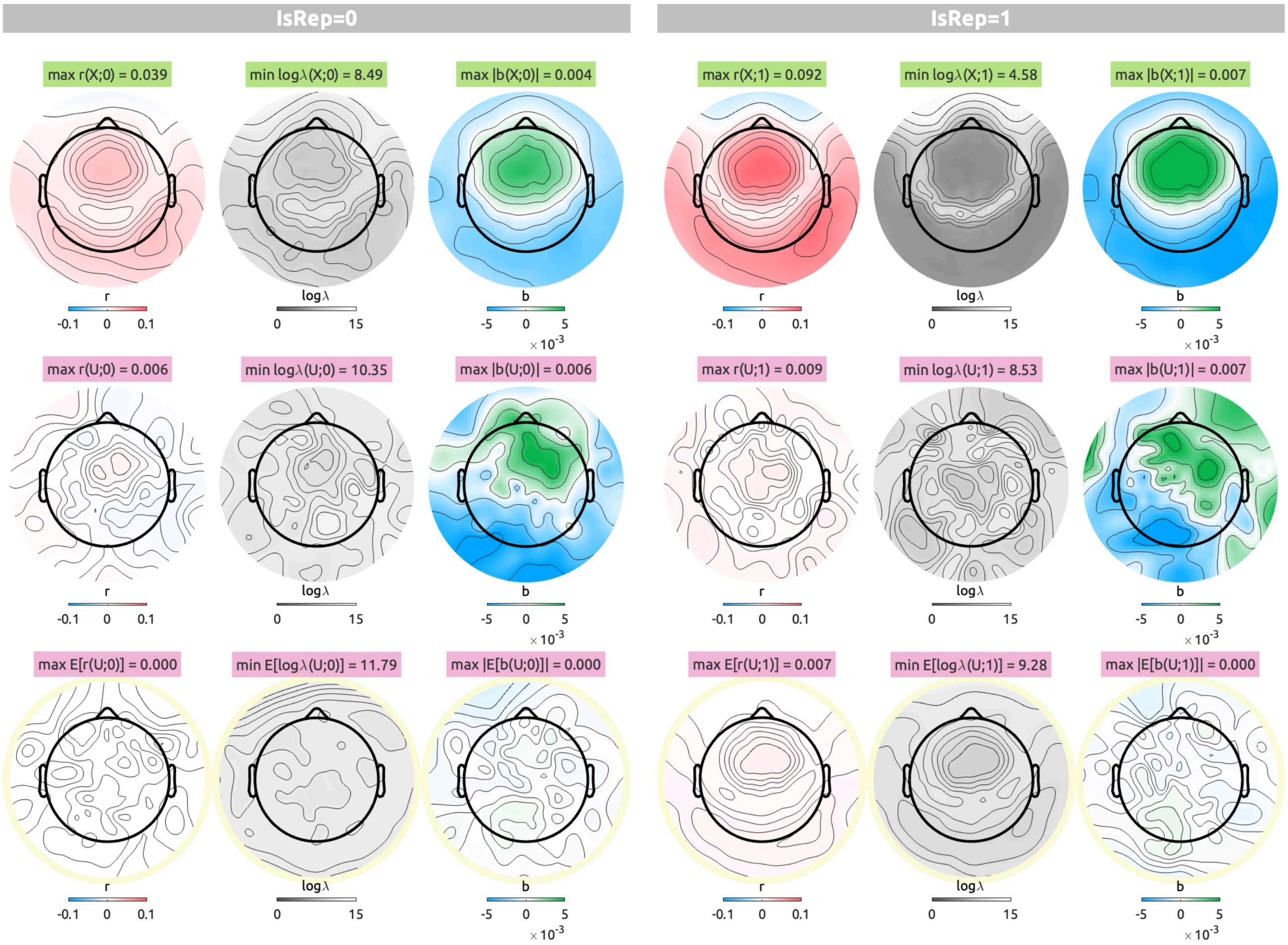


Figure S12: EEG topographies of linearised encoding analysis results with delays from 0 to 1 sec with the normal noise as the null feature. The visualisation scheme is identical to Figure S5.

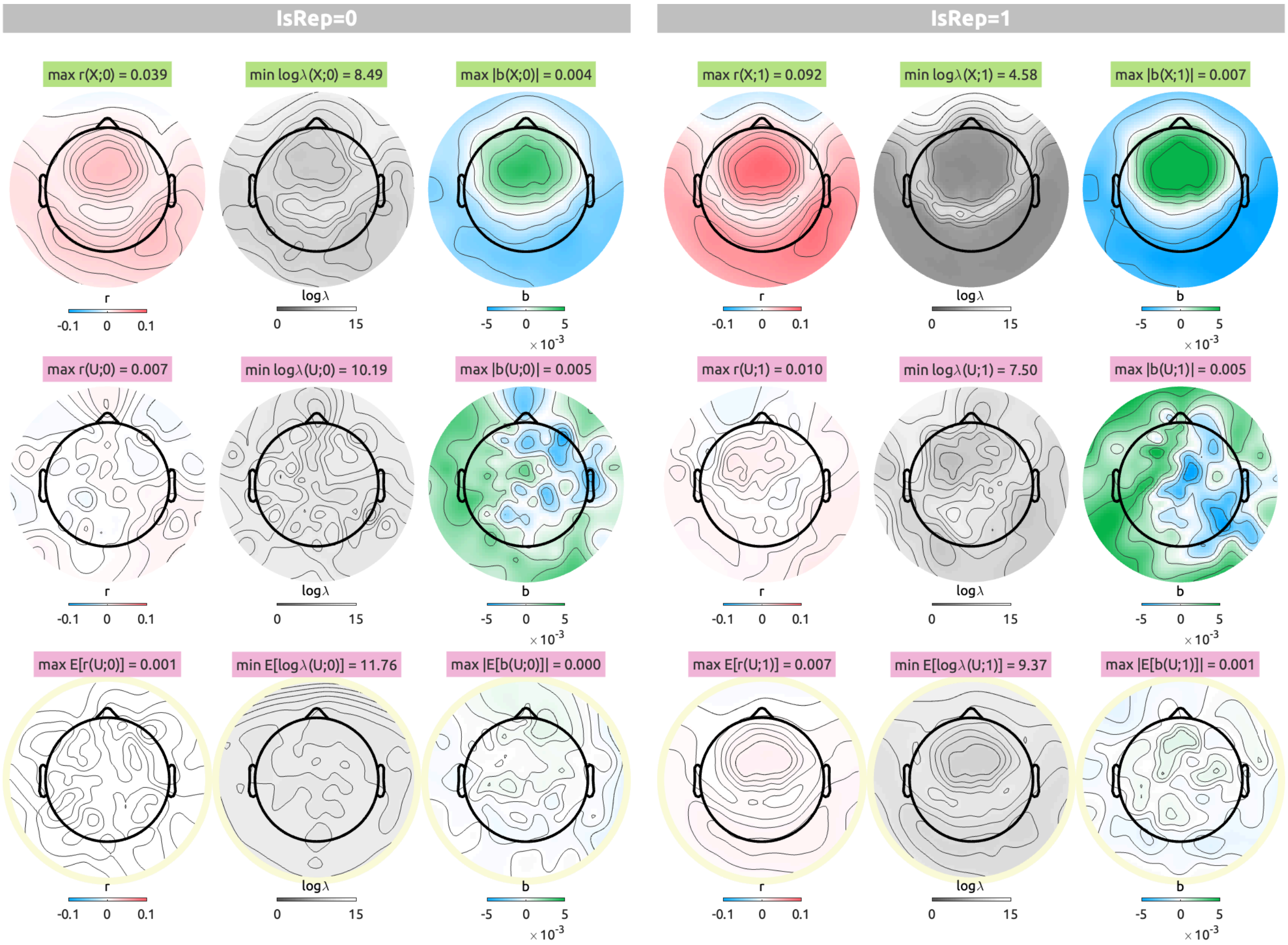


Figure S13: EEG topographies of linearised encoding analysis results with delays from 0 to 1 sec with the uniform noise as the null feature. The visualisation scheme is identical to Figure S5.

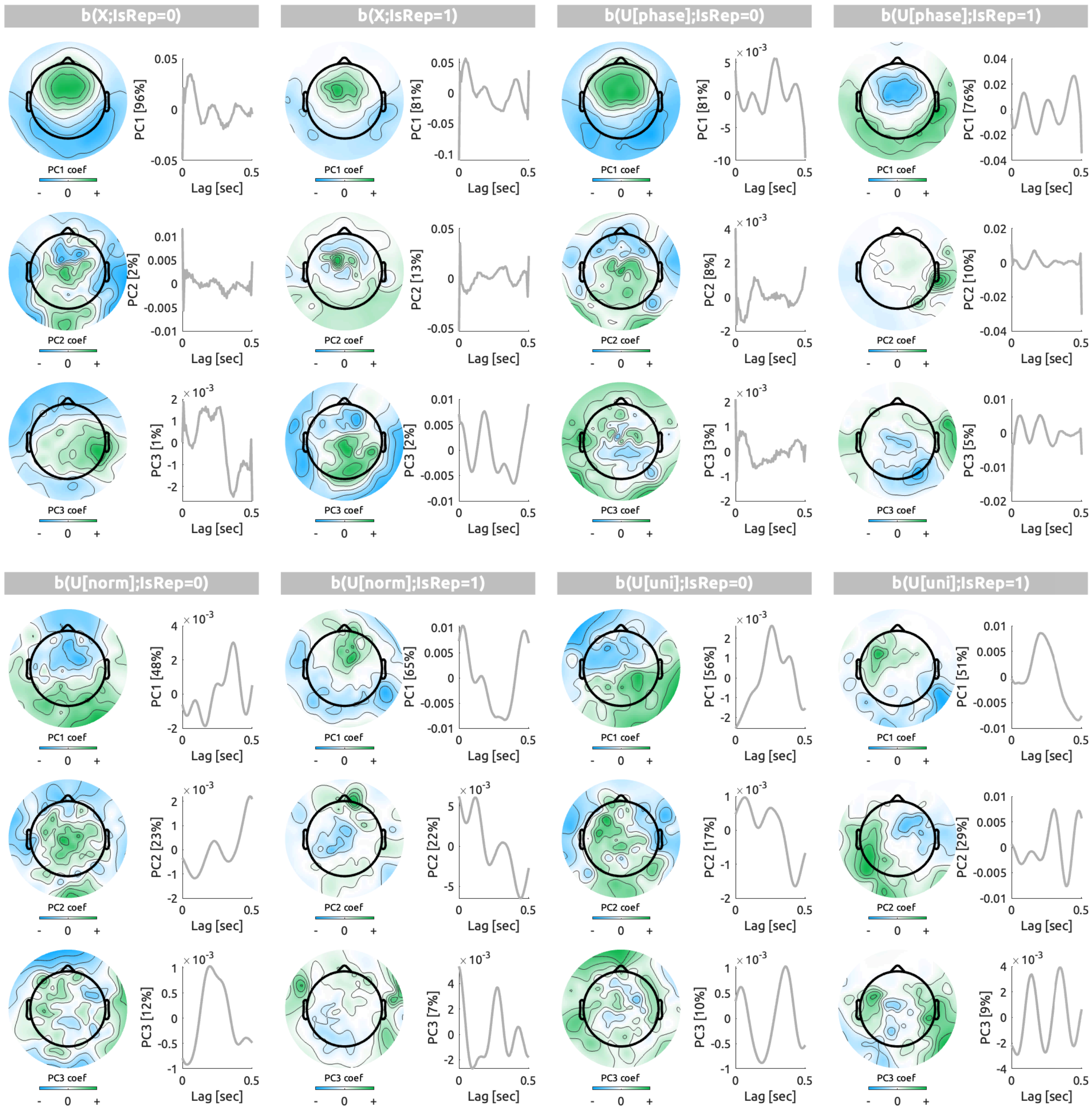


Figure S14: First three principal components (PCs) of the transfer function weights ($b$) of the models with delays from 0 to 0.5 sec are displayed by the topography of eigenvectors and the time series of eigenvariates with the explained variance noted. The weights are grouped by features ($X$, true audio envelope; $U[phase]$, phase-randomised envelope; $U[norm]$, normal noise; $U[uni]$, uniform noise) and the CV schemes ($\texttt{IsRep} = 0$, $\texttt{IsRep} = 1$).

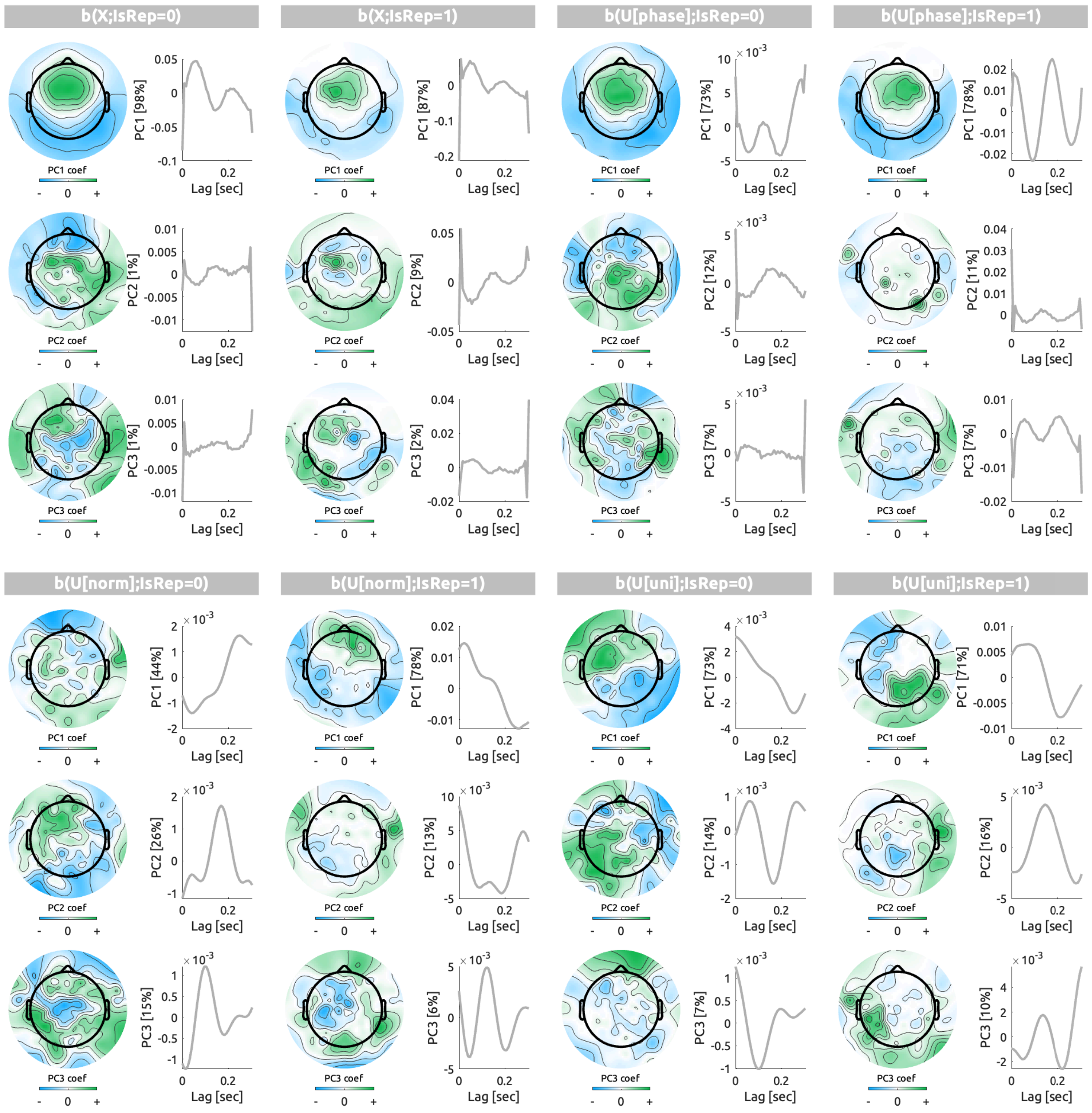


Figure S15: First three principal components (PCs) of the transfer function weights ($b$) of the models with delays from 0 to 0.3 sec are displayed. The visualisation scheme is identical to Figure S14

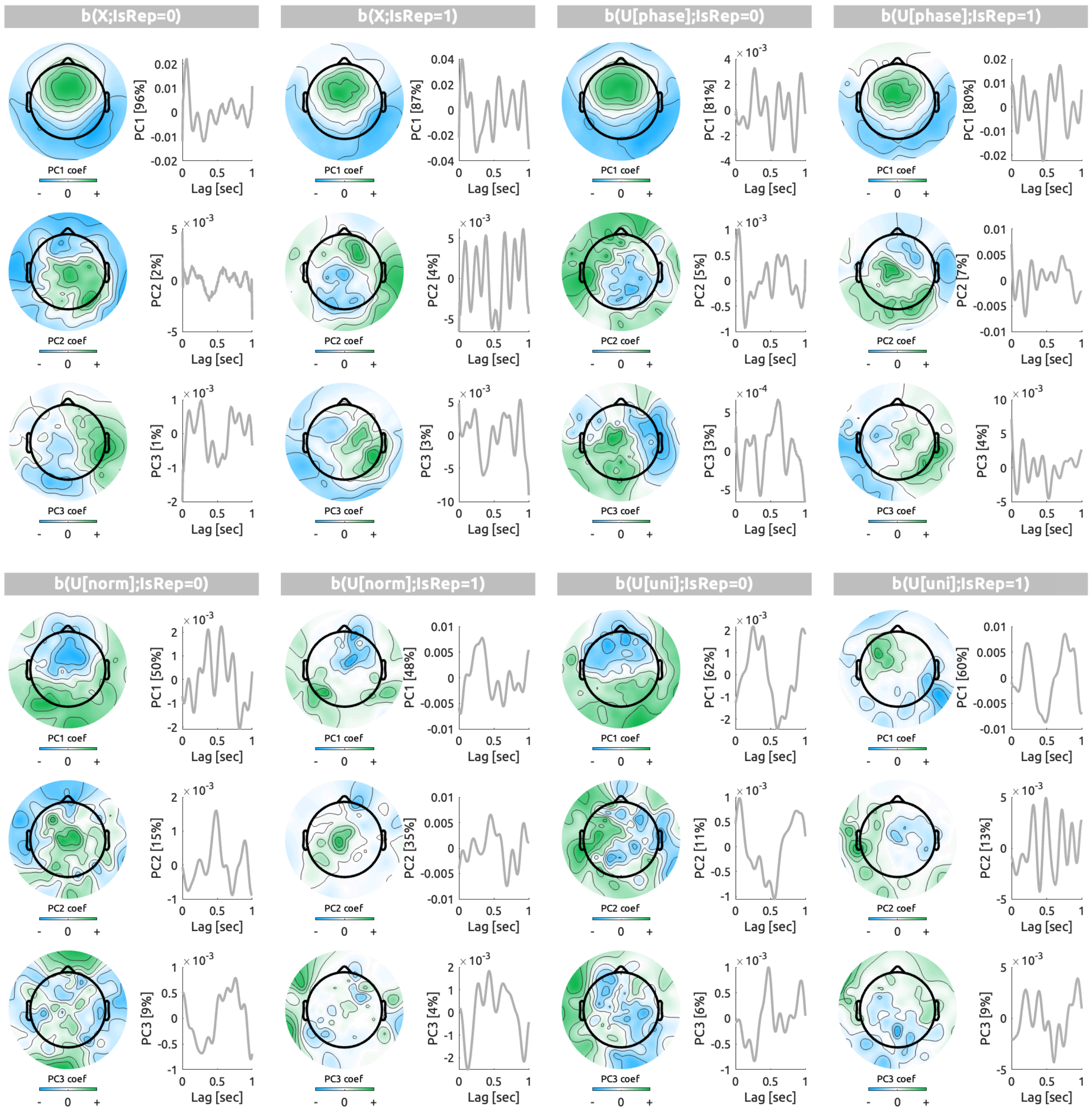


Figure S16: First three principal components (PCs) of the transfer function weights ($b$) of the models with delays from 0 to 1 sec are displayed. The visualisation scheme is identical to Figure S14

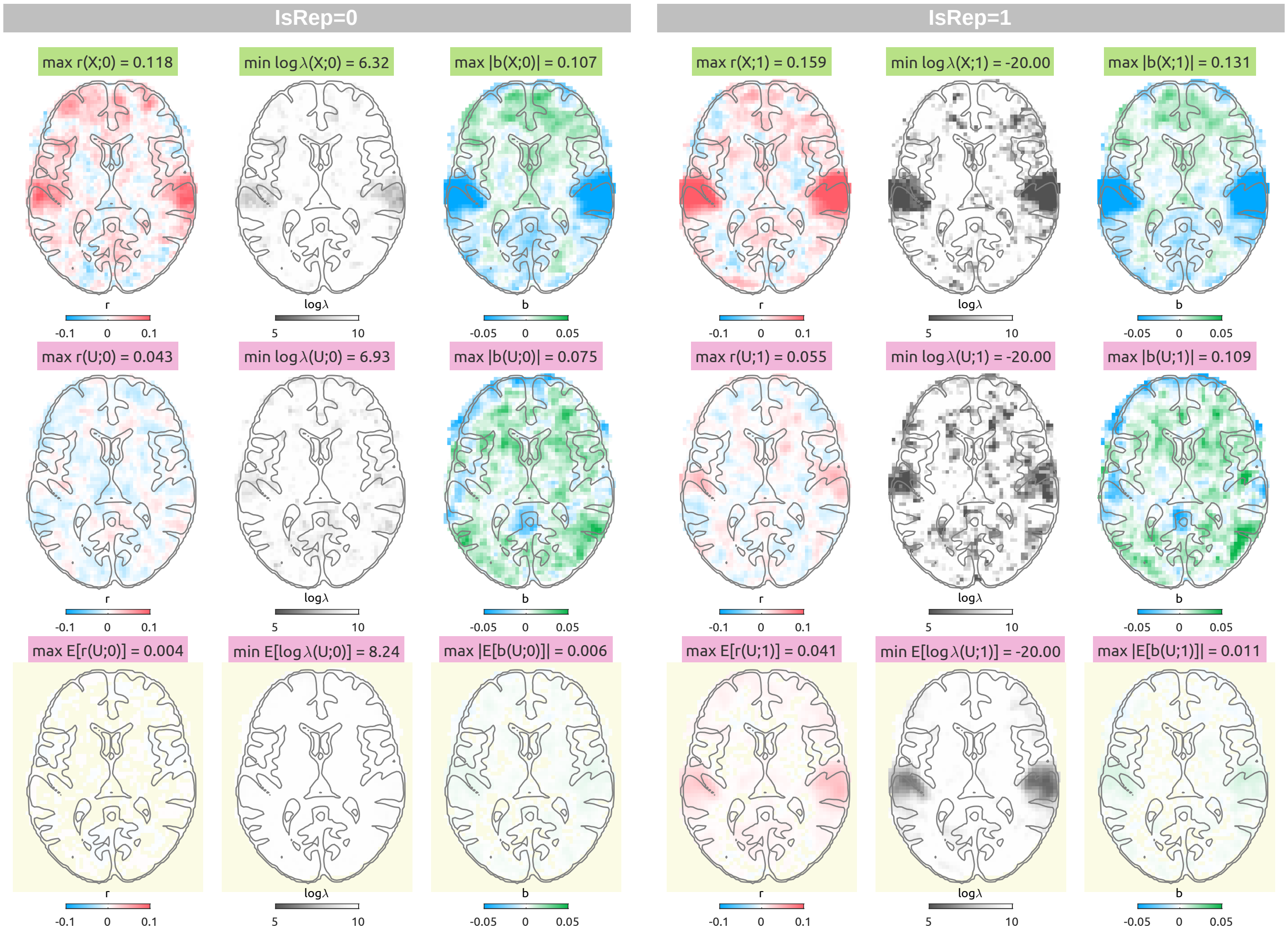


Figure S17: fMRI linearised encoding analysis results with delays from 3 to 9 sec with an audio envelope (top row), a single case of the normal noise (middle row), and an average of 100 phase-randomised envelopes (bottom row, pale yellow background). For each CV scheme ($\texttt{IsRep} = 0$, left panels; $\texttt{IsRep} = 1$, right panels), prediction accuracy ($r$, blue to red), logarithmic ridge hyperparameter ($\log_{10} \lambda$, gray to white), transfer function weights that are summed over delays ($b$, blue to green) are shown along the columns. The 3-D volumes can be viewed with the NeuroVault web viewer (https://identifiers.org/neurovault.collection:19626).

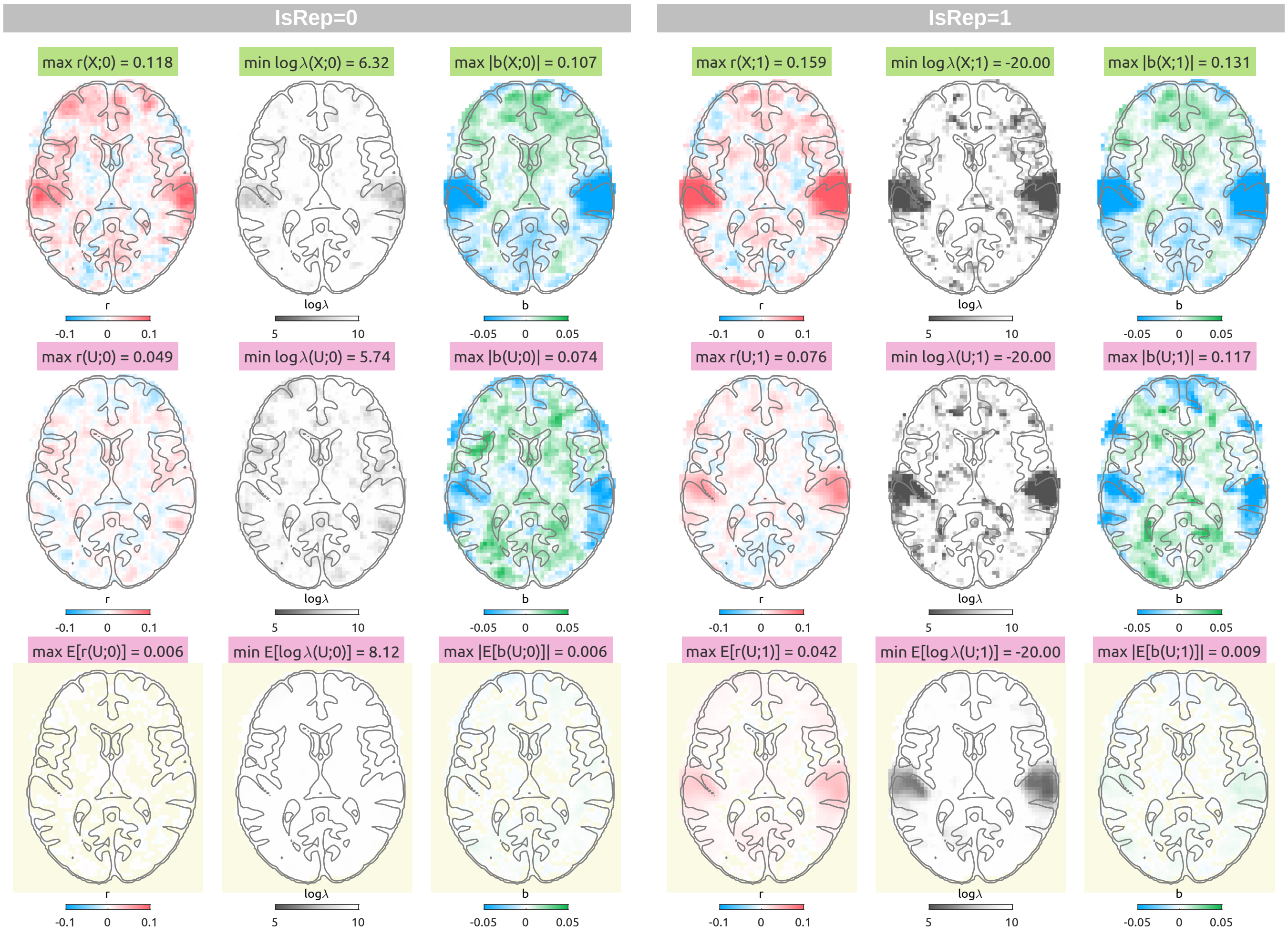


Figure S18: fMRI linearised encoding analysis results with delays from 3 to 9 sec with the uniform noise as the null feature. The visualisation scheme is identical to Figure S17.

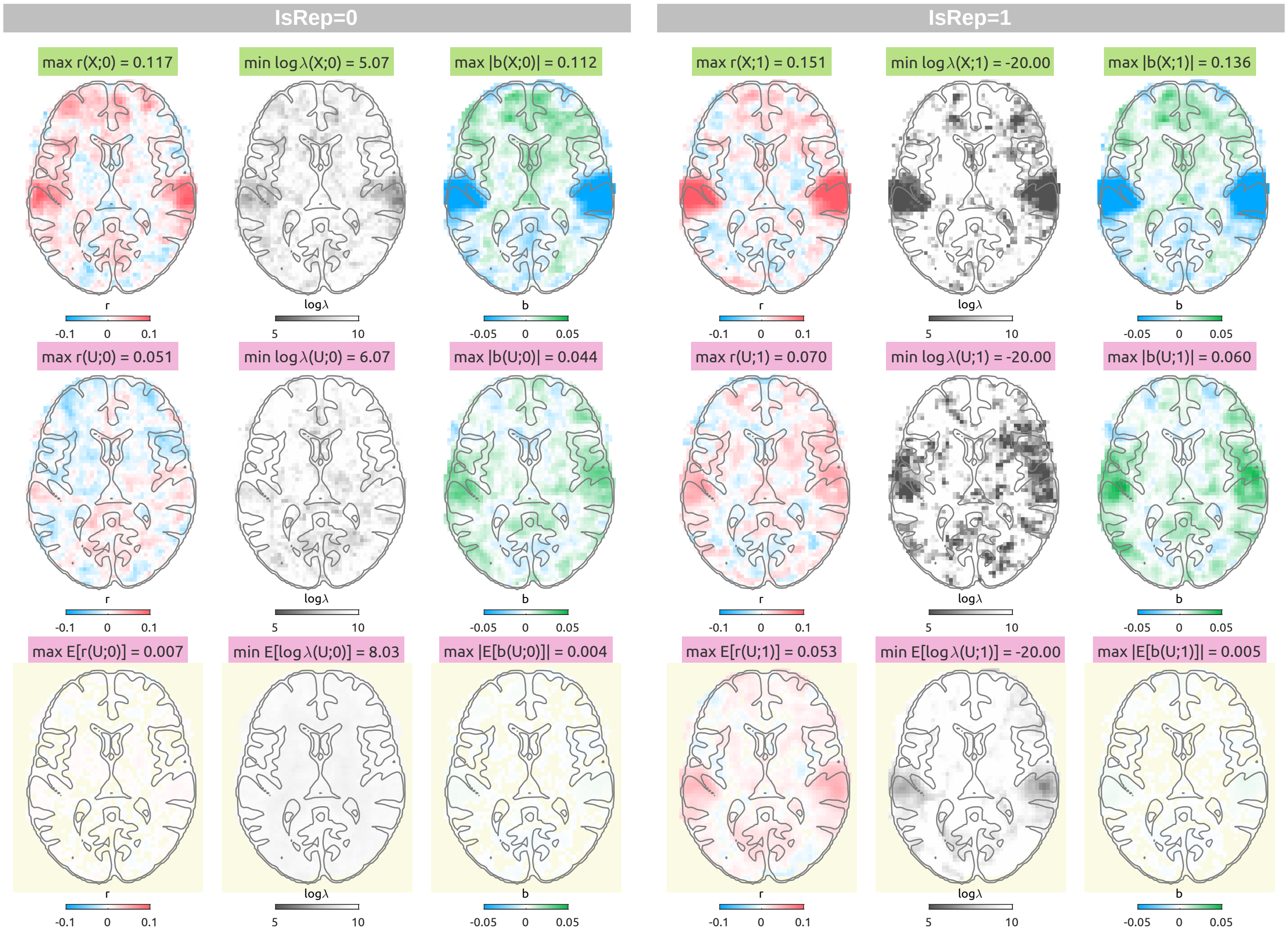


Figure S19: fMRI linearised encoding analysis results with delays from 4 to 6 sec with the phase-randomised envelope as the null feature. The visualisation scheme is identical to Figure S17.

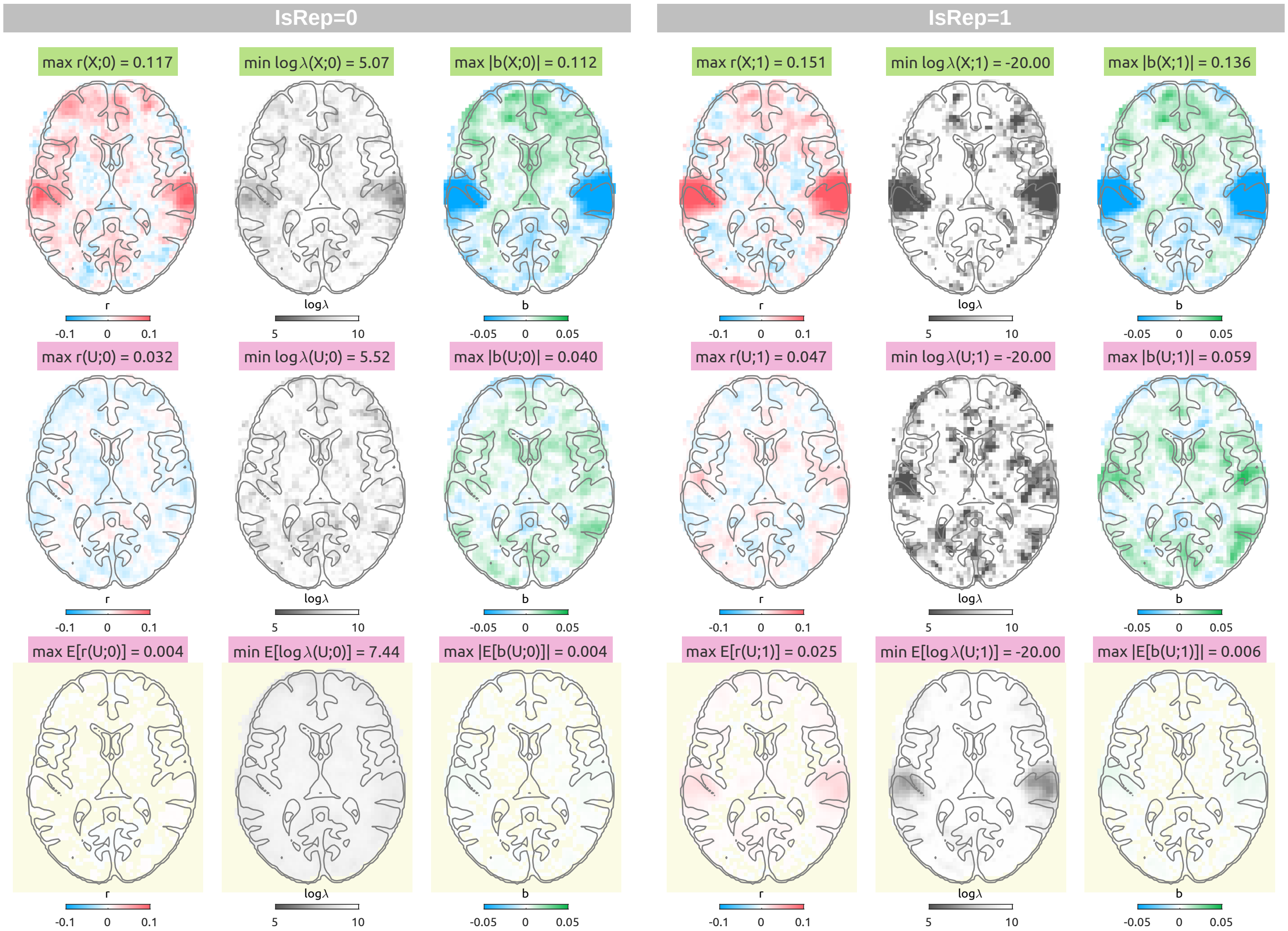


Figure S20: MRI linearised encoding analysis results with delays from 4 to 6 sec with the normal noise as the null feature. The visualisation scheme is identical to Figure S17.

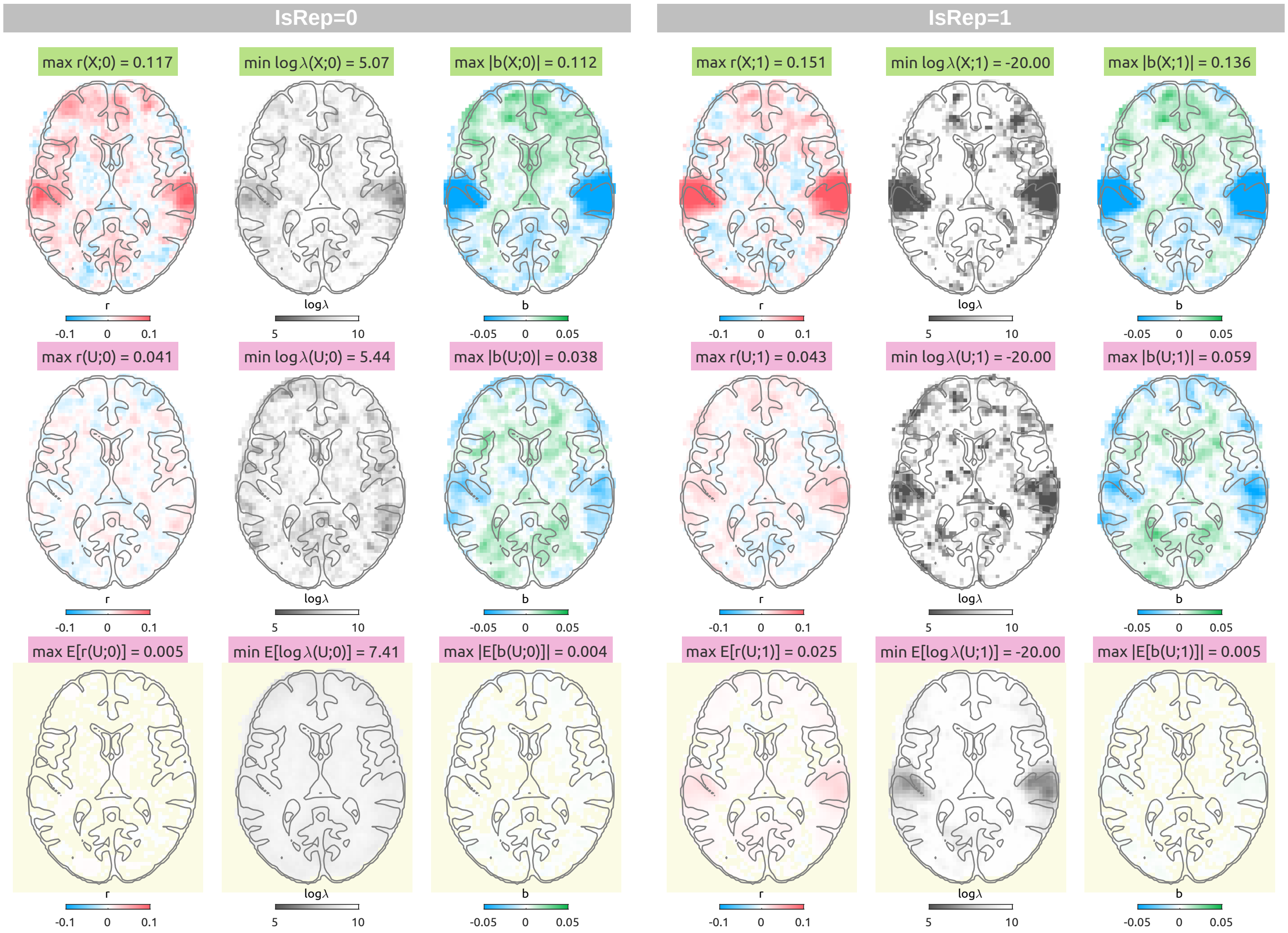


Figure S21: fMRI linearised encoding analysis results with delays from 4 to 6 sec with the uniform noise as the null feature. The visualisation scheme is identical to Figure S17.

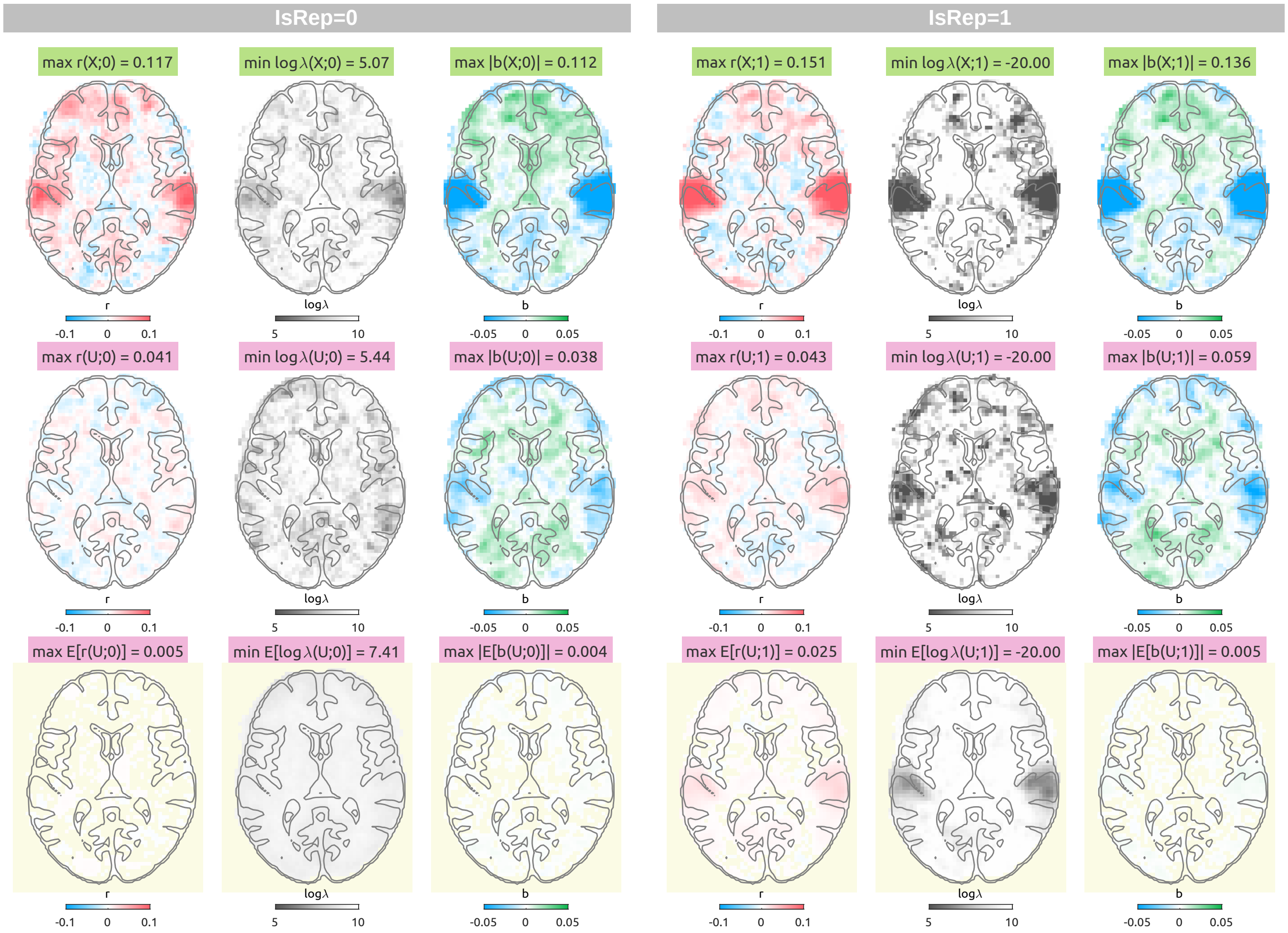


Figure S22: fMRI linearised encoding analysis results with delays from 0 to 12 sec with the phase-randomised envelope as the null feature. The visualisation scheme is identical to Figure S17.

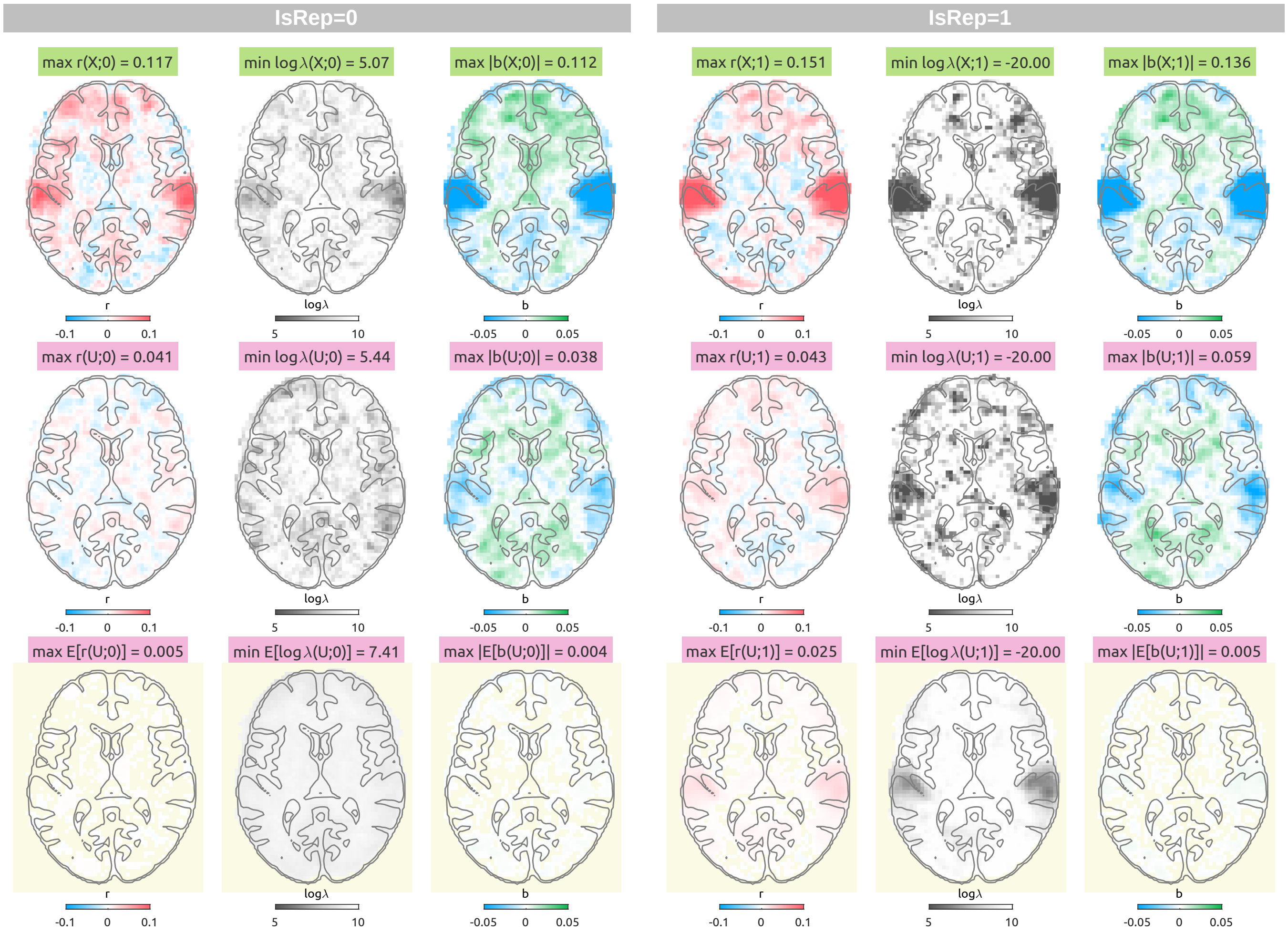


Figure S23: fMRI linearised encoding analysis results with delays from 0 to 12 sec with the phase-randomised envelope as the null feature. The visualisation scheme is identical to Figure S17.

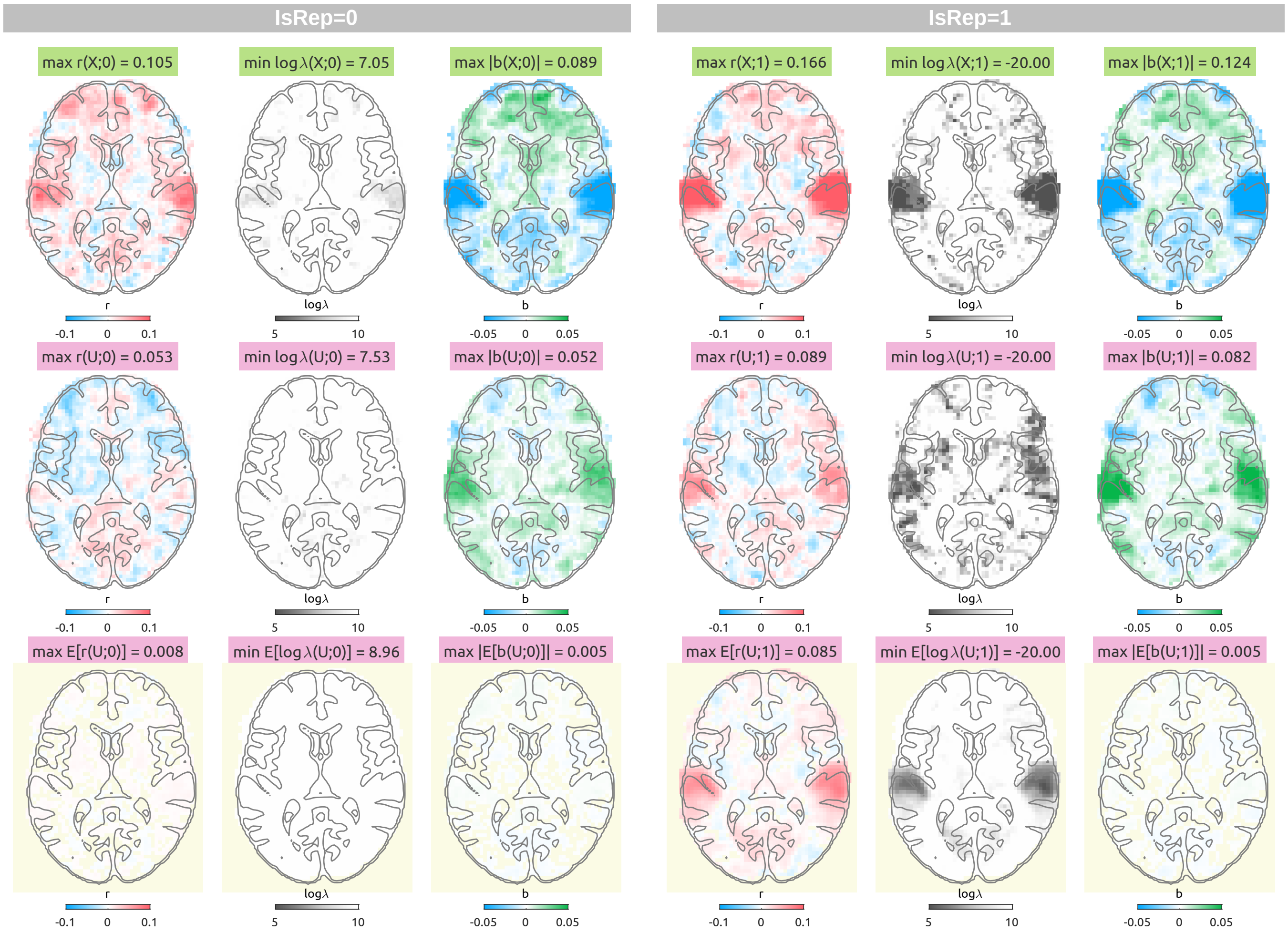


Figure S24: fMRI linearised encoding analysis results with delays from 0 to 12 sec with the phase-randomised envelope as the null feature. The visualisation scheme is identical to Figure S17.

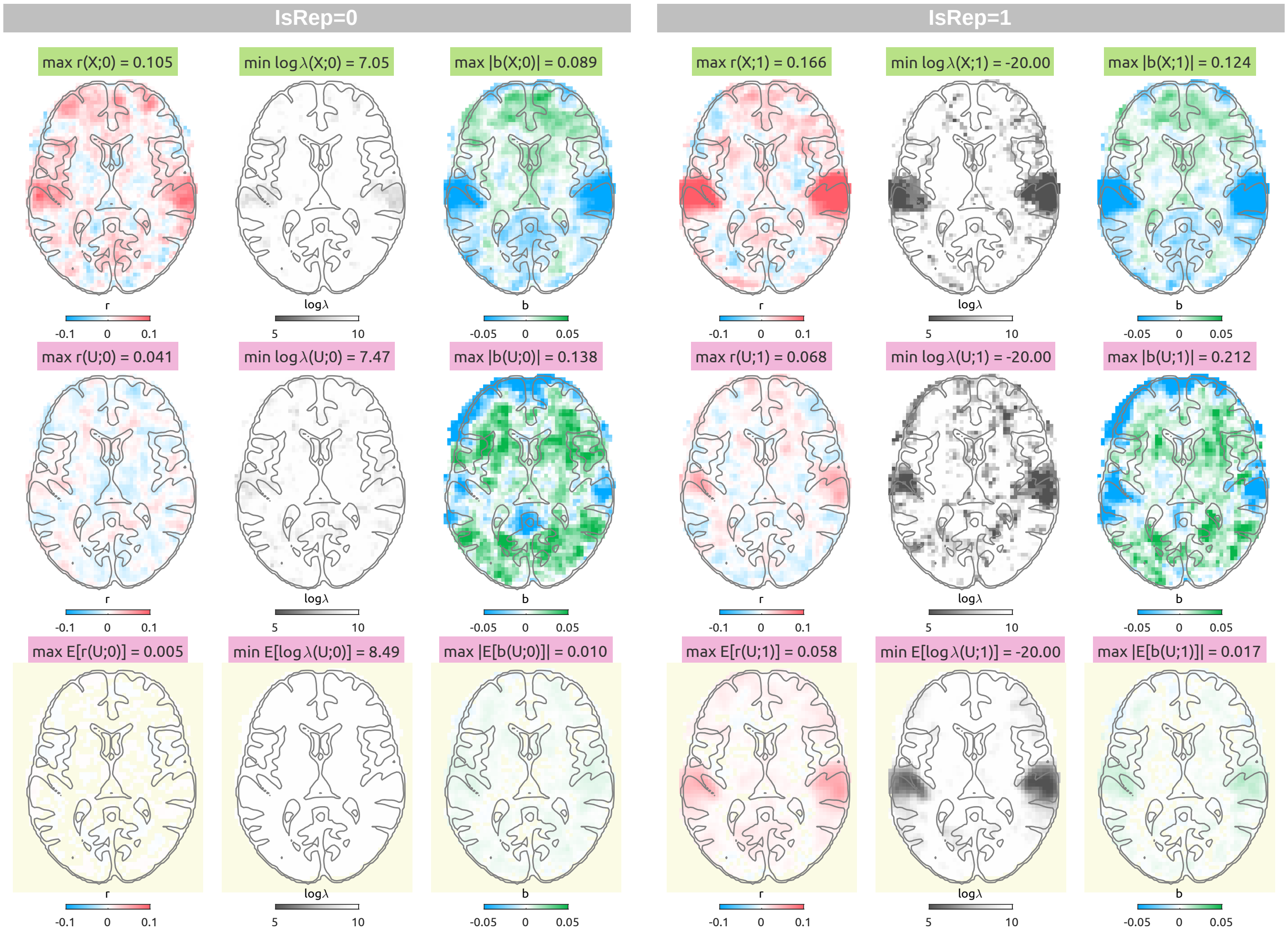


Figure S25: fMRI linearised encoding analysis results with delays from 4 to 6 sec with the normal noise as the null feature. The visualisation scheme is identical to Figure S17.

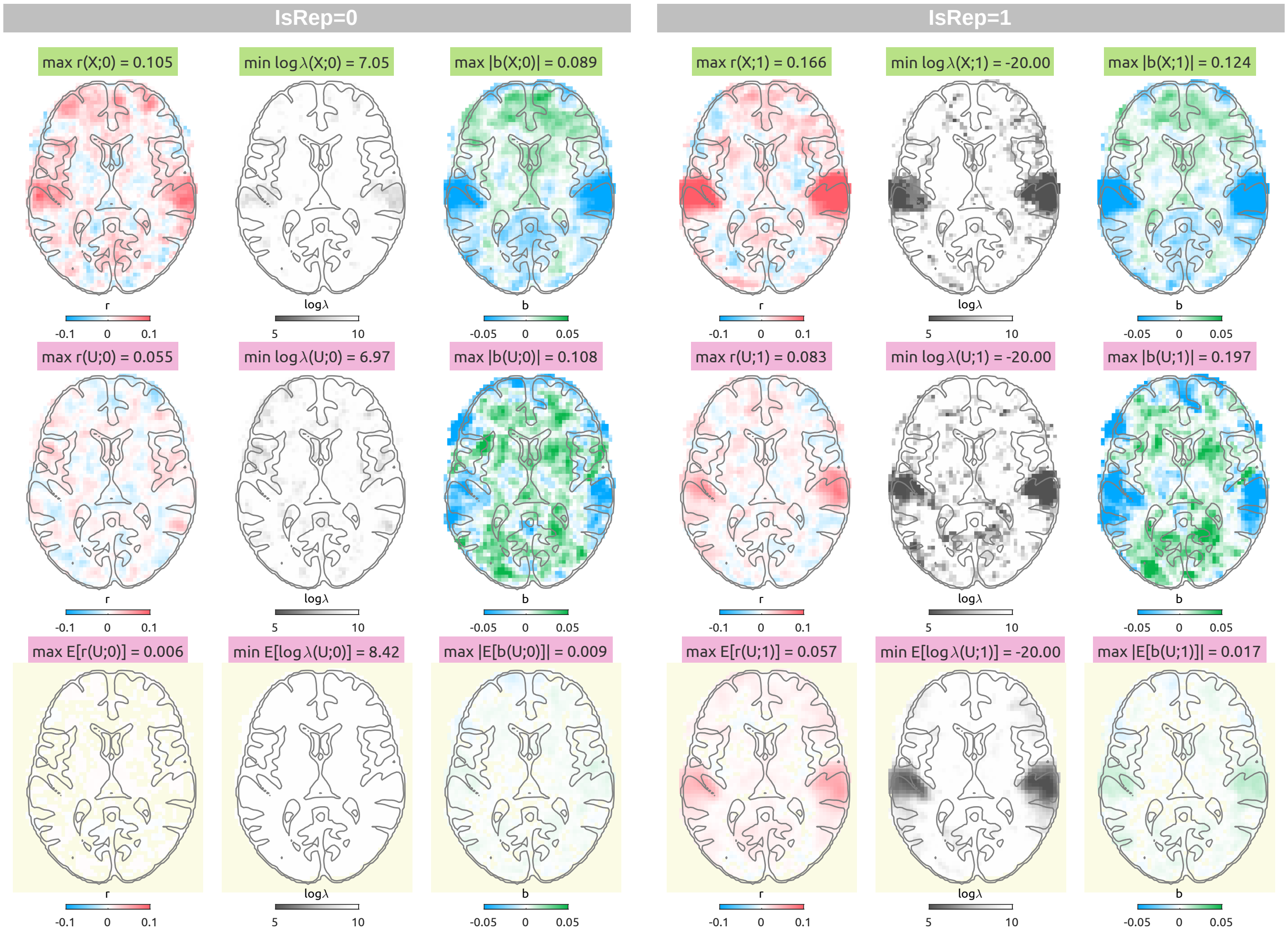


Figure S26: fMRI linearised encoding analysis results with delays from 0 to 12 sec with the uniform noise as the null feature. The visualisation scheme is identical to Figure S17.

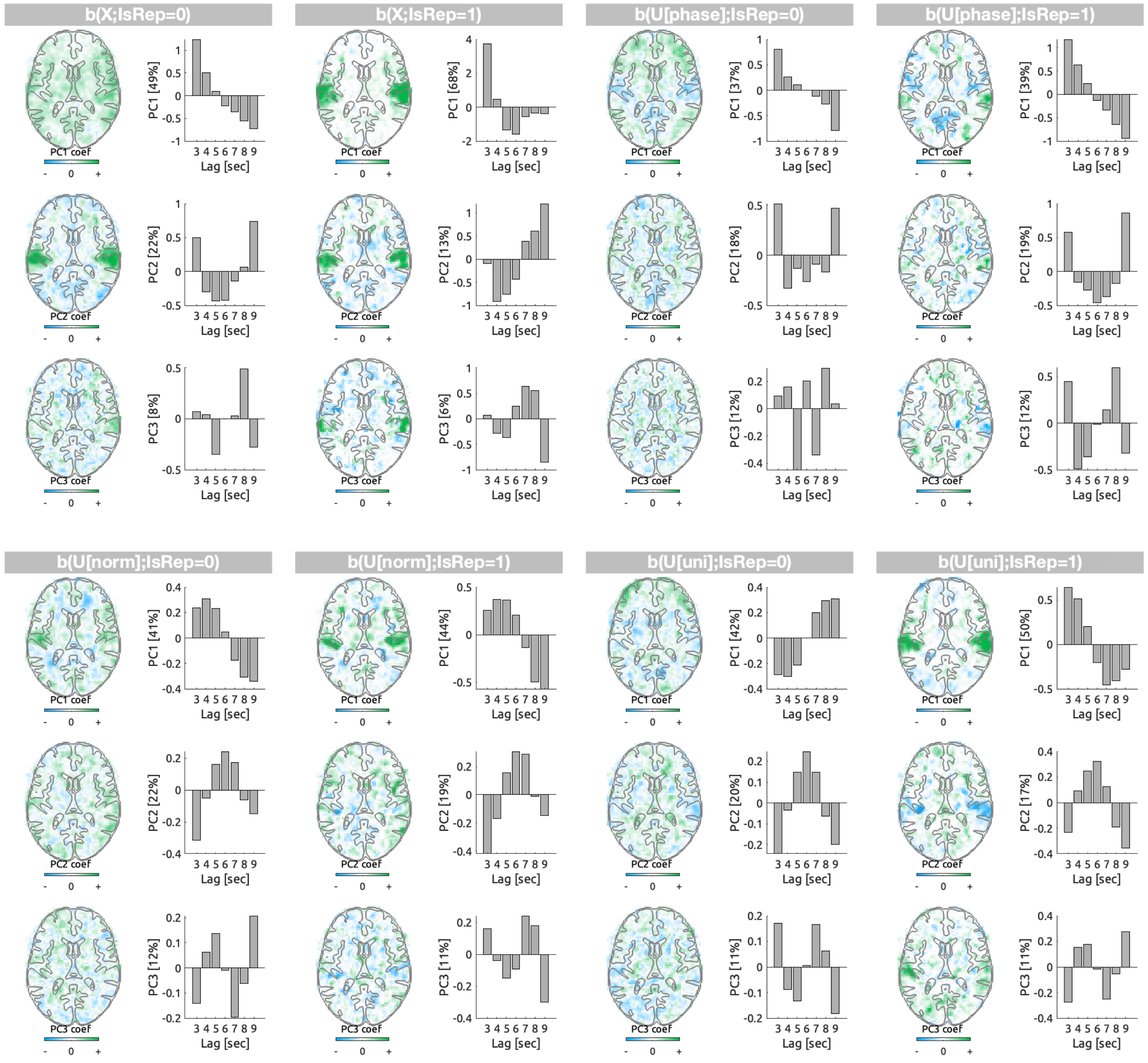


Figure S27: First three principal components (PCs) of the transfer function weights ($b$) of the models with delays from 3 to 9 sec are displayed in the transverse slice of eigenvectors and the time series of eigenvariates with the explained variance noted. The weights are grouped by features ($X$, true audio envelope; $U[phase]$, phase-randomised envelope; $U[norm]$, normal noise; $U[uni]$, uniform noise) and the CV schemes ($\texttt{IsRep} = 0$, $\texttt{IsRep} = 1$).

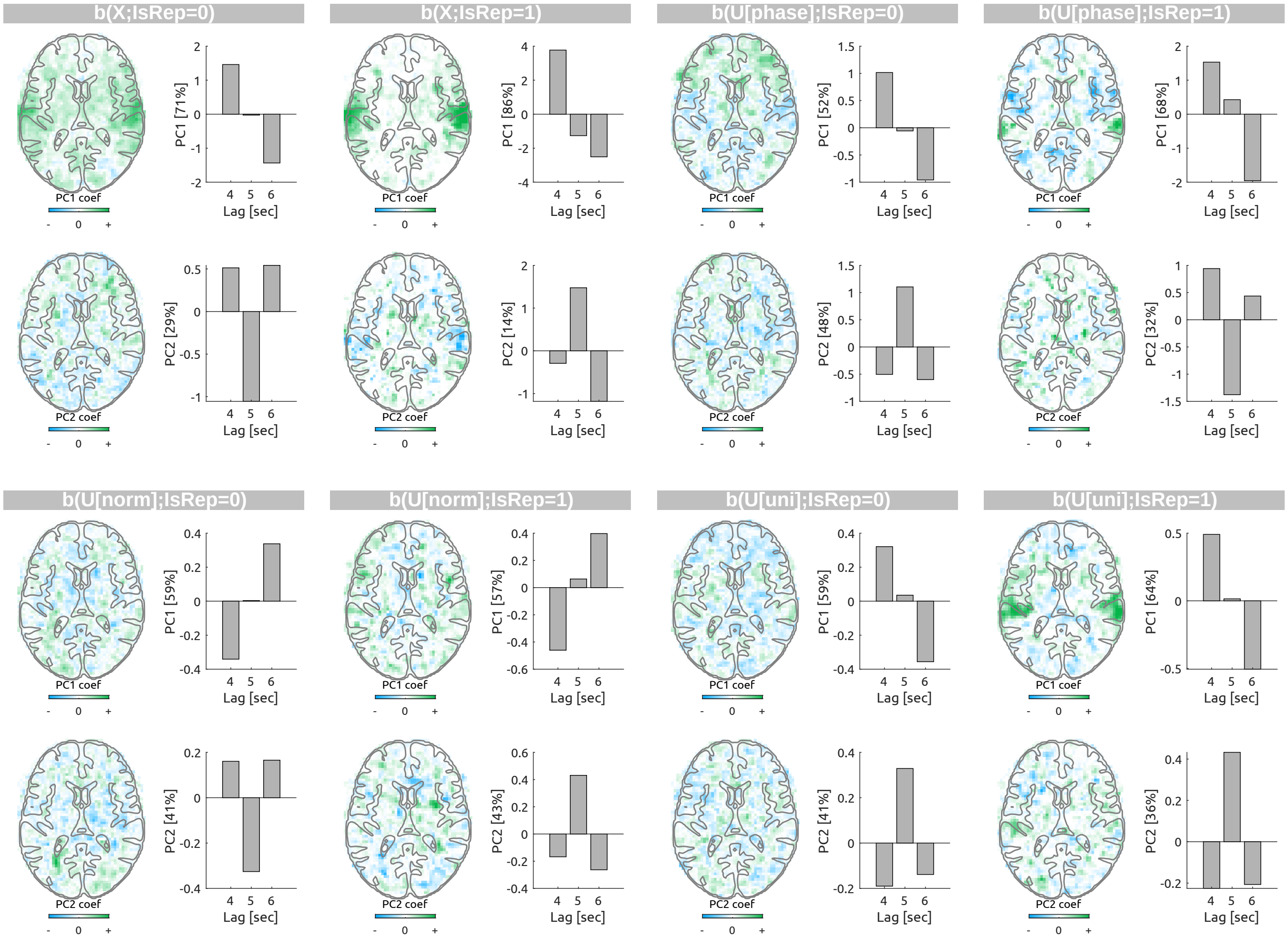


Figure S28: First two principal components (PCs) of the transfer function weights ($b$) of the models with delays from 4 to 6 sec are displayed. The visualisation scheme is identical to Figure S27.

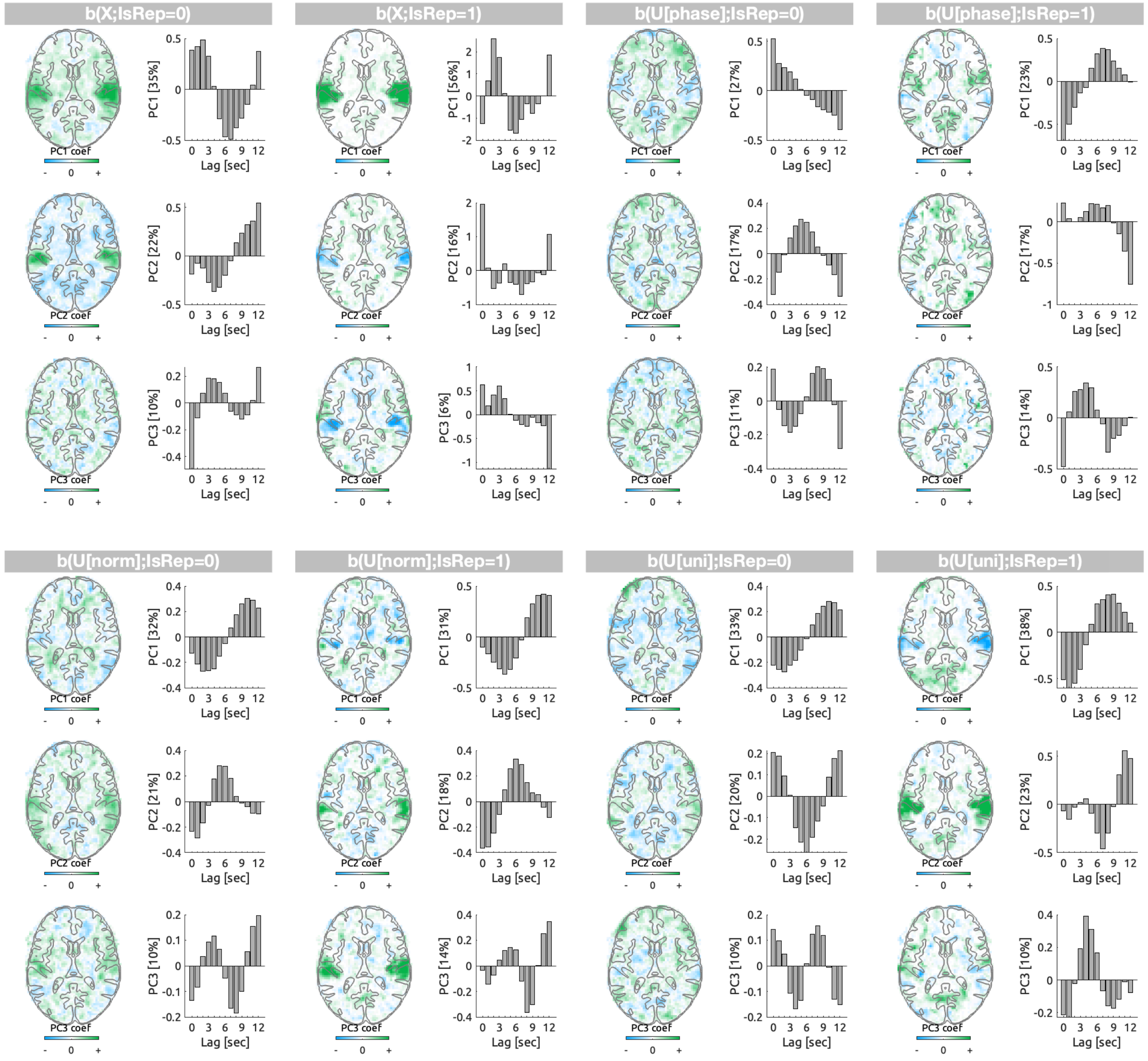


Figure S29: First three principal components (PCs) of the transfer function weights ($b$) of the models with delays from 0 to 12 sec are displayed. The visualisation scheme is identical to Figure S27.

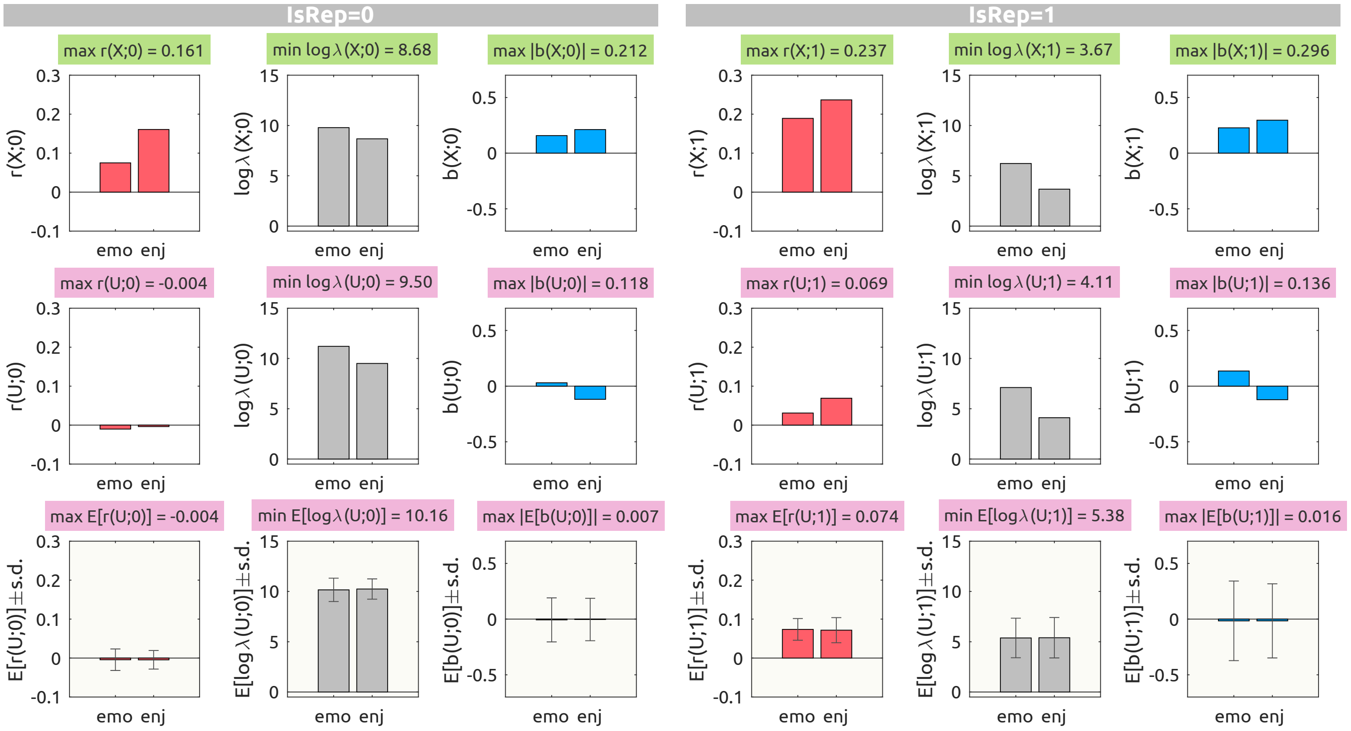


Figure S30: behavioural linearised encoding analysis results with delays from 0 to 10 sec with an audio envelope (top row), a single case of a phase-randomised envelope (middle row), and an average of 100 phase-randomised envelopes (bottom row, pale yellow background). For each CV scheme ($\texttt{IsRep} = 0$, left panels; $\texttt{Is Rep} = 1$, right panels), prediction accuracy ($r$, red bars), logarithmic ridge hyperparameter ($\log_{10} \lambda$, gray bars), transfer function weights that are summed over delays ($b$, blue bars) are shown along the columns.

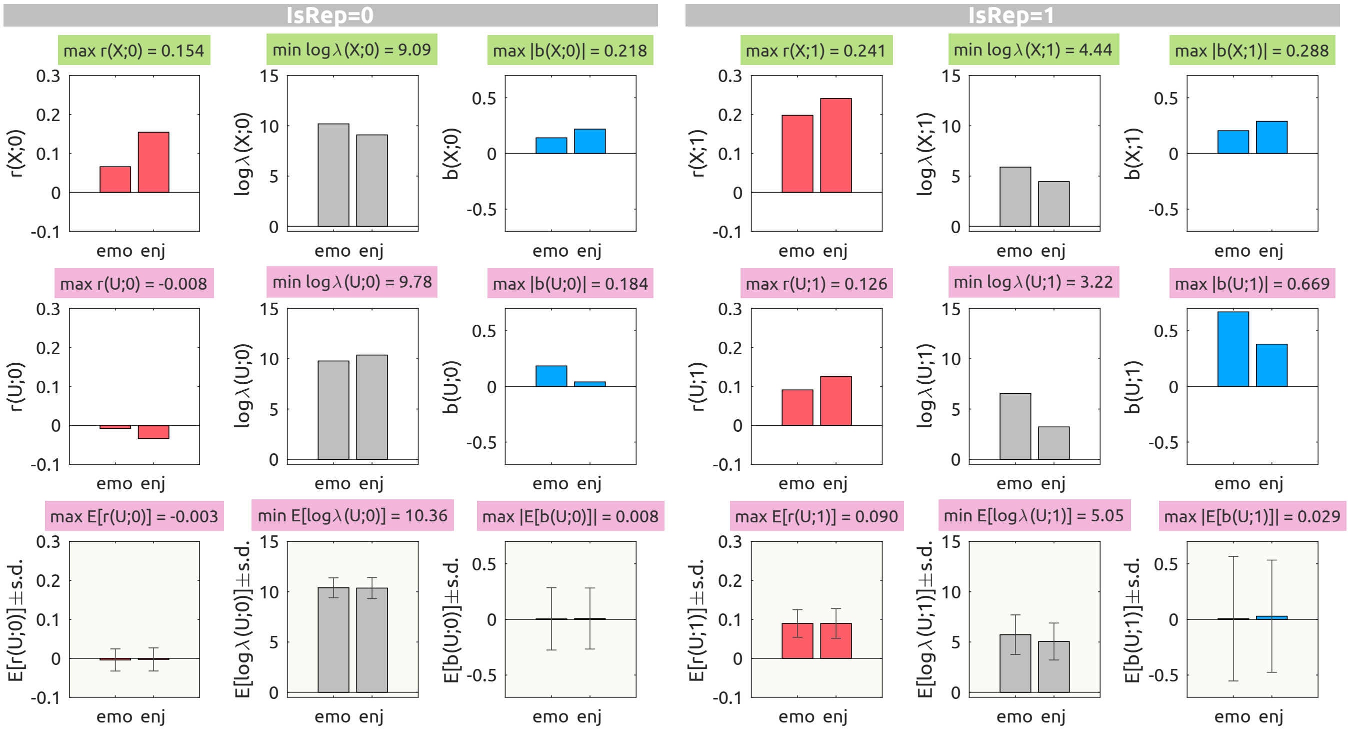


Figure S31: behavioural linearised encoding analysis results with delays from 0 to 10 sec with the uniform noise as the null feature. The visualisation scheme is identical to Figure S30.

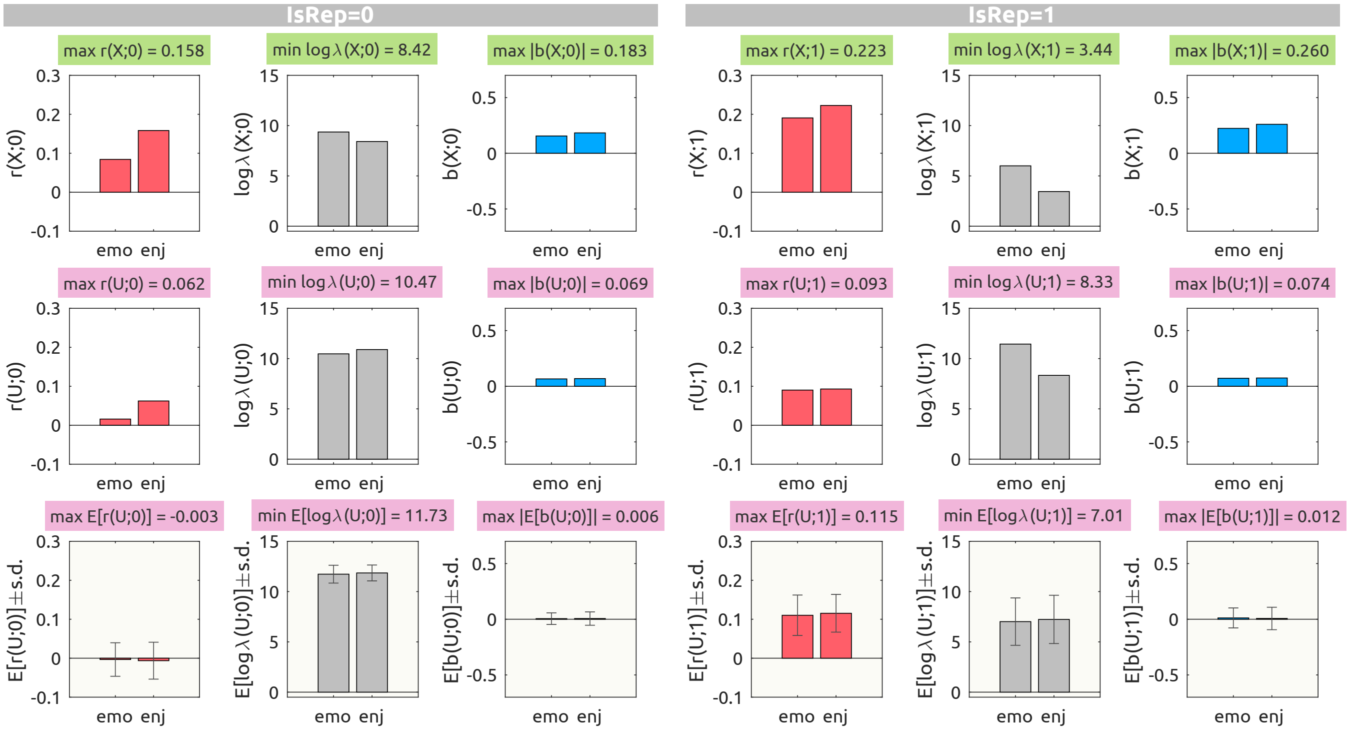


Figure S32: behavioural linearised encoding analysis results with delays from 0 to 5 sec with the phase-randomised envelope as the null feature. The visualisation scheme is identical to Figure S30.

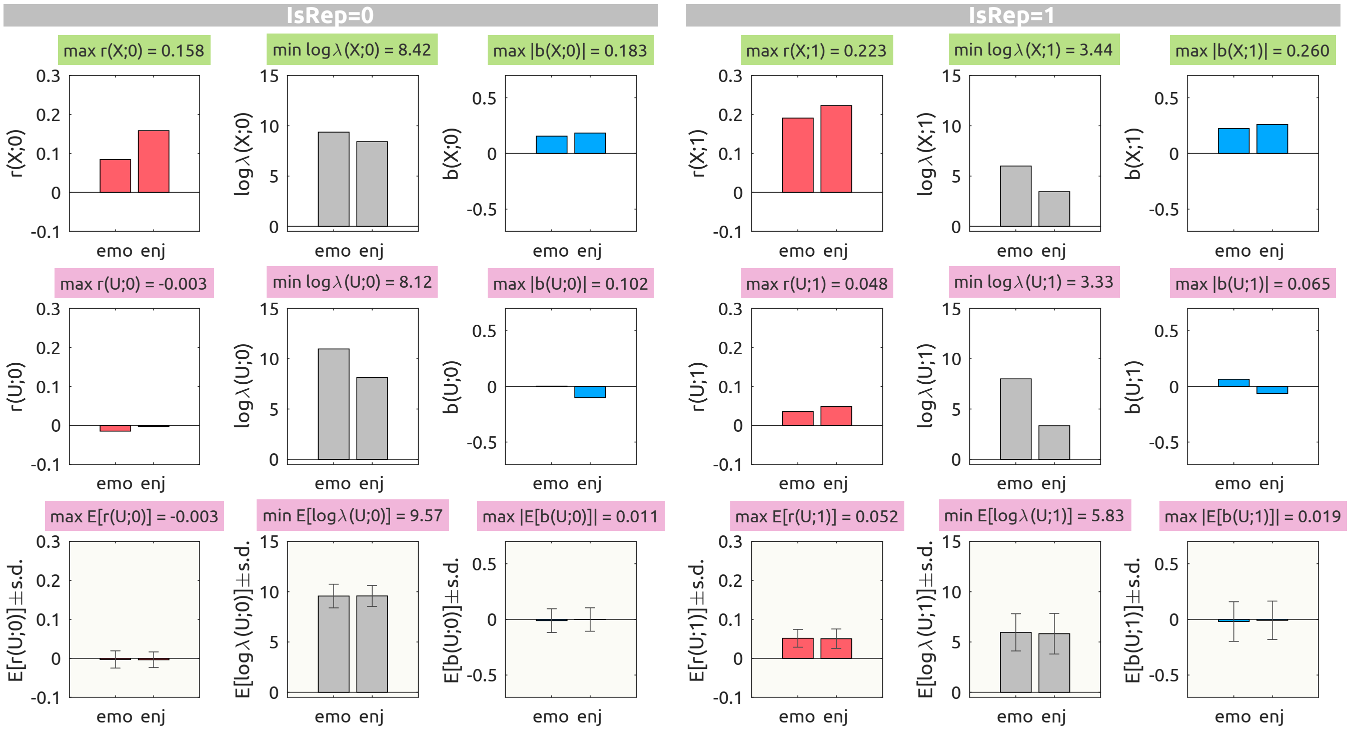


Figure S33: behavioural linearised encoding analysis results with delays from 0 to 5 sec with the phase-randomised envelope as the null feature. The visualisation scheme is identical to Figure S30.

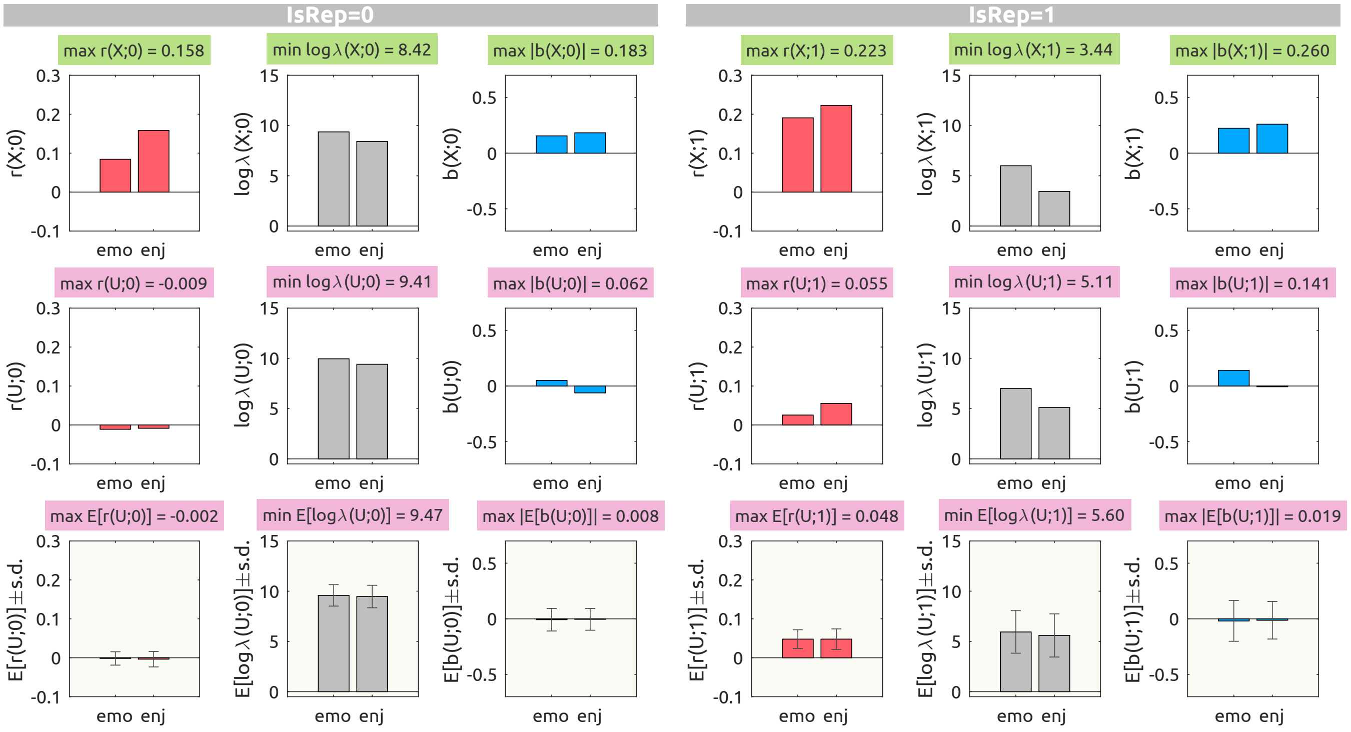


Figure S34: behavioural linearised encoding analysis results with delays from 0 to 5 sec with the uniform noise as the null feature. The visualisation scheme is identical to Figure S30.

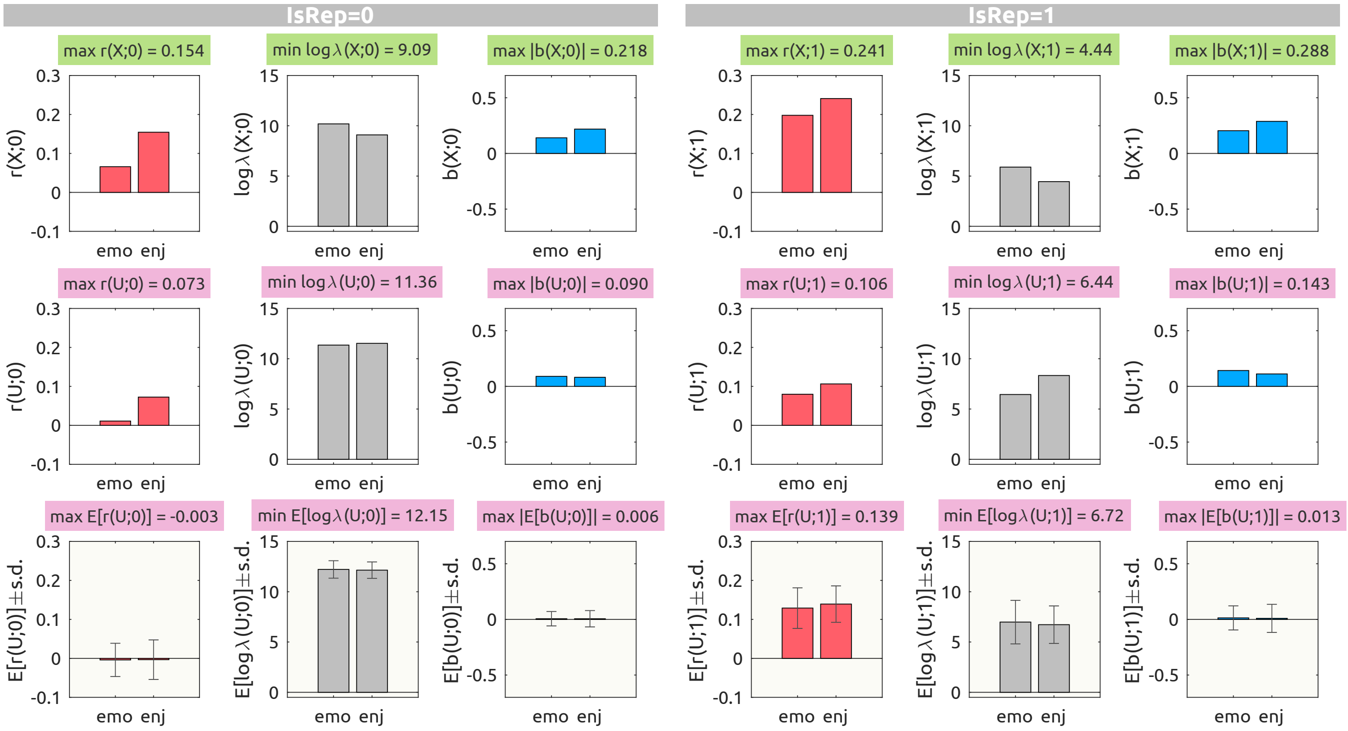


Figure S35: behavioural linearised encoding analysis results with delays from 0 to 15 sec with the phase-randomised envelope as the null feature. The visualisation scheme is identical to Figure S30.

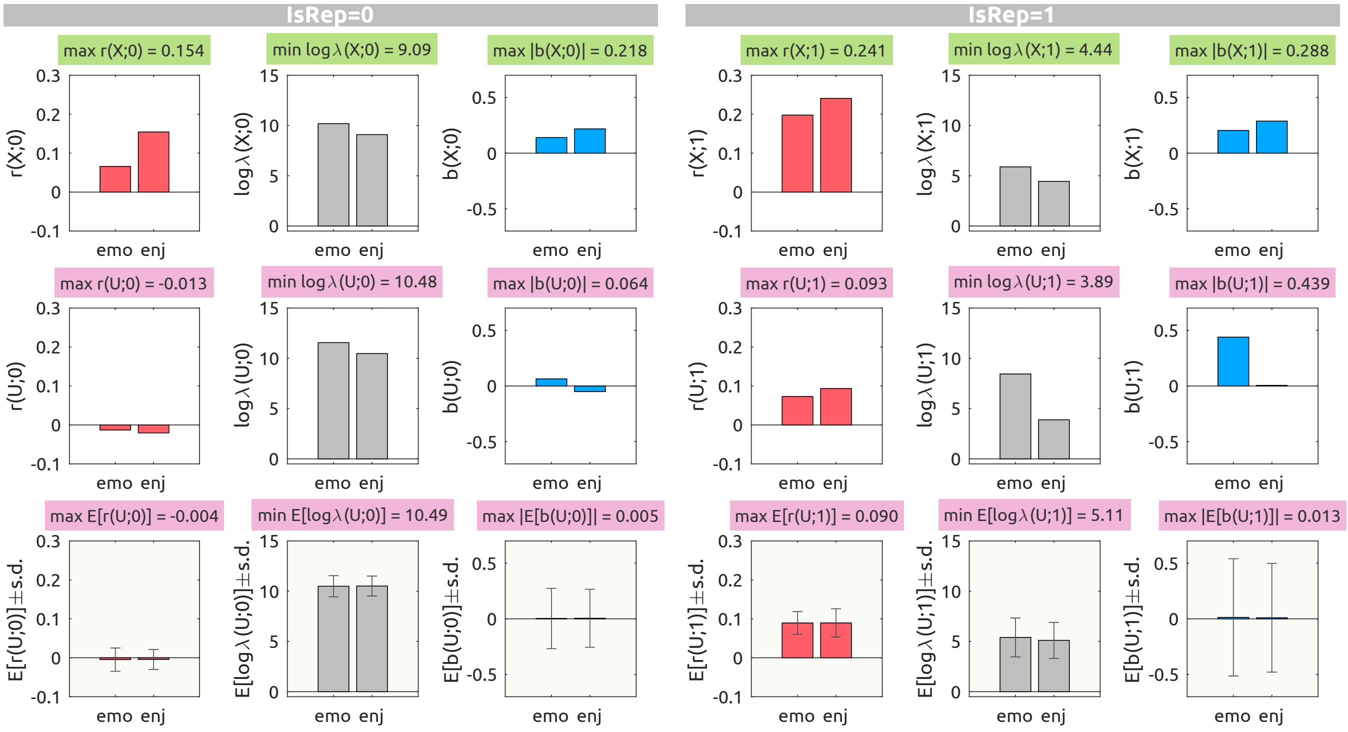


Figure S36: behavioural linearised encoding analysis results with delays from 0 to 15 sec with the phase-randomised envelope as the null feature. The visualisation scheme is identical to Figure S30.

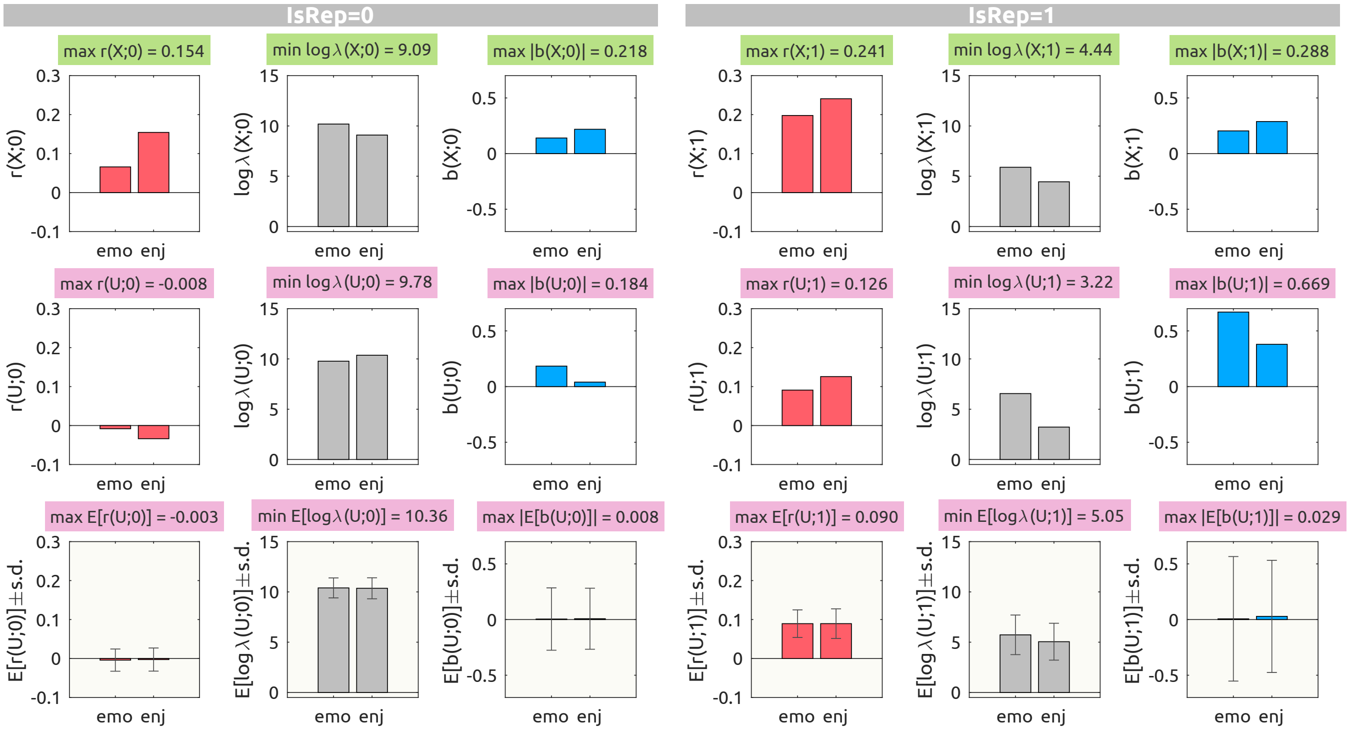


Figure S37: behavioural linearised encoding analysis results with delays from 0 to 15 sec with the uniform noise as the null feature. The visualisation scheme is identical to Figure S30.

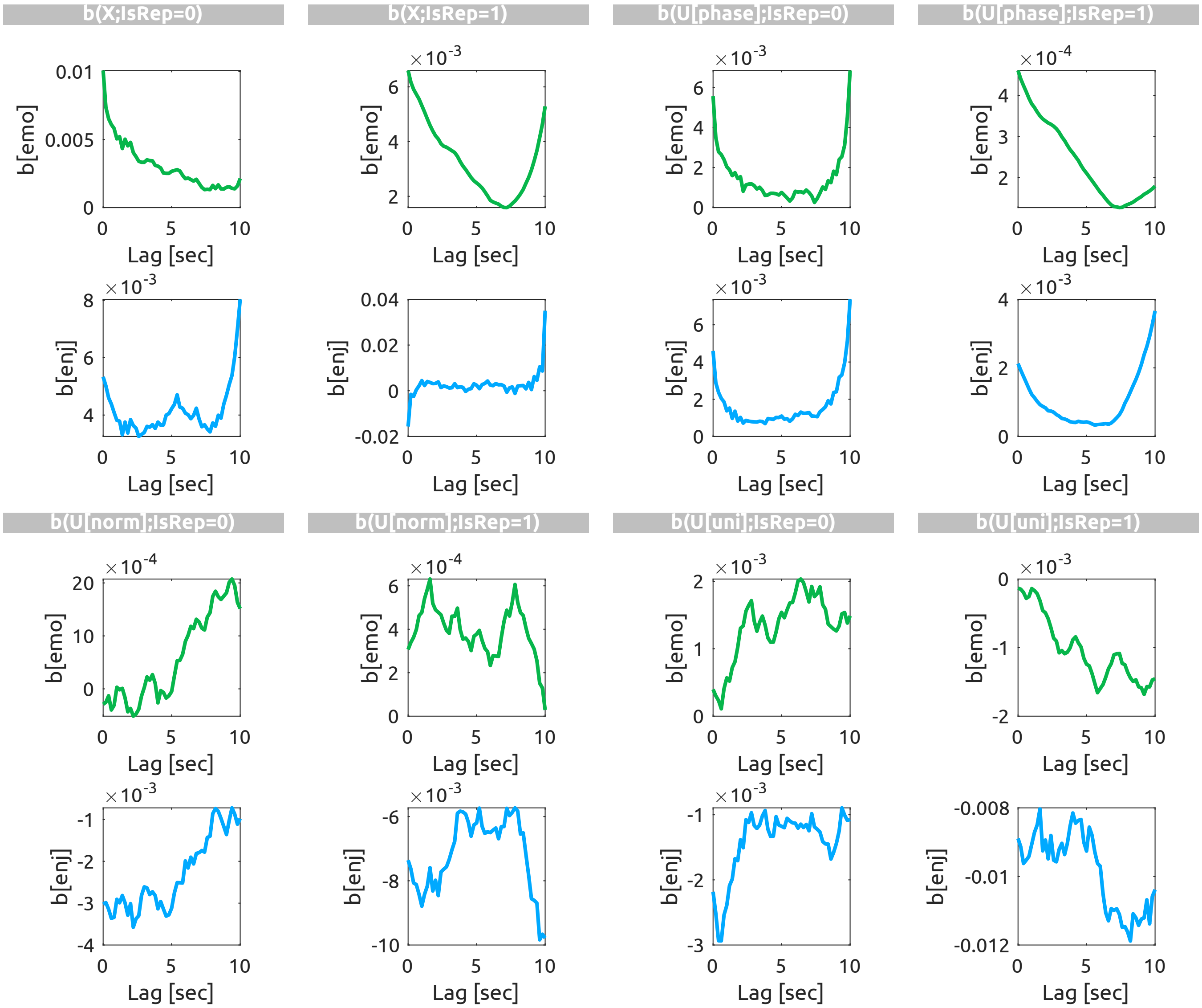


Figure S38: Transfer function weights ($b$) of Emotionality (green) and Enjoyment (blue) for the models with delays from 0 to 10 sec. The weights are grouped by features ($X$, true audio envelop; $U[phase]$, phase-randomised envelop; $U[norm]$, normal noise; $U[uni]$, uniform noise) and the CV schemes ($\texttt{IsRep} = 0$, $\texttt{IsRep} = 1$). Note that each weight time series is individually scaled.
/mul

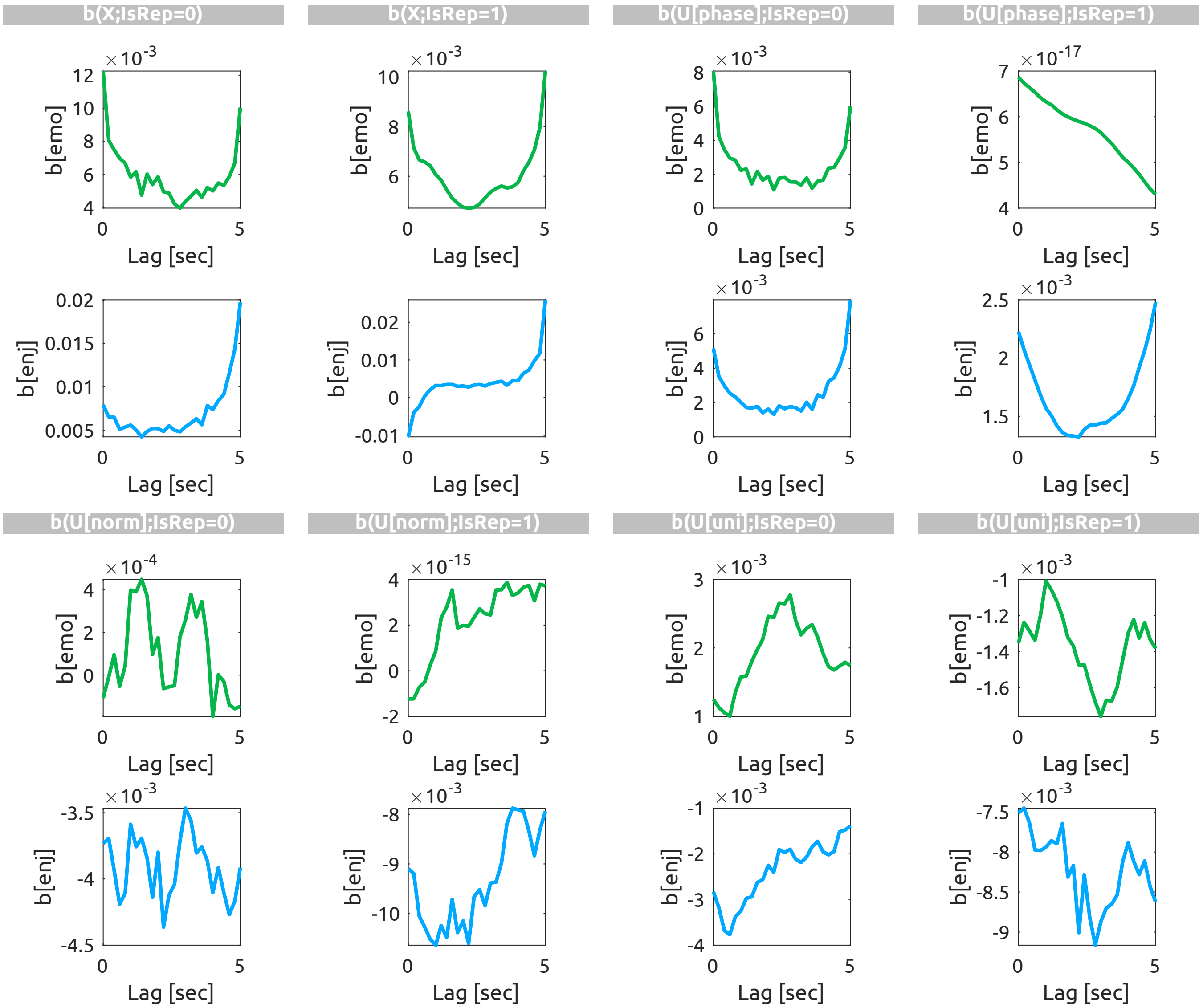


Figure S39: Transfer function weights ($b$) of Emotionality (green) and Enjoyment (blue) for the models with delays from 0 to 5 sec. The visualisation scheme is identical to Figure S38.

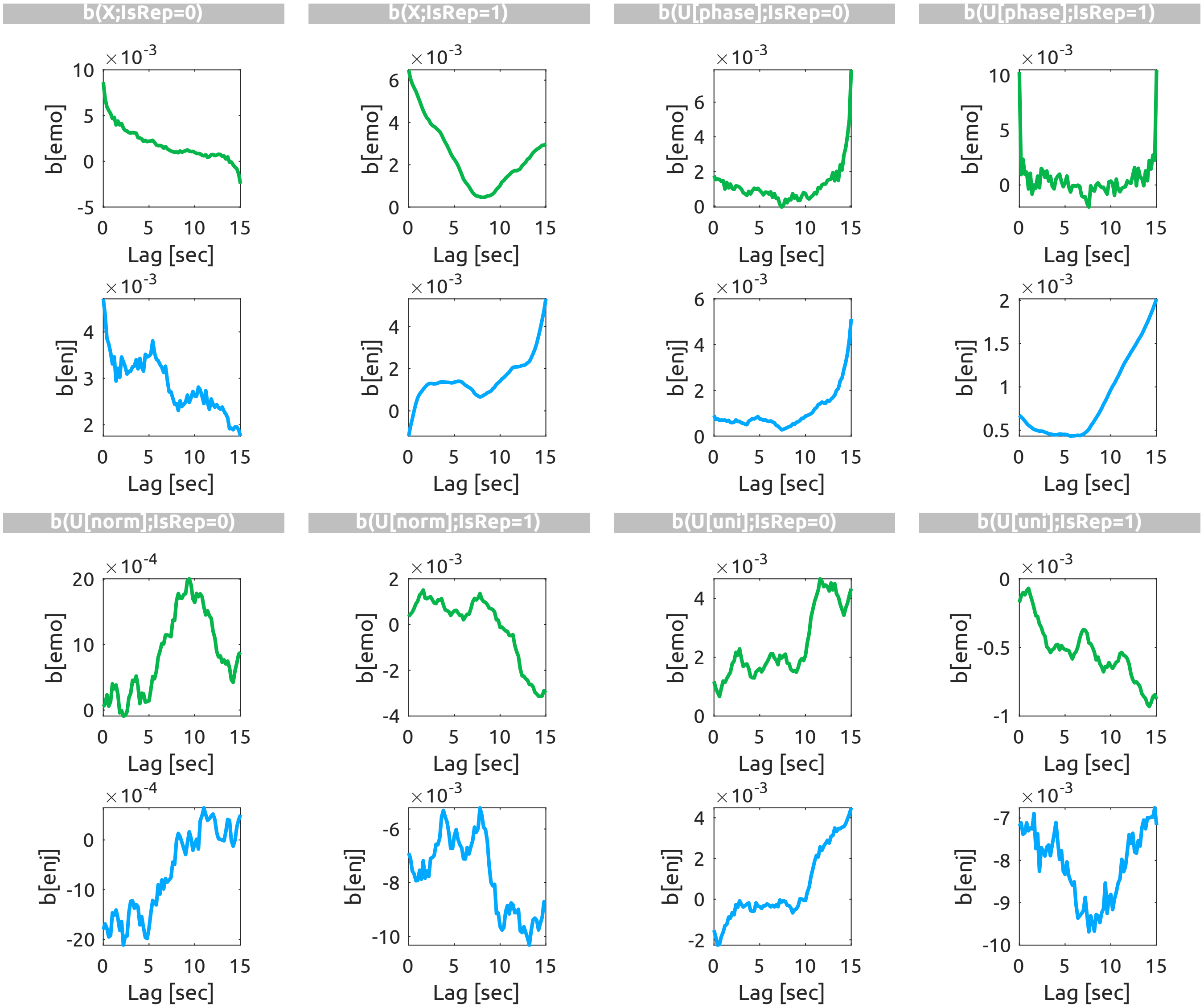


Figure S40: Transfer function weights ($b$) of Emotionality (green) and Enjoyment (blue) for the models with delays from 0 to 15 sec. The visualisation scheme is identical to Figure S38.
